# Starfysh reveals heterogeneous spatial dynamics in the breast tumor microenvironment

**DOI:** 10.1101/2022.11.21.517420

**Authors:** Siyu He, Yinuo Jin, Achille Nazaret, Lingting Shi, Xueer Chen, Sham Rampersaud, Bahawar S. Dhillon, Izabella Valdez, Lauren E Friend, Joy Linyue Fan, Cameron Y Park, Rachel Mintz, Yeh-Hsing Lao, David Carrera, Kaylee W Fang, Kaleem Mehdi, Madeline Rohde, José L. McFaline-Figueroa, David Blei, Kam W. Leong, Alexander Y Rudensky, George Plitas, Elham Azizi

## Abstract

Spatially-resolved gene expression profiling provides valuable insight into tissue organization and cell-cell crosstalk; however, spatial transcriptomics (ST) lacks single-cell resolution. Current ST analysis methods require single-cell RNA sequencing data as a reference for a rigorous interpretation of cell states and do not utilize associated histology images. Significant sample variation further complicates the integration of ST datasets, which is essential for identifying commonalities across tissues or altered cellular wiring in disease. Here, we present Starfysh, the first comprehensive computational toolbox for joint modeling of ST and histology data, dissection of refined cell states, and systematic integration of multiple ST datasets from complex tissues. Starfysh uses an auxiliary deep generative model that incorporates archetypal analysis and any known cell state markers to avoid the need for a single-cell-resolution reference in characterizing known or novel tissue-specific cell states. Additionally, Starfysh improves the characterization of spatial dynamics in complex tissues by leveraging histology images and enables the comparison of niches as spatial “hubs” across tissues. Integrative analysis of primary estrogen receptor-positive (ER^+^) breast cancer, triple-negative breast cancer (TNBC), and metaplastic breast cancer (MBC) tumors using Starfysh led to the identification of heterogeneous patient- and disease-specific hubs as well as a shared stromal hub with varying spatial orientation. Our results show the ability to delineate the spatial co-evolution of tumor and immune cell states and their crosstalk underlying intratumoral heterogeneity in TNBC and revealed metabolic reprogramming shaping immunosuppressive hubs in aggressive MBC. Starfysh is publicly available (https://github.com/azizilab/starfysh).

## Introduction

In multicellular systems, the function of diverse cell types is strongly influenced by their immediate surroundings. Uncovering the spatial organization and communication between cell types in tissues provides insight into their development, function, maintenance, response to stimuli, and how they adapt to their microenvironment or transform into malignant or diseased cells^1^. By sampling the entire transcriptome, recent advances in spatial transcriptomics (ST) enable unbiased gene expression mapping in a spatially-resolved manner, thus providing an unprecedented opportunity to study the spatial arrangement of cells and microenvironments^2^. These technologies have been employed in diverse fields, including organ development, disease modeling, and immunology^3, 4, 5^. However, sequencing-based methods (Visium, DBiTseq^6^, Slide-seq^7^, etc.) are limited in cellular resolution due to technical artifacts from lateral RNA diffusion^2^. Measurements from capture locations (spots) thus involve mixtures of multiple cells leading to analytical challenges in dissecting the cellular disposition, particularly in complex systems such as cancerous tissues.

Accurate characterization of cell types and refined cell states is particularly critical for comparing spatial organization and intercellular communications across tissues, e.g., for studying changes in cellular wiring and organization during development or disease progression. For example, in complex tumor tissues, the signal from patient-specific tumor cells is mixed with that from shared immune cell types, hindering the comparison of anti-tumor immune mechanisms between patients, disease subtypes, or diseases. The majority of existing computational methods for analyzing ST data (Cell2location^8^, DestVI^9^, Tangram^10^, Stereoscope^11^, RCTD^12^, BayesPrism^13^, etc.) require paired and annotated single-cell sequencing data as a reference to overcome this challenge and are not capable of integrating tissue samples. The references, whether from the same tissue or public databases, could introduce biases without accounting for sample or batch variation as well as variability in cell density across spots.

Importantly, access to paired single-cell data may not be cost-feasible or practical, e.g., in the case of clinical core biopsy samples, further motivating reference-free methods capable of integrating prior knowledge of cell type markers and data from multiple tissues to improve statistical power. Proposed reference-free methods such as STdeconvolve^14^, Smoother^15^, and CARD^16^ deconvolve spots into latent factors. However, some factors cannot be explicitly mapped to refined cell states in tumor tissues (**Supplementary Fig. 1)**. These methods also do not allow the integration of multiple ST datasets, and batch correction methods designed for single-cell RNA sequencing are not feasible in ST samples dominated by sample-specific cell types such as tumor cells. While some methods have utilized histology images to align spots between replicate tissues^8^ or to predict high-resolution gene expression from histology^17^ current methods fail to utilize spatial dependencies and paired histology for improving cell state deconvolution.

To address this major need, we developed a comprehensive toolbox for multi-modal analysis and integration of ST datasets dubbed Spatial Transcriptomic Analysis using Reference-Free auxiliarY deep generative modeling and Shared Histology (Starfysh). Through joint modeling of transcriptomic measurements and histology images, Starfysh infers the proportion of fine-grained and context-dependent cell states, as well as the spatial distribution of cell density, while obtaining cell type-specific gene expression profiles for downstream analysis. Integration of gene expression and histology accounts for tissue architecture, cell density, structured technical noise, and spatial dependencies between measurements, which improve the characterization of cell states and their arrangement. Importantly, by integrating multiple tissues, Starfysh can identify shared or sample-specific niches and their underlying cell-cell crosstalk.

The innovation of our machine learning approach is in incorporating archetypal analysis and any known cell type markers as priors in an auxiliary deep generative model that maps both transcriptomic and histology to a joint latent space. Archetypes, intuitively capturing spots with the most different expression profiles, are used to construct or refine cell type markers, as opposed to conventional clustering of spots which obtain markers corresponding to aggregated cell types^18^. This feature empowers Starfysh to characterize novel or context-specific cell states and present a hierarchy among them.

The performance of Starfysh on simulated data demonstrates successful, robust deconvolution without requiring single-cell references. Using Starfysh, we recapitulate the cell proportions of published datasets of breast tumors^19^. Additionally, we profiled tumor samples from ER^+^, TNBC, and MBC patients to demonstrate the utility and power of Starfysh for spatial mapping of inter- and intra-tumoral heterogeneity of human cancer and to investigate the role of microenvironmental niches in determining immune infiltration. Using the archetypal analysis feature of Starfysh, we successfully characterized patient-specific tumor cell states and their spatial arrangement within the primary tumor, revealing how the underlying biology of tumor states and environmental signals alter the immune response. We further identified metabolic reprogramming and communication enriched in the rare and aggressive MBC subtype by integrating our data with previously published ST datasets. Starfysh thus presents a powerful analytical platform for systematic interrogation and comparative studies of complex tissues in health and disease through the lens of spatial transcriptomics and histology.

## Results

### Starfysh performs reference-free deconvolution of cell states by combining spatial transcriptomics and histology

Starfysh is an end-to-end toolbox for multi-modal analysis and integration of ST datasets (**Fig. 1a**). In short, the Starfysh framework consists of reference-free deconvolution of cell types and fine-grained cell states, which can be improved with the integration of paired histology images of tissues, if available. To facilitate the comparison of tissues between healthy or disease contexts and deriving differential spatial patterns, Starfysh is capable of integrating data from multiple tissues and further identifies common or sample-specific spatial “*hubs”*, defined as niches with a unique composition of cell states. To uncover mechanisms underlying cell communication, Starfysh performs downstream analysis on the spatial organization of hubs and identifies key genes with spatially varying patterns as well as co-evolving cell states and co-localization networks.

To circumvent the need for a matched or external single-cell reference in deconvolving cell types, Starfysh leverages two key concepts to determine spots with the most distinct expression profiles as *“anchors”* that pull apart and decompose the remainder of spots (**Fig. 1b**). First, Starfysh incorporates a compendium of known cell state marker genesets as well as any additional custom markers of choice. Assuming that the spots with the highest overall expression of a geneset corresponding to a cell state are likely to have the highest proportion of that cell state, these spots form an initial set of anchors. Second, since cell state markers can be context-dependent or not well-characterized, Starfysh utilizes archetypal analysis to refine the initial anchor set and add any non-overlapping archetypes as additional anchors to enable the discovery of novel cell states or a hierarchy of cell states (**Methods, Supplementary Fig. 2**). This feature is paramount in characterizing context-specific cell states such as patient-specific tumor cell states, their phenotypic transformation and dynamic crosstalk within the microenvironment.

**Figure 1.**
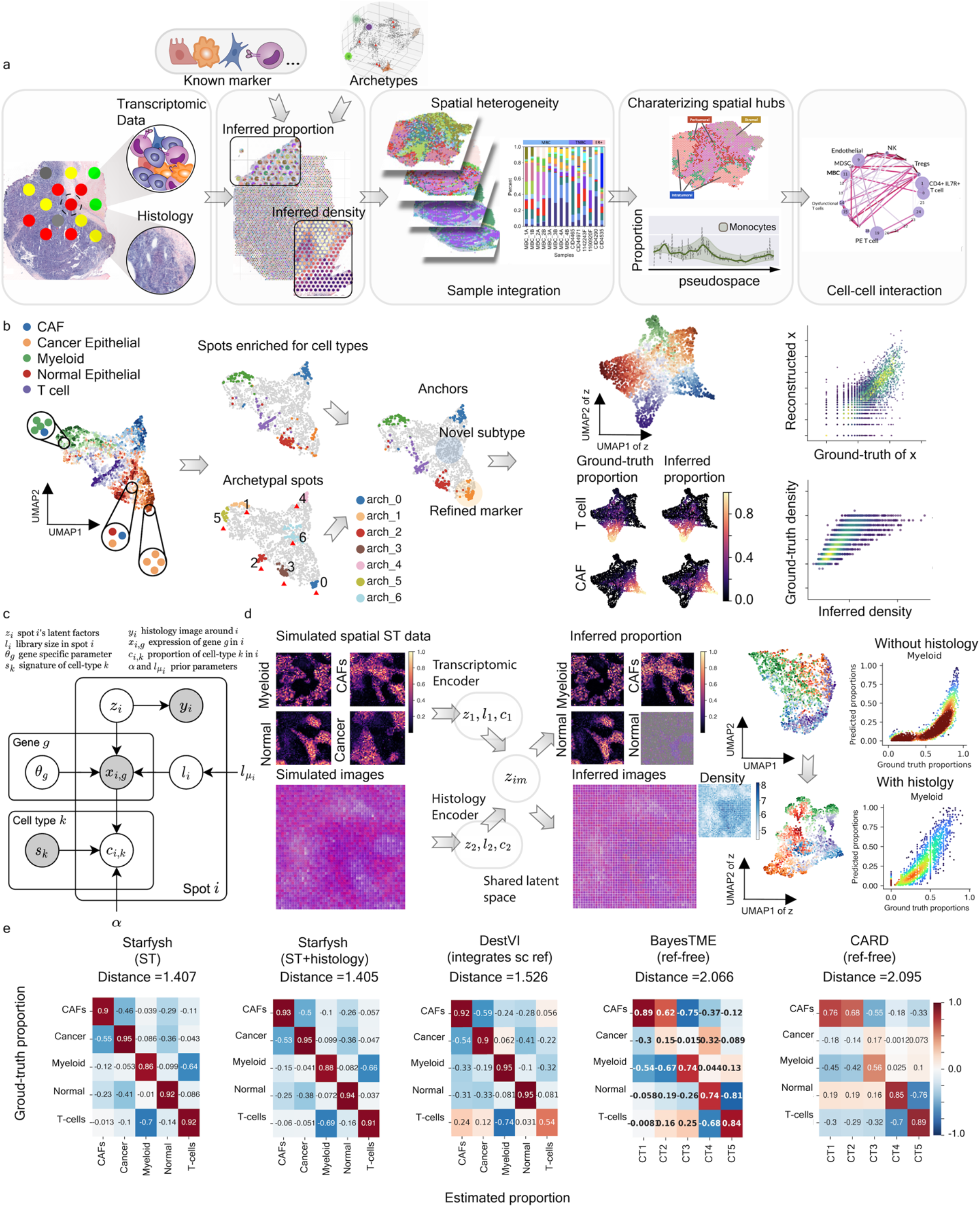
Starfysh overview and performance on simulated data. **(a)** Overview of the Starfysh workflow. From left to right: Starfysh input: spatial transcriptomics (ST) dataset, signature gene lists for cell types or cell states, and paired histology image (optional); Deconvolution: Starfysh defines anchor spots representative of cell types or states with the aid of archetypal analysis (**Supplementary Fig. 2**), and infers cell type/state proportions and densities by accounting for ST technical artifacts; Sample Integration and downstream analysis: Upon deconvolution, Starfysh jointly integrates multiple samples and characterizes spatial “hubs”, and further infers cell-cell interactions within each hub. **(b)** Left: UMAP of ST data with 2,500 spots, 29,631 genes, and 5 cell types simulated from mixtures of single-cell RNA-seq data of breast tumor tissues, colored by the proportion of most enriched cell type in the ground truth. Starfysh collectively utilizes signature gene sets and archetypal analysis to identify anchor spots, refine marker genesets, and discover potential novel cell states. Right: Comparison of ground truth cell-type proportions and densities in simulated data and Starfysh reconstruction (**Methods, Supplementary Fig. 4)**. **(c)** Graphical representation of the deep generative model integrating transcriptomic data and paired histology image to infer a joint latent space. **(d)** Spatial distribution of ground truth and inferred cell types with the same parameters in (b) and histology (**Methods)**. The integration of histology image improves the deconvolution performance in spatially-dependent simulated ST data **(Supplementary Fig. 8; Methods)**. **(e)** Benchmarking Starfysh against other methods on the data shown in (b,d): Pearson correlation of ground-truth and estimated proportions per cell type in data **(Supplementary Fig. 9; Methods)**. The performance of each method is summarized by computing the distance between the correlation matrix and an identity matrix (**Methods**). Benchmarking on real breast tumor ST data is shown in **Supplementary Fig. 1**.

Inspired by successful implementations of deep generative models in single-cell omics analysis (scvi-tools^20^, scvi^21^, totalVI^22^, scArches^23^, trVAE^24^, scANVI^25^, MrVI^26^), Starfysh jointly models ST and histology as data observed from a shared low-dimensional latent representation while incorporating anchors as priors (**Fig. 1c; Supplementary Fig. 3a, b;** see **Methods**). The innovation in our variational inference is in using the cell-state proportions as auxiliary variables^27^, and in integrating the transcriptomes with the histology (see **Methods; Supplementary Fig. 3c**).

To test the performance of Starfysh, we simulated ST data from real single-cell RNA-sequencing data from primary breast tumor tissues^19^ with and without spatial dependencies (**Supplementary Fig. 4; Methods**). We found the successful recovery of cell types proportions (R2 score=0.85, Pearson correlation = 0.93 with p<1e-30) and cell density (Pearson correlation = 0.90 with p<1e-30) (**Supplementary Fig. 5, 6; Fig. 1b**).

Starfysh is capable of integrating histology to correct for artifacts in transcriptomic measurements by accounting for spatial dependencies between spots and incorporating tissue structure which improves the estimation of cell density and in turn, characterization of cell types and neighborhoods, in particular in complex tissues. The integration of two data modalities is accomplished using the product of experts (PoE^28^), which calculates the joint posterior distribution for gene expression and images (**Fig. 1d;** see **Methods**). We simulated ST data with spatial dependencies using a Gaussian Process model^8^ (see **Methods**) and simulated images by training a ResNet18^29^ encoder followed by a decoder to paired ST expression and histology images (**Supplementary Fig. 7;** see **Methods**). Simulated ST data harbored cell clumps and histology patterns resembling real tissues (**Supplementary Fig. 4**). PoE integrates the latent factors from transcriptomic and histology data, and shows significantly improved performance in predicting the proportion of cell types (e.g., myeloid cells) and reconstructs high-density regions (**Fig. 1d,e; Supplementary Fig. 8)**. The deconvolution performance of Starfysh is comparable to existing state-of-the-art methods that require a single-cell reference including DestVI^9^, Cell2Location^8^, Tangram^10^ and BayesPrism^13^ (**Fig. 1e; Supplementary Fig. 9**). Additionally, compared to reference-free methods such as CARD^16^, BayesTME^30^, Starfysh shows a significant improvement in deconvolving major cell types (**Fig. 1e**; Mann Whitney U test p=2.6e-5; see **Methods**). Importantly, Starfysh shows substantial improvement in disentangling fine-grained cell states (Mann Whitney U test p<1e-30) and runtime compared to other methods (**Supplementary Fig. 1**). These results confirmed the ability to extract signal corresponding to relatively smaller cells such as tumor-infiltrating immune cells and constructing hierarchies of cell types and states. Such distinctions are not possible with other methods and are crucial in understanding heterogeneous immune responses in healthy and pathological tissues^31^.

### Starfysh dissects the spatial heterogeneity of primary breast tumors

To demonstrate the capability of Starfysh to characterize heterogeneous and complex tissues, we sought to examine the spatial dynamics of immune response in primary breast adenocarcinomas. Heterogeneity in the immune cell composition of individual tumors has been connected to the variability of patient responses to cancer treatments, including immunotherapies^32, 33, 34^. We previously showed that the tissue of residence is a significant determinant of the diversity of immune phenotypic states, and that T and myeloid lineage cells exhibit a continuous phenotypic expansion in the tumor compared to matched normal breast tissues^35^. Heterogeneous T cell states were highlighted by combinatorial expression of genes reflecting responses to various microenvironmental stimuli while being tightly associated with T cell receptor (TCR) utilization^35^. These data thus suggested that TCR specificities may contribute to the spatial organization of T cells through the disposition of cognate antigens, facilitating their exposure to niches differing in the extent of inflammation, hypoxia, expression of activating ligands and inhibitory receptors, and nutrient supply.

To investigate this hypothesis, we performed ST profiling of 8 primary tumors from an ER^+^, a classic TNBC, and two metaplastic TNBC breast cancer (MBC) patients (2 biological replicates for each) (**Supplementary Table 1**; **Methods**). The resulting data, alongside published datasets^19^ from 6 ER^+^ and TNBC breast cancer patients (1 biological replicate for each patient) were analyzed using Starfysh.

We first dissected the spatial heterogeneity in a classic TNBC tumor and characterized 29 diverse cell states, including normal epithelial, cancer epithelial, immune cells (naive CD4^+^ T cell, effector memory CD4^+^ T cell, myeloid-derived suppressor cells - MDSC, macrophages, CD8^+^ T cells) and stromal cells (endothelial, PVL immature, PVL) (**Fig. 2a,b**; **Methods, Supplementary Table 2, Supplementary Fig. 10, 11)**. Importantly, due to the heterogeneity of tumor cells^36^, Starfysh defined patient-specific tumor cell states by aligning spots enriched for known tumor cell genesets with archetypal spots that capture extreme phenotypic states, resulting in a refined anchor set that guided the deconvolution of spots (**Fig. 2a-d**). The process of identifying example anchors, including regulatory T cells (Tregs), and two tumor cell states, showed an improved separation of cell states after updating genesets according to aligned archetypes (**Fig. 2a-d**, see **Methods**). Additionally, the estimated cell density and reconstructed image were consistent with the histology (maximal information coefficient = 0.33; compared to 0.18 for shuffled pixels in histology) (**Fig. 2e;** see **Methods**).

**Figure 2.**
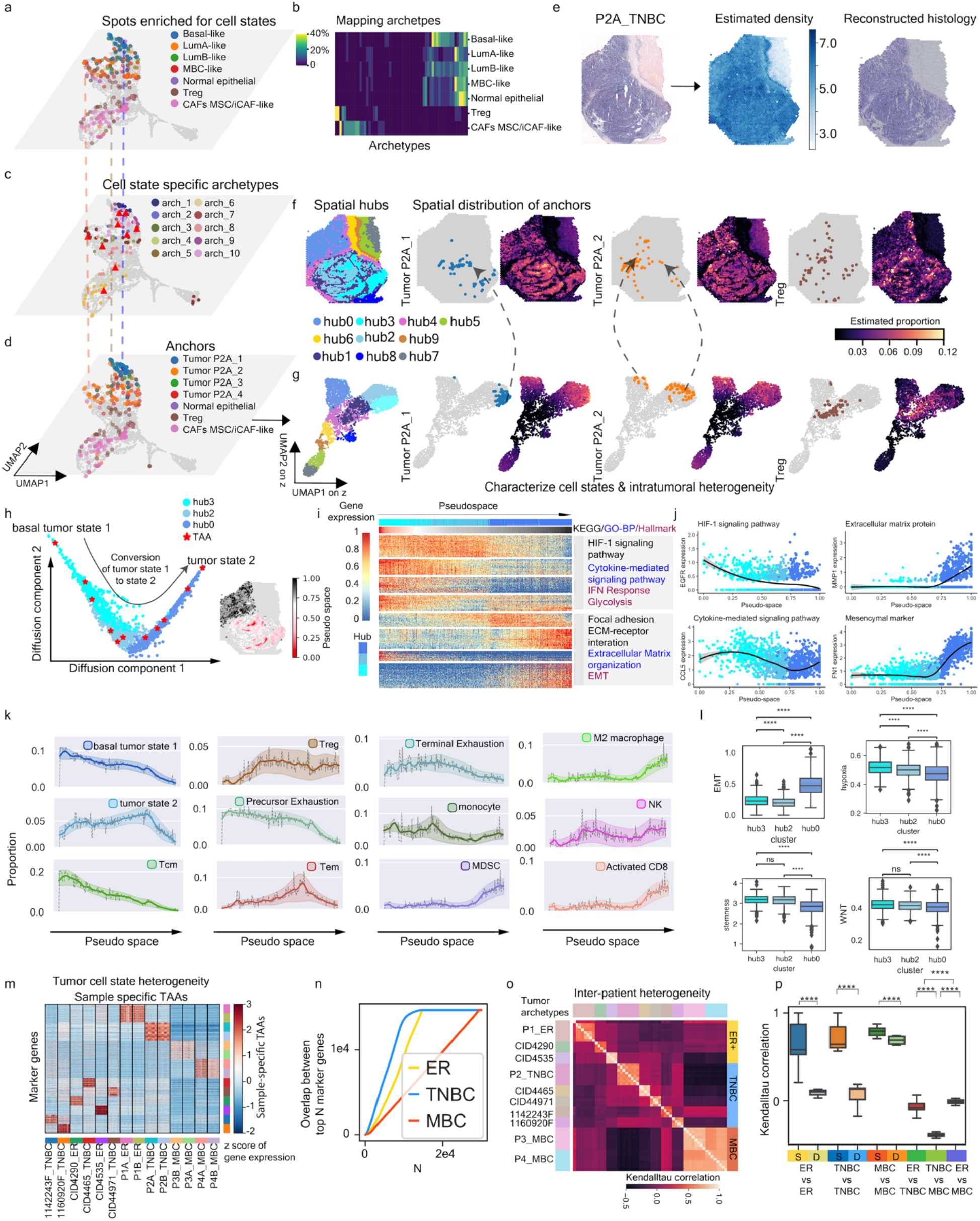
Characterizing spatial tumor heterogeneity in breast carcinoma. **(a)** UMAP 2D projection of ST gene expression data from a TNBC patient (sample P2A), with each dot representing a spot (grey). Spots enriched for four example cell types are highlighted in color. **(b)** Mapping all archetypes to the cell types shown in (a). The mapping score is defined by the percentage of overlap between neighborhoods of archetypal spots and anchor spots (**Methods**). (c) Archetypal spots associated with cell types in (a) with mapping score >=35% (**Methods, Supplementary Fig. 10**). **(d)** Spots enriched for cell types are combined with archetypes to achieve a refined anchor set e.g. identifying patient-specific tumor states. **(e)** Histology for P2A_TNBC, and reconstructed histology and tissue density using Starfysh. **(f-g)** Spatial hubs, distribution of anchors, and inferred proportions, for two tumor cell states and Tregs, in the spatial context of the tissue (f) and UMAP projection of inferred latent factors from Starfysh (g). **(h)** Diffusion map analysis revealing a continuous trajectory between spots enriched for tumor cell states. Tumor-associated anchors (TAAs) are shown with red stars. The dominant trajectory was inferred with Scorpius^46^ and shown in the tissue context representing a pseudo-space axis. **(i)** Top row: Tumor states identified with Starfysh. Second row: Pseudo-space for spots sorted along the trajectory inferred in (h). Bottom: Heatmaps of expression of modules of genes with positive or negative correlation with the projection of cells along the trajectory and select pathways enriched with gene set enrichment analysis (GSEA). **(j)** Expression of marker genes in pathways shown in (i) in spots projected on the trajectory. **(k)** Changes in the proportion of tumor and immune cell states across the pseudo-space axis inferred in (h). **(l)** Expression of genesets demonstrating enrichment in any intratumoral hub. **(m)** Heatmap of expression of top 20 gene markers (rows) differentially expressed in TAAs (columns) grouped by sample. **(n)** Overlap between top N marker genes differentially expressed in TAAs in any pair of patients. **(o-p)** Kendall’s Tau correlation between rankings of genes according to differential expression scores in TAAs (o), and grouped by patient subtype (p) (S: correlations among sample patients; D: correlations among different patients).

To understand the association between the tumor cell phenotypes and the TME, we defined spatial “*hubs*” as groups of spots with similar composition by applying Phenograph^37^ to the inferred composition of spots (**Fig. 2f; Supplementary Fig. 12**). This analysis revealed that heterogeneous tumor cell states reside in different spatial hubs with more basal-like tumor cells enriched in hub 3 while a second tumor cell state expressing a subset of luminal A markers (e.g., *MTDH, SMARCB1, FOXA1*) are present in hub 0 (**Fig. 2f; Supplementary Fig. 12)**, and the two states correspond to two terminal branches in the inferred latent space (**Fig. 2g**). This analysis also uncovered regions with varying composition of infiltrating immune cell types exemplified by hub 1 and hub 4 composed of Treg-enriched spots (**Fig. 2f, g; Supplementary Fig. 10, 11**). Altogether, these findings showed the capability of Starfysh in elucidating intratumoral transcriptional heterogeneity and characterizing diverse and patient-specific tumor cell states, in part determined by their spatial context in the tissue and co-localization with immune subsets.

### Starfysh delineates a co-evolving tumor-immune transition in TNBC

Further analysis of spots enriched for tumor cells using diffusion maps^38, 39^ (**Supplementary Fig.13;** see **Methods**) revealed a surprisingly continuous transition between the two tumor cell states which also corresponds to a spatial gradient (**Fig. 2h**). The inferred pseudo-space axis is associated with upregulation of extracellular matrix organization and ECM-receptor interaction pathways and loss of IFN and cytokine-mediated signaling related gene expression (**Fig. 2i, j**). Furthermore, the tissue-repair (M2) macrophage signature was also increased along the transition. These tumor-associated macrophages (TAMs) have been implicated in promoting the invasion, migration, and proliferation of TNBC cells^40^. This observation, indicative of locally induced epithelial-mesenchymal transition and up-regulation of collagen genes, associated with metastatic potential^41, 42, 43^, reproduced in the adjacent tissue sample (**Supplementary Fig. 14**), suggests that intratumoral heterogeneity is a continuum rather than two extreme states of tumor cells. Indeed, projecting all anchors enriched for tumor genesets as “*tumor-associated anchors*” (TAAs), showed they are uniformly distributed along the pseudo-space axis (**Fig. 2h**), thus representing different stages of this transformation.

We then sought to investigate whether different immune cell states are associated with regions with varying tumor phenotypes. Remarkably, we found a compositional shift from central memory and precursor exhausted T cell states^44^ to effector memory, terminally exhausted and Treg states, as co-localized tumor cells lose basal properties along the pseudo-space axis spanning intratumoral hubs, while activated T cells are observed at the tumor margins (**Fig. 2k, Supplementary Fig. 13**). Interestingly, the tumor state transformation axis coincides with a loss of stemness, down-regulation of WNT signaling pathway, and changes in metabolic states from hypoxia and glycolysis to glucose deprivation and tissue repair (**Fig. 2l, Supplementary Fig. 15**). This observation indeed suggests that different T cell states are associated with various niches of the TME shaped by varying nutrient supply, oncogenic signals and tumor cell differentiation states (**Supplementary Fig. 16**). For example, WNT, NOTCH, HIPPO, PI3K, NRF2, TP32, RAS pathways are depleted, while TGFβ is enriched along this tumor transformation. In parallel, we see a shift from monocytes to MDSCs and TAMs, thus showing the ability of Starfysh to spatially resolve tumor immune response and identify co-evolving tumor-immune phenotypic changes that underlie tumor heterogeneity.

### Starfysh enables quantification and comparison of tumor heterogeneity across tissues

The patient-specific characterization of tumor states also enables the quantification of intra- and inter-tumor heterogeneity. Specifically, we used differential gene expression analysis to identify markers characterizing tumor associated anchors (TAAs) in all breast tumor samples as well as published samples^19^. We found that marker genesets corresponding to tumor states in biological replicates originating from the same patient tumor are significantly overlapping as expected, while distinct modules of non-overlapping markers illustrate intra-patient heterogeneity (**Fig. 2m**). Quantifying the overlap in top marker genes associated with tumor states in each sample, exhibited greater divergence in markers representing MBC tumor states, implicating higher inter-tumoral heterogeneity among MBC samples compared to TNBC and ER samples (**Fig. 2n**), in line with the known morphological heterogeneity of MBCs^45^. This result was further supported by comparing the ranking of differentially expressed genes, where we found a lower correlation between TAA gene rankings in MBC patients compared to TNBCs (**Fig. 2o,p**).

### Starfysh characterizes spatial hubs from the integration of breast tumor samples

To demonstrate the potential of Starfysh in deriving commonalities among heterogeneous samples and disease subtypes, we performed an integrated analysis of 37,517 spots from all 14 samples of 10 patients (**Supplementary Table 3;** see **Methods**). UMAP dimensionality reduction of ST data without Starfysh revealed no overlap among patients, partly due to patient-to-patient variation, given that replicate samples overlap (**Fig 3a**). Moreover, the aggregation of patient-specific tumor cells with other cell types within spots hindered the comparison of shared immune states and spatial neighborhoods between patients. While batch correction methods designed for single-cell data failed in correcting the variations between patients (**Supplementary Fig.17**), Starfysh successfully integrated all datasets (**Fig 3b**). It yielded more mixing of shared immune states quantified with the entropy of the local distribution of patients (see **Methods**), yet preserved differences between patient-specific tumor cells (**Fig. 3c, d**). Overall, this analysis showed that luminal B tumors displayed lesser heterogeneity compared to basal and luminal A tumors.

**Figure 3.**
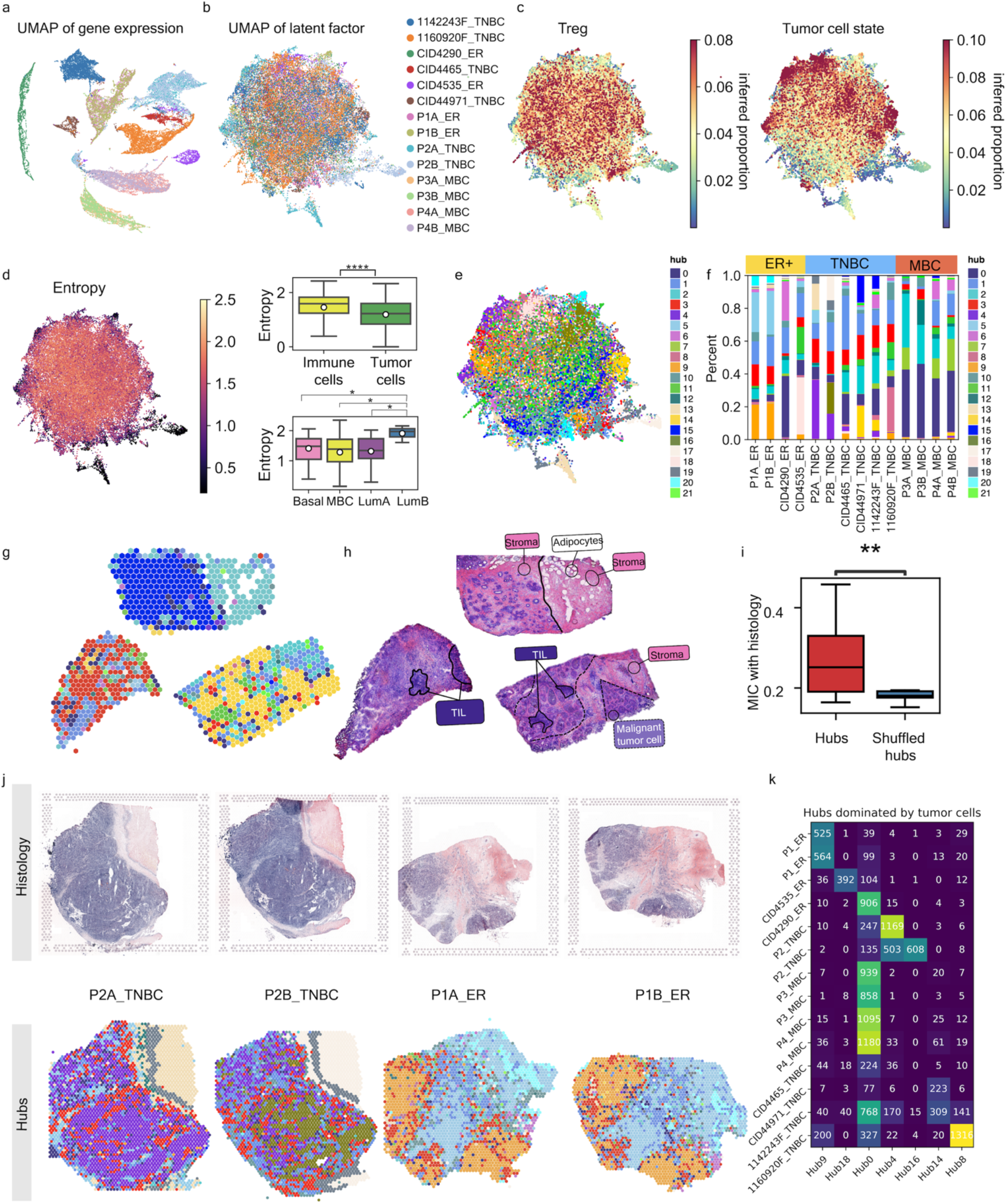
Characterizing tumor-immune hubs from the integration of samples. **(a)-(b)** UMAP visualization of gene expressions of 4 MBC samples, 6 TNBC samples, and 4 ER+ samples before **(a)** and after **(b)** Starfysh integration. **(c)** UMAP visualization of Starfysh inferred proportions from integration of spots from all samples colored by the proportions of a tumor cell state and an example immune cell state (Treg) in the integrated space. **(d)** UMAP of integrated space colored by Shannon’s entropy in each spot and boxplots of entropy grouping spots by disease subtype. **(e)** UMAP of integrated space colored by hubs identified by clustering spots based on inferred cell-type proportions.. **(f)** Spatial hub distribution for each sample **(g)-(h)** Spatial arrangement of hubs (g) and pathological histology annotation of sample 44971_TNBC (h). Inferred hubs align well with annotated ductal carcinoma in situ (DCIS) (red hub), lymphocyte infiltrated (olive green hub), and stroma (yellow hub) regions. **(i)** Maximum information coefficient (MIC) for alignment of hubs with histology. **(j)** Paired histology and spatial arrangement of hubs for a TNBC and ER patient showing consistencies between replicates of the same patients and with histology. **(k)** Number of spots assigned to intratumoral hubs in each patient.

To understand the similarities and differences in the organization of cell states among patients and samples, we identified spatial hubs by merging and clustering spots from all samples according to inferred cell state proportions (**Fig. 3e**). The majority of hubs were detected in more than one patient (**Fig. 3f**). The distribution of hubs, however, varied between disease subtypes and patients. The spatial arrangement of hubs showed a marked similarity to expert-annotated histology, including in rare normal epithelium regions, tumor infiltrated regions, and immune cells enriched regions (**Fig. 3g,h**), which was quantified using maximum information coefficient (MIC) (**Fig. 3i**, see **Methods)**. As expected, hub distributions had similar patterns between biological replicates, i.e., adjacent sections of tumor tissues (e.g., P1A_ER and P1B_ER), whereas hubs dominated by tumor cells were different between patients (e.g., P1 and P2) (**Fig. 3j, k**).

### Metabolic remodeling shapes an immunosuppressive niche in metaplastic breast tumors

Through the integration of ST datasets, we performed a systematic comparison of tumor heterogeneity and its interplay with tumor-immune characteristics across subtypes of breast cancer. Specifically, we investigated potential differences in the cellular organization in MBC tumors compared to other TNBCs that might explain their worse clinical outcomes ^47^. MBC is a rare and aggressive subset of breast cancers, making for 1-2% of all breast cancer^41^. MBC is typically characterized as TNBC because these tumors largely lack the expression of estrogen receptor (ER), progesterone receptor (PR), and human epidermal growth factor 2 receptor (HER2). However, MBCs have a worse prognosis compared to conventional TNBC and exhibit greater resistance to chemotherapy^41, 48, 49^. A hallmark of MBC is its significant morphological heterogeneity reflected in its name^45, 50^. This distinguishing feature alongside enrichment in macrophages and immunosuppressive Tregs^51^ motivates the spatial characterization of tumor-immune crosstalk in the MBC TME to help guide the development of novel therapeutic approaches tailored to MBC’s unique biology.

We partitioned hubs according to their spatial arrangement around tumor regions (**Fig. 4a, Supplementary Fig.18, 19**) into intratumoral, peritumoral, and stromal hubs, representing hubs with enriched tumor cells, regions adjacent to intratumoral hubs, and distant regions, respectively (**Fig. 4a, Supplementary Fig. 20)**. Notably, tumor samples did not show significant overlaps in intratumoral hubs, supporting heterogeneity of tumor cells across patients (**Fig. 3k, Fig. 4b**). To understand phenotypic differences in MBC tumor states, we projected TAAs in the inferred joint space from the integration of all samples provided by Starfysh (**Methods**). Using diffusion map analysis, we found a tumor state transition trajectory from a TNBC-enriched state to an MBC-specific state (**Fig. 4c, Supplementary Fig. 21**) which correlated with the regulation of tumor growth and reduction of glycolytic process signature. Importantly, MBC-specific states were associated with inflammatory response, hypoxia, EMT, and tumor necrosis (**Fig. 4d, Supplementary Fig. 22**). The expression of EMT and hypoxia-related gene sets, and the distribution of samples along the trajectory also confirmed the enrichment of EMT and hypoxia in MBC intratumoral hubs (**Fig. 4e, f**). Oncogenic pathways such as PI3K/AKT and p53 were also enriched in MBC intratumoral hubs, while G1/S and G2/M pathways were down-regulated (**Supplementary Fig. 23**). Interestingly, the spatial distribution of EMT and hypoxia indicated a gradual diminution in their prominence in peritumoral and stromal regions in MBC (**Fig. 4g)**, while TNBC and ER did not show this trend (**Supplementary Fig. 24**).

**Figure 4.**
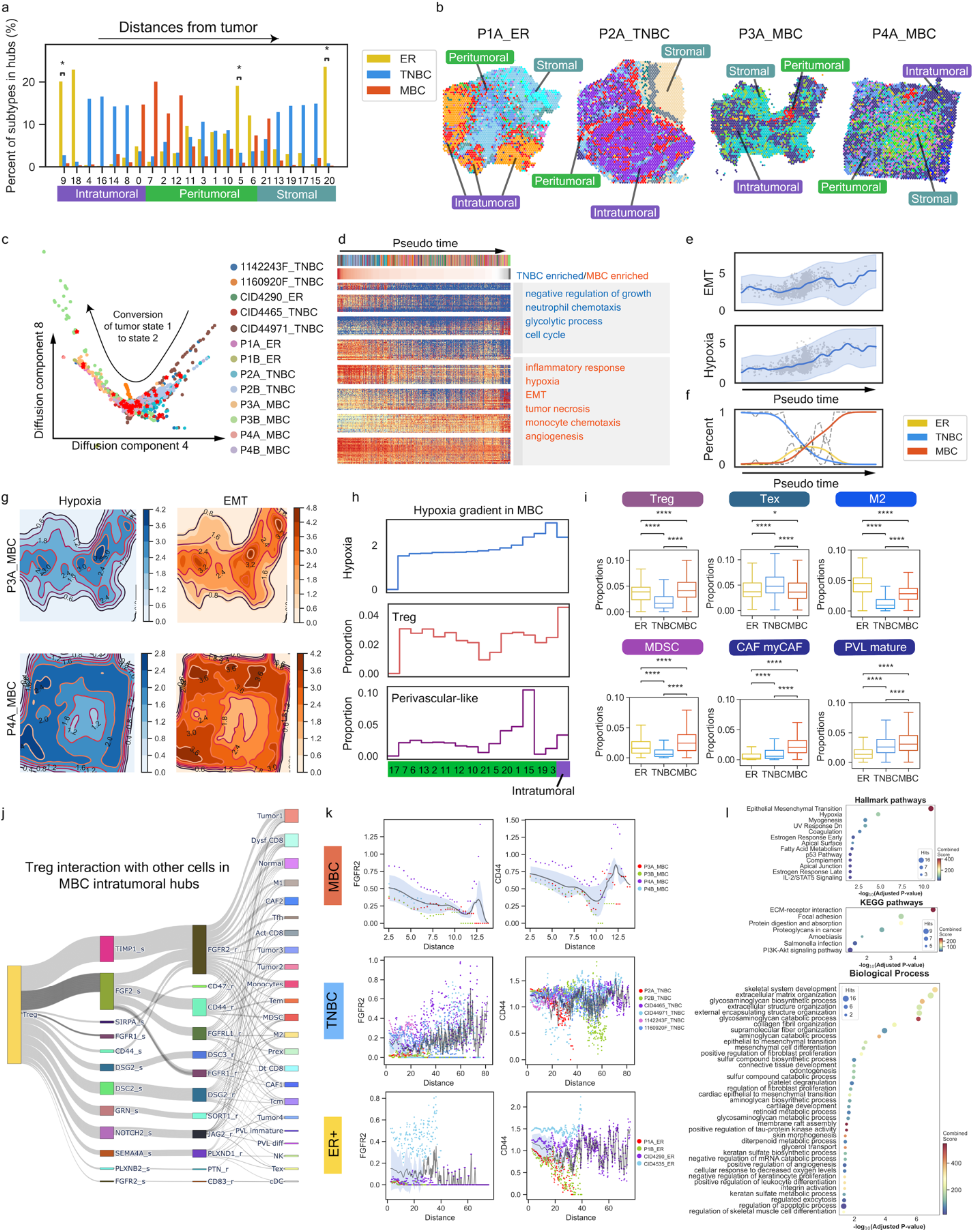
Intratumoral inflammation and heterogeneity in MBC epithelia. **(a)** Classification of spatial hubs according to distance from the tumor and matched histology. Percentage of spots from subtypes in each hub. **(b)** The spatial arrangement of hubs. **(c)** Diffusion map analysis reveals a continuous trajectory between TAAs across different patient samples. Archetypes are shown with red stars to represent the most distinct states for TAAs. The dominant transition trajectory was inferred with Scorpius^46^. **(d)** Top row: Spots ordered by inferred pseudo-time using Scorpius based on diffusion component in (c). Second row: Pseudo-time for spots sorted along the trajectory inferred in (c). Bottom: Heatmaps of expression of modules of genes with positive or negative correlation with the projection of cells along the trajectory and select pathways enriched with gene set enrichment analysis (GSEA). **(e)** The expression of EMT and hypoxia-relevant genesets show highly correlated dynamics along pseudo-time. **(f)** Percentage of ER+, TNBC, and MBC spots along the inferred pseudo-time. **(g)** Contour map showing the expression gradients of EMT- and hypoxia-related gene sets. Upper: sample P3A_MBC; Lower: sample P4A_MBC. **(h)** Expression of hypoxia geneset and predicted proportions of Tregs and PVLs (binned and sorted by stromal (left) to intratumoral (right) hubs. **(i)** Comparison of inferred intratumoral cell-state proportions across tumor subtypes. **(j)** Predicted significant receptor-ligand interactions between Tregs (ligand) and other cell types (receptor) in MBC intratumoral regions. **(k)** Expression of FGF2 and CD44 averaged across spots in each tumor subtype after binning according to distance from Treg-enriched hub 3. **(l)** Enrichment analysis for MBC intratumoral hubs. Significant pathways with FDR<0.05 are shown.

In agreement with increased hypoxia in the intratumoral hub, we observe an enrichment of Tregs and perivascular-like cells in MBC (**Fig. 4h**). In fact, the enrichment of Tregs in intratumoral hubs was only detected in MBC, implicating Treg-infiltration as a potential hallmark of this subtype of breast cancer. Besides Tregs, other immunosuppressive cells such as M2-like macrophages, MDSC, and cancer-associated fibroblasts (CAFs), in addition, perivascular-like cells were uniquely enriched in MBC intratumoral hubs compared to TNBC and ER (**Fig. 4i; Supplementary Fig. 25**). Previous literature has shown hypoxia affecting EMT in cancer by regulating EMT signaling pathways, EMT-associated miRNA, and lncRNA networks^52^. Both hypoxia and EMT were reported to modulate the TME by recruiting immunosuppressive cell types such as Tregs^53,54^. In line with this observation, a negative correlation was observed between CD8+ tumor-infiltrating lymphocytes and EMT markers in MBC tumors (**Fig. 4i, Supplementary Fig. 25)**. Hypoxia is also known to confer therapy resistance by inducing cell cycle arrest, inhibiting apoptosis, and mitochondrial activity^55^. Therefore, a subpopulation of tumor cells that survive hypoxia may be those responsible for resistance to chemo- and radiotherapy.

To understand communication patterns utilized by MBC tumor-infiltrating Tregs, we predicted receptor-ligand interactions potentially supporting crosstalk between Tregs and other cell states enriched in intratumoral hubs, using CellPhoneDB^56^ (**Fig. 4j, Supplementary Fig. 26, 27;** see **Methods**), revealing immunosuppressive pathways such as *FGF2, TIMP1, FGFR1, CD44*. Notably, FGF2 is a pro-tumor angiogenesis factor and induces drug resistance in chemotherapy in breast cancer^57^. TIMP1 is also found to be correlated with cancer progression and significantly increased in tumor-infiltrating lymphocytes (TILs) and Tregs ^58 59^. FGFR1 induces the recruitment of macrophages and MDSCs in the tumor via CX3CL1^60^, while CD44 is a known marker of breast cancer stem-like cells and stabilizes Treg persistence and function^61^. Interestingly, the diffused expression of these receptors with distance from Treg-enriched spots was only observed in MBC (**Fig. 4k**) further supporting their involvement in intratumoral Treg communication. Moreover, co-localization of cell states quantified using the spatial correlation index (SCI^62^, **Methods, Supplementary Fig. 26, 27**) indicates the potential involvement of Tregs in triads with tumor cells and terminally-differentiated exhausted T cells. These results demonstrate complex crosstalk in response to the immunosuppressive signals generated by Tregs.

Gene enrichment analysis in MBC intratumoral hubs also revealed EMT, hypoxia, p53 pathway, IL-2/STAT5 signaling, ECM, and PI3K-AKT signaling in MBC samples (**Fig. 4l**). Notably, the genomic landscape of MBCs shows frequent mutations in TP53 and genes related to the PI3K/AKT/mTOR pathway^63, 64^. Our data thus suggests possible coordination of nutrient uptake including glucose through HIF1 and PI3K/Akt pathways^65^ supporting enhanced growth and proliferation^66^ in intratumoral MBC hubs^66^ while this metabolic reprogramming is associated with immunosuppressive crosstalk (**Supplementary Fig. 28**). The diffused expression of HLA genes with distance from Tregs suggests the selection of adaptive Tregs could be allowed by increased MHCII driven by EMT (**Supplementary Fig. 28**).

### Spatial organization and interactions in the stromal breast TME

To dissect the stromal TME responding to the unique microenvironment niches, such as gradients of EMT and hypoxia in MBC, we characterized the cellular composition of peritumoral regions including hubs 1, 2, 3, 7, 10, 12 (**Supplementary Fig. 20, 25**). Intriguingly, Treg-enriched hub 3 (**Fig. 4b**, red), was present in all samples but showed unique patterns in each disease subtype. For example, it enveloped tumor hubs in the ER sample while being spatially scattered in the TNBC tumor (**Fig. 4b; Supplementary Fig. 29**). In contrast, in MBC, hub 3 was concentrated at certain locations close to intratumoral hubs, and the proportion of Tregs was lower than in intratumoral hubs, confirming Treg infiltration into MBC tumor regions.

Activated CD8^+^ T cells were enriched in peritumoral hub 3 in MBC samples compared to intratumoral regions, in contrast to their enrichment in intratumoral hubs in TNBC and ER^+^ (**Fig. 5a**). In addition to the spatial shifts of T cell states, endothelial cells were also enriched in hub3 in MBC, indicative of heightened angiogenesis in the stromal TME of MBC (**Fig. 5a**), which was particularly apparent in the histology of the region (**Fig. 5b, 4g**) likely as an adaptation to hypoxia. Predicted Treg interactions in hub 3 show expression of *CCL5, CXCR3*, and *CXCL11, CXCL2* in TNBC (**Fig. 5c**), compared to the predicted expression of neuropilin 1 and 2 NRP1/2 uniquely in MBC (**Supplementary Fig. 30**) which has been associated with stimulation of angiogenesis, consistent with the identification of interaction of VEGFA in our samples, enriched immunosuppressive roles, and worse prognosis of MBC patients^67^. Cytotoxic CD8^+^ T cells failing to infiltrate MBC tumors were associated with a unique ligand-receptor predicted link between SPP1 mainly secreted by tumor epithelia, CD8^+^ T cell deletional tolerance^68^, cDC, and MDSC cells, and *CD44* expression on cytotoxic CD8^+^ T cells. SPP1/OPN interacting with CD44 has been identified as a process for T cell suppression^69^, associated with epithelial cell migration, activation of PI3K/AKT signaling, and a hallmark of lower patient survival and chemo-resistance^69,70,71^. GAS6/AXL interaction^72^, uniquely shown in MBC, is also known to promote tumor cell proliferation, survival, migration, and angiogenesis^73^ (**Fig. 5d**).

**Figure 5.**
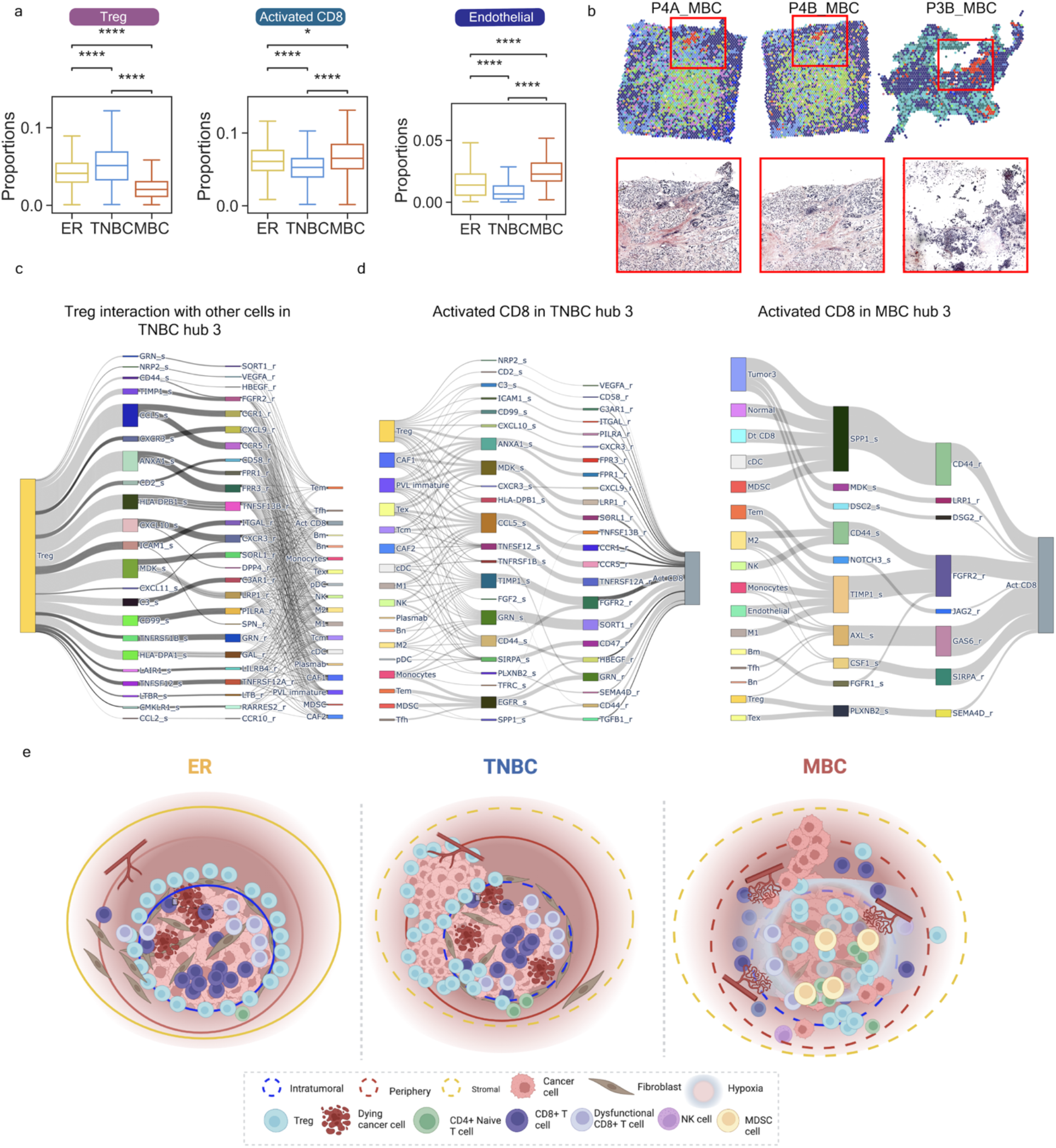
Spatial heterogeneity of the stromal breast TME. **(a)** Treg, activated CD8 T cell, and endothelial cell states in peritumoral hub 3 in ER, TNBC, and MBC samples. **(b)** The spatial arrangement of hubs and corresponding histology indicate blood cells and vessels around hub 3 in MBC. **(c)** Predicted interactions with Tregs in TNBC stromal hub 3. **(d)** Interactions involving activated CD8 T cells in hub 3 in TNBC and MBC tumors **(e)** Summary diagram.

Overall, Starfysh enabled the characterization of the spatial TME in MBC differing from TNBC and ER which is also summarized in **Fig. 5e**. We illustrate the enriched tumor suppressive cells possibly blocking CD8+ T cell infiltration, and underlying enriched hypoxia, EMT potential, and angiogenesis in the MBC TME.

## Discussion

By incorporating archetypal analysis and prior knowledge of markers of cell states in a deep generative model, Starfysh dissects the spatial heterogeneity of complex tissues from the combination of spatial transcriptomic and histology imaging data, without the need for a single-cell reference. Deconvolution of refined cell states is accomplished through the introduction of auxiliary variables in the inference scheme and is improved with the integration of histology, providing information on the tissue architecture, cell density, and spatial dependencies between transcriptomic measurements. Importantly, Starfysh can integrate multiple heterogeneous tissue samples, disentangle fine-grained cell states, and identify shared or tissue-specific cell states and spatial hubs to understand their role in tissue function and disease pathogenesis. These key features make Starfysh an ideal tool to discover and query spatial hubs from integrated large-scale datasets, increasing our power to detect features of complex and rare diseases that could drive future therapeutic strategies.

We applied Starfysh to investigate the role of spatial breast tumor heterogeneity in defining the continuous phenotypic expansion of tumor-infiltrating immune cells^35^. We showed the ability to characterize patient-specific heterogeneous tumor cell states and identified a tumor cell state transition within a complex TNBC tumor. Remarkably, the tumor cell transformation from basal states to mesenchymal states dovetails with a shift in the distribution of immune cell states. Specifically, a T cell activation axis coincided with moving farther away from tumor cells that exhibit stemness features and reside in hypoxic regions, supporting the hypothesis that the spatial orientation of tumor cell subsets determines immune differentiation.

We demonstrate the power of Starfysh in integrating multiple complex tissue samples using our newly generated data alongside previously published ST datasets, to quantify intra- and inter-tumoral heterogeneity, and identify spatial hubs as regions with a similar composition of cell states. Importantly, integrating samples enabled the comparison of rare chemo-resistant metaplastic breast tumors to other breast cancer subtypes. Notably, we found intratumoral infiltration of Tregs, M2-like macrophages, and MDSCs in MBC shaping an immunosuppressive niche enriched in EMT and hypoxia (**Fig. 5e**). Crosstalk with immunosuppressive Treg and MDSCs was predicted to be mediated through FGF2, TIMP1, FGFR1, CD44 signaling pathways which would be top candidates for future functional studies. Indeed, FGFR signaling is known to maintain EMT-mediated drug-resistant populations^74^. Enrichment of p53 and PI3K-AKT pathways in MBCs also suggests reprogramming of metabolic activity in MBC tumors. Our data thus motivates further investigation of FGFR inhibitors^75^ as well as other approaches for targeting glucose metabolism^76^ and immunosuppressive Tregs for the treatment of metaplastic breast cancers.

In addition to the spatial characterization of the TME specific to this rare subtype of breast cancer, this integration identified a stromal hub shared across breast cancer subtypes while exhibiting varying spatial patterns. Within this stromal hub, we observed compositional shifts with the replacement of Tregs with activated CD8^+^ T cells in MBC compared to other TNBCs. Additionally, our observation of enriched endothelial cells in MBC stroma alludes to mechanisms of local adaptation to hypoxic regions through possible vascular formation. Altogether, these results imply that the underlying biology of the tumor impacts stromal response and immune infiltration.

Overall, Starfysh has shown the capability to analyze complex spatial transcriptomics, integrate patient samples with distinct microenvironments and tissue sources, and demonstrate robustness in characterizing spatial interactions in within-sample, across-sample replicates as well as across-patient analyses. These features enabled the extraction of new biological insights from a limited cohort of breast cancer patients. In a recent study, we applied Starfysh to disentangle the spatial dynamics of activated and exhausted T cell subsets in Slide-Seq V2^77^ data from anti-PD-1 treated melanoma tumors^78^, showing its applicability to other ST technologies and cancer systems. In future work, the incorporation of archetypal analysis in the probabilistic framework and extensions to multi-omics integration with proteomics or chromatin accessibility will improve our ability to achieve comprehensive characterization of spatial heterogeneity. Additionally, integration with high-resolution images can explicitly account for cell morphology (e.g., nucleus density, nucleus/cytoplasm ratio, etc.), which can be incorporated with priors in the integrative model.

## Methods

### Starfysh model

#### Model overview

Deep generative models parameterized by neural networks have proven effective in analyzing single-cell RNA expression data (scvi-tools^20^, scvi^21^, totalVI^22^, scArches^23^, trVAE^24^, scANVI^25^, MrVI^26^, etc). However, the presence of multiple cell types in each spot in ST data makes it difficult for these models to disentangle the cell-type-specific features. To overcome this limitation, Starfysh introduces a novel generative model with a special variational family that is structured to model the presence of multiple cell states per spot in ST data. The generative model uses geneset signatures (either existing signatures or signatures computed with archetypal analysis) as an empirical prior to help disentangle cell types. The inference method combined with the Starfysh generative model forms an auxiliary deep generative model^27^. We first detail the generative model of Starfysh and then introduce the structured variational family with cell state proportions as auxiliary variables.

#### Starfysh generative process

Starfysh models the vectors of gene expression *x_i_* ∈ ℜ^G^ (with *G* being the number of observed genes) for each spot *i* with a generative model. In the Starfysh generative process (**Fig. 1c, Supplementary Fig. 3a**), each spot *i* is associated with a low-dimensional latent variable *z_i_* ∈ ℜ^M^ (default *M* set to 10 dimensions). This latent variable is a compressed representation of all the cell states contained in the spot. The latent variable *z_i_* is transformed with a neural network *f* into the normalized mean expression of each gene for the spot *i*. This normalized gene expression is then scaled by the library size of the spot *l_i_*. The library sizes are sampled from a log-normal distribution with the mean set empirically to 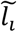, the average observed library size (defined as the sum of counts per spot) in the adjacent spots (detailed in the construction of the empirical prior below). The observed transcripts *x*_ig_ of the gene *g* for the spot *i* are finally sampled from a negative binomial distribution with the previously calculated means and *x_i_* denotes the vector of expression of all genes in the spot *i*. In addition to *z_i_*, Starfysh associates each spot *i* with a vector of cell state proportions *c_i_* ∈ ℜ^K^. The number of cell states *K* is computed in advance with our archetypal analysis. For each cell state *k* = {1,…, *K*} that is discovered during our archetypal analysis, we compute the score of the associated gene expression signature *S_k_*, denoted by *A*(*x_i_*, *S*_k_) (detailed in the construction of the empirical prior below). In each spot, the proportions of cell states are then sampled from a Dirichlet prior distribution with a prior parameter set empirically to the observed scores (*A*(*x_i_*, *S_k_*))_*k*_ of the cell state signatures for the spot. As an example, if *x_i_* highly expresses gene *g*′, a known marker for cell state *k*′, then the score *A*(*x_i_*, *S_k′_*) will be high and our empirical Dirichlet prior parameter will place a higher probability on cell state *k*′. Additionally, a parameter *α* controls the strength of this prior. The generative model is defined as 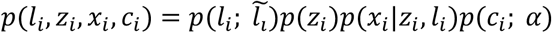 with

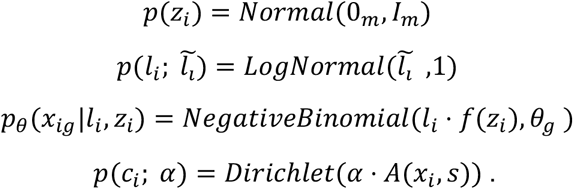

In the generative process above, the average library size observed in spot *i*’s spatial neighborhood is defined by 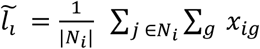 where *N_i_* is the set of spots physically located adjacent to spot *i*, and also includes *i*. The negative binomial dispersions *θ_g_* are gene-specific parameters learned during the inference. The parameters of the neural network *f* with the default including two linear convolution neural layers, followed by ReLU and Softplus activation function respectively, are also learned during inference.

#### Integration with histology image

Though histology H&E images are usually provided along with ST data (e.g., from the commercial Visium platform that aligns spots between replicate tissues), current methods fail to combine the transcriptomic information with the paired histology of the tissue in deconvolving cell types. Histology, however, can provide useful information about morphology, tissue structure, cell density, and spatial dependency of cells. Integrating histology and transcriptomes in a joint model is challenging as the two data modalities are very different: the genome-level transcripts are high-dimensional vectors whereas the histology data consists of three-color channel images. Thus the integrative approach should address the mismatch of these two types of data while preserving the cell-type specific information of gene expression, and cell morphology-specific information of histology images. The integration with histology image in our model is formulated with a deep variational information bottleneck approach^28^.

In the generative model, the histology image is introduced as the variable *y_i_* representing image intensity and is assumed to be generated from the same latent variable *z_i_* that informs gene expression (**Supplementary Fig. 3b**) with a distribution *p*(*y_i_* | *z_i_*) parametrized by two neural networks *g_μ_*,*g_σ_* each consisted of one linear convolutional neural layer followed by a batch normalization layer, for mean and variance of distribution for *y_i_* respectively:

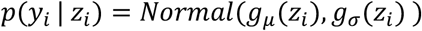

#### Construction of the empirical prior

Based on marker genesets for cell states expected to reside in the tissue, Starfysh first evaluates the feasibility of the provided marker genes and filters genes not expressed in any spots to obtain binary variable *S_k_* ∈ ℜ^*G*^, *k* = {1,…, *K*}. Then, spots enriched for cell states are determined by those ranked in the top N according to mean expression (without normalization) of marker genes for each cell state, forming an initial anchor spot set that can be updated with archetypal analysis explained below. Two priors are calculated before running Starfysh, including a prior for the cell state proportion, and a prior for the library size:

1. Prior for the cell type proportion *A*(*x_i_*, *S_k_*): *A*(*x_i_*, *S_k_*) is defined as the z-scored mean expression: 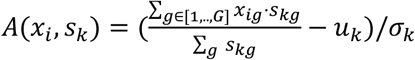 if *i* is an anchor spot and 0 otherwise, where *u_k_* and *σ_k_* denote the expectation and standard deviation of mean expression for cell state *k*. Notably, the prior weight is different between anchor spots and non-anchor spots to enhance the effects anchors, as purest spots with the least mixing, in disentangling cell states in spots with relatively more mixing of cell states.
2. Prior for the library size: Stafysh also considers the spatial dependency of spots when generating the prior for library size. 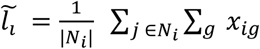 where *N_i_* is the set of spots physically located around the spot *i*, also includes *i*, which denoted. 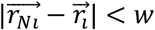. *w* is an adjustable parameter for window size (default set to 3). 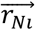 is the coordinate vector for neighbors of spot *i*, and 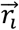 is the coordinates for spot *i*.

#### Archetypal analysis

Marker genes representing cell states may be context-dependent or unknown. To address these limitations and enable the discovery and improved characterization of tissue-dependent and heterogeneous cell states, we developed a geometric preprocessing step, leveraging archetypal analysis^79^, to refine marker genes and identify novel cell states.

Archetypal analysis fits a convex polytope to the observed data, finding the prototypes (archetypes) that are most adjacent to the extrema of the data manifold in high dimension. Previous works^80,81,82^ have applied archetypal analysis on single-cell RNA-seq data to characterize meaningful cell types. In the context of spatial transcriptomics, we hypothesize that the archetypes identify the purest spots that contain only one or the fewest number of cell states, while the rest of the spots are modeled as the mixture of the archetypes.

We applied the PCHA algorithm^83^ to find archetypes that best approximate the “extrema’’ spots on a low-dimensional manifold. Specifically, let *X^norm^* ∈ ℜ^*S*×*G*^ be the log-normalized spot (*S*) by gene (*G*) spatial count matrix such that 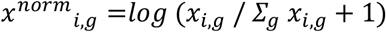. We selected the first P=30 principal components (PCs) (*X*′ ∈ ℜ^*S*×*P*^) to denoise the data. We denote matrices *W* ∈ ℜ^*S*×*K*^, *B* ∈ ℜ^*K*×*S*^, and *H* = *BX*′ ∈ ℜ^*K*×*P*^, where *K* represents the number of archetypes. The algorithm optimizes the parameters of *W* and *B* alternately, minimizing ∥*X*′ − *WX*∥^2^ = ∥*X*′ − *WBH*′ ∥^2^ subject to 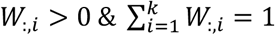, and 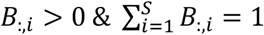 where *S* spot counts and *K* archetypes are convex combinations of each other^84^. To find a suitable choice of *K*, we applied Fisher Separability analysis^85^ to infer the Intrinsic Dimension (ID) as its lower bound, and iterate through different *K* values until the explained variance converges. We also implemented a hierarchical structure to fine-tune the archetypes’ granularity with a resolution parameter *r* ^86^ (default set to 20). For archetype *a_i_*, *i* ∈ 2,…, *k*, if it resides within euclidean distance of *r* from any archetype *a_j_*, *j* ∈ 1,…, *i* − 1, we merge*a_i_* with the closest *a_j_*. The archetypes distant from each other are kept after the shrinkage iteration and used in subsequent steps.

We define *archetypal spots* as the N-nearest neighbors to each archetype by constructing *k* clusters. Then, for each cluster *i*, we identify the top 30 marker genes by performing a Wilcoxon rank sum test between in-group and out-out-group spots with Scanpy^87^. We then designed an iterative pipeline to refine cell state markers according to archetypes. First, we align the archetypal spots with the best 1-to-1 overlap (default 1_*aligned*_= 0.35 (**Supplementary Fig. 1e, Supplementary Fig. 10)** with the initial anchor spots consisting of those with the highest expression of markers for a given cell state, and then append its signature gene list with the archetypal marker genes. Then, we update the anchor spots according to the updated gene list. In practice, the refinement of signature genes to anchor spots can be updated in multiple iterations if needed. Alternatively, to find novel cell states, we rank the archetypal clusters from the most distant to the least distant to the anchor spots of known cell states, and the archetypal clusters distant from all anchor spots represent potential novel states for further study.

The overall archetypal analysis algorithm in Starfysh is summarized as follows:

1. Estimate the Intrinsic Dimension (ID) of the count matrix, and find *k* archetypes that identify the hypothesized purest spots.
2. Find the N-nearest neighbors of each archetype, and construct archetypal spot clusters.
3. Find the most highly and differentially expressed genes for each archetypal spot cluster, and select the top *n* genes (default: *n* = 30) as the “archetypal marker genes”.
4. *<u>If the signature gene sets are provided</u>*: align the archetypal spot clusters to the anchor spots of known cell types, update the signature genes by appending archetypal marker genes to the aligned cell type, and re-calculate the anchors.
5. *<u>If the signature gene sets are absent</u>*: apply the archetypes and their corresponding marker genes as the signatures.

We found that archetypes alone are not sufficient for disentangling cell states (**Supplementary Figure 1e**), however, used as empirical priors to the deep generative model, they can guide the successful deconvolution of cell states (**Supplementary Fig. 1a; Fig. 1e)**.

#### Starfysh structured variational inference without the histology

Given observed ST data, the Starfysh generative model induces a posterior on its parameters. We use variational inference to approximate the posterior. We first describe the inference procedure without the histology integration (no variable *y_i_*). The variables *c_i_* and *l_i_* (cell states and library size) are approximated by amortized mean-field distributions *q*(*c_i_* | *x_i_*) and *q*(*l_i_* | *x_i_*). For the latent variables *z_i_* of the spots, we use a specially structured variational distribution *q*(*z_i_* | *c_i_*, *x_i_*, *l_i_*) that uses the cell-state proportions as an auxiliary variable to sample the latent variables *z_i_*. Because each spot contains multiple cell states with proportions *c_i_*, the structured variational distribution is assumed to decompose as a combination of cell-state-specific terms (denoted by *ζ*(*k*, *x_i_*, *l_i_*) for each cell type *k*), weighted by the proportion of cell-states *c_i_*. The variational family factorizes in the form *q*(*z_i_*, *c_i_*, *l_i_* | *x_i_*) = *q*(*c_i_* | *x_i_*) · *q*(*l_i_*|*x_i_*) · *q*(*z_i_* | *c_i_*, *x_i_*, *l_i_*) (**Supplementary Fig. 3b**). The three distributions are parametrized by neural networks *λ*, *γ* and *ζ* as follow:

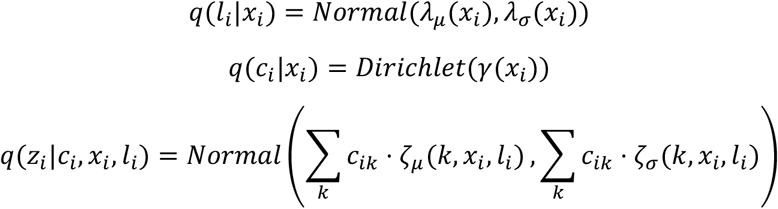

In summary, for each cell state *k*, the function *ζ*(*k*, *x_i_*, *l_i_*) deconvolves the contribution of cell-state *k* to the latent representation of *z_i_*. Each *z_i_* is a combination of the cell-state contributions *ζ*(*k*, *x_i_*, *l_i_*) weighted by the proportions *c_i_*. The cell-state proportions are inferred with the neural network *γ*, which is guided toward the prior to match the cell-type genesets.

Then, the standard variational inference that maximizes the Evidence Lower BOund (ELBO) is performed^88^. The ELBO in our case can be written as:

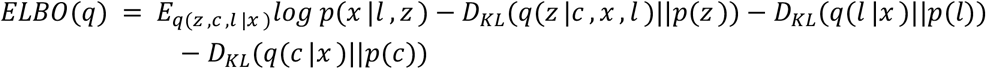

We find the *q* that maximizes the ELBO by running stochastic gradient descent.

#### Starfysh structured variational inference with the histology

To integrate the histology in the inference method, we use the deep variational information bottleneck approach^28^ which models the approximate posterior over the latent variables with the Product of Expert distributions (PoE). For the latent variables *z_i_*, the posterior distribution *q*(*z_i_* | *c_i_*, *x_i_*, *l_i_*) becomes dependent on *y_i_* and we parameterize it as a Product of Expert distribution as described in the original method^28^:

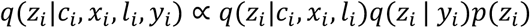

where *q*(*z_i_* | *c_i_*, *x_i_*, *l_i_*) is as described above without the histology, *p*(*z_i_*) is the prior, and *q*(*z_i_* | *y_i_*) is the contribution of the histology image, parameterized as *q*(*z_i_* | *y_i_*) = *Normal*(*h*_μ_(*y_i_*), *h*_σ_(*y_i_*)) with a neural network *h*.

The previous ELBO can be updated with this new variational approximation for the joint modeling of histology and transcriptome. To train such an inference model, we use the information bottleneck approach^28^ instead of just maximizing the ELBO. With this approach, not only the ELBO of the joint model is maximized, but also the ELBO of the transcriptomic model alone, and the ELBO of the histology model alone are optimized. These objective functions are added together to form one objective function to be maximized (**Supplementary Fig. 3c**).

#### Starfysh implementation

The Starfysh model is implemented as a Python package using PyTorch^89^, and optimizations are performed via Adam^90^. The data input is structured to be compatible with Scanpy^87^. Hyperparameters such as early stopping, training times for achieving the best performance, and the number of epochs is adjustable in the Starfysh package with default values provided. Starfysh and tutorials on simulated and real data are publicly accessible (https://github.com/azizilab/starfysh).

#### Prediction of cell-state-specific expression

To predict the cell-state-specific expression, we utilize the decoder in which the parameters have been learned and optimized by the variational inference. The proportion *c_i_* is adjusted to 1 for a specific cell state, and 0 for other cell states. The reconstructed expression and histology are considered as the cell state-specific expression and histology.

### Simulation of ST data

We construct our ST simulation using mixtures of single-cell RNA-seq data previously collected from primary TNBC tumor tissues (CID44971_TNBC)^16^ with three levels of constraints, each building upon the previous one to emulate real-data effects.

#### Spatially-independent simulation

We modify the approach used in Stereoscope^11^ to construct a 5-cell type (Cancer Epithelial, Normal Epithelial, T-cells, CAFs, Myeloids) synthetic ST data. Each spot is independently assigned with the number of cells (5-15) and the number of cell types (1-5) uniformly, and the final cell-type assignment is determined by a uniform Dirichlet distribution. Then, we randomly sample cell indices (with replacement) with the assigned cell type from the scRNA-seq reference. To construct the synthetic ST matrix, selected single-cell expressions for each spot are added and multiplied with a global capture rate parameter.

#### Spatially-dependent simulation

To address the spatial dependencies among neighboring spots, we adopt the pipeline from Cell2Location^8^. Specifically, we define a 50 × 50 matrix, with each pixel representing a synthetic ST spot. We select 5 major cell types (CAFs, Cancer Epithelial, Myeloid, Normal Epithelial, T-cells) from the reference scRNA-seq, and apply 2D Gaussian Process models to simulate their spatial proportions in each spot, respectively. We further assign an expected library size for each synthetic spot with a Gamma distribution fit to the real ST dataset, representing the spatial variation of capture rates among spots. For each spot, we sample single-cell transcriptomes from the reference by searching for candidate cells with a library size closest to the expected library size. The synthetic ST simulation pipeline compares results to the ground-truth cell-type proportions in each spot.

#### Spatially-dependent simulation with paired histology image

To test the impact of multi-modal integration and accounting for spatial dependencies between measurements, cell density, tissue structure, and technical noise effects, we further generate pseudo-histology images paired with the synthetic ST expression counts. Specifically, we design a supervised encoder-decoder neural networks model (**Supplementary Fig. 7**), with real ST expression as input and their histology images as output. First, the expression matrix is projected to a low-dimensional latent space with a ResNet18 encoder, and the histology image is reconstructed with a standard linear decoder with dimension transformation. Around 2000 image patches and corresponding expression matrices are trained from 14 ST samples. A mean-squared loss (MSE) is implemented to fit the predictions to the real ST images. During the generative phase, we use the synthetic ST expression matrix as input to the trained model to obtain their paired synthetic histology image.

#### Signature gene set retrieval in simulated data

For a fair benchmarking not favoring Starfysh, we build the signature gene sets in an unbiased fashion by choosing the top 30 differentially expressed genes (DEGs) for each cell type (highest logFC scores) in single-cell data reported by Wu et al.^16^.

### Benchmarking of Starfysh and comparison to other methods with simulated ST data

We benchmarked Starfysh against reference-based (DestVI, Cell2Location, Tangram, BayesPrism) and reference-free (CARD-free, BayesTME, STDeconvolve) deconvolution methods with the aforementioned simulations. For the reference-based method, we used paired scRNA-seq data for sample TNBC sample CID44971 as the reference. For reference-free methods without inferred cell state annotations, we report the best alignment with the ground-truth proportions upon permutation. We report cell-type specific correlations (ground-truth vs. predicted proportions per spot) as the benchmark metric for each method.

#### Starfysh

We trained Starfysh with 3 independent restarts, where Kaiming initialization is applied to all neural network parameters in Starfysh (**Supplementary Fig. 2**). In each restart, we trained 100 epochs with early-stop, and selected the model with the lowest ℒ_c_. The variational mean *q*_ϕ_(*c_ik_* | *x_ig_*, *l_i_*) is used as the inferred cell-state proportions.

#### BayesPrism

We followed the tutorial on the BayesPrism website: https://www.bayesprism.org/pages/tutorial_deconvolution. We subsetted the common protein-coding gene between the scRNA-seq and ST data with further highly variable gene selection, as the default setting suggested. We ran the BayesPrism Gibbs Sampler *run.prism* with 4 cores and extracted the updated cell-type fractions *θ_n_* as the final deconvolution output.

#### Cell2Location

We followed the tutorial on the Cell2Location website: https://cell2location.readthedocs.io/en/latest/notebooks/cell2location_tutorial.html. We trained the reference regression and Spatial mapping models with 1,000 epochs and 8,000 epochs, respectively. Both parameters were set with converged ELBO loss. The predicted cell-type proportions were obtained from the normalized 5% quantile values of the posterior distribution 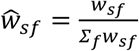.

#### DestVI

DestVI is a single-cell reference and conditional generative model which aims to identify continuous variation in cells that are of the same type via continuous latent variables. We followed the tutorial athttps://docs.scvi-tools.org/en/stable/tutorials/notebooks/DestVI_tutorial.html.

#### Tangram

We followed the Tangram tutorial using default settings: https://github.com/broadinstitute/Tangram/blob/master/tutorial_tangram_with_squidpy.ipynb. We found the optimal alignment for scRNA-seq profiles with 1000 epochs, “cell” mode, and “rna_count_based” reaching a score of 0.957.

#### CARD (reference-free)

We followed the CARD reference-free tutorial: https://yingma0107.github.io/CARD/documentation/04_CARD_Example.html. The default settings were used to generate the cell type proportions (minCountGene = 100, and minCountSpot = 5).

#### BayesTME (reference-free)

BayesTME utilizes a hierarchical probabilistic model that corrects technical errors and models spot-bleeding and then estimates the spot-level UMI (unique molecular identifiers) counts, enabling reference-free analysis of spatial transcriptomic data. We followed the BayesTME deconvolution tutorial (https://github.com/tansey-lab/bayestme/blob/main/notebooks/deconvolution.ipynb), and benchmarked with other methods using the same metric stated above.

#### STDeconvolve

We followed the tutorial on the STDeconvolve website: https://jef.works/STdeconvolve/. We set the optimal number of cell types *K* as 3 and 5, given our 3-type and 5-type simulations, and selected the top 1,000 overdispersed genes suggested by the tutorial. The predicted cell-type proportions were obtained from the output *deconProp*.

##### Quantification of performance in deconvolution of cell types

The performance of each method was quantified by computing the correlations between predicted and ground truth proportions to obtain a matrix *A* (**Fig. 1e**) and its distance to the identity matrix, i.e. ideal deconvolution, with the Frobenius norm *d* = ∥*A* − *I*∥_*F*_. This metric thus evaluates one-to-one (diagonal) mapping of cell types and factors, and penalizes one-to-multi (off-diagonal) mapping.

To evaluate improvement in performance with Starfysh against reference-free methods (STdeconvolve, BayesTME), a Mann-Whitney U test was performed between distance metrics from sampled submatrices by permuting cell states. In particular, for each 5×5 correlation matrix of major cell types, we computed an array of distance values 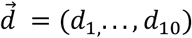 from all *c*(5, 3) = 10 possible 3×3 submatrices {*A*′_1_,…, *A*′_10_} with different cell type combinations, where each *d_i_* = ∥*A*′_*i*_ − *I*_3_∥_*F*_. Then, we tested the distance values obtained from Starfysh against the combination of all other reference-free methods.

### Benchmarking of Starfysh and comparison to other methods with real ST data

We further benchmarked Starfysh with reference-based (Cell2Loation) and reference-free (STDeconvolve) deconvolution methods on TNBC sample CID44971 ST data **(Supplementary Fig. 1)**. We calculated the correlation between the average z-scored expressions of genesets **(Supplementary Table 2)** and the deconvolution profile for each cell state.

#### Starfysh

We followed the same procedure from the simulation benchmark, and reported the variational mean *q*_ϕ_(*c*_*ik*_ | *x*_*ig*_, *l_i_*) as the deconvolution profile.

#### BayesPrism & Cell2Location

For both methods, we followed the same procedures as the simulation benchmark, except for replacing the synthetic ST data with TNBC sample CID44971 real ST data. We applied the TNBC sample CID44971 scRNA-seq annotation from the “subset” classification tier from Wu et al.^16^. For correlation calculation, intersections between single-cell annotations^16^ and our signature cell types are shown, as BayesPrism and Cell2Location only deconvolve cell types that appear in the reference.

#### STDeconvolve

We iterated the number of factors (*k*) from 20 to 30, and choose the optimal *k* as 30 given the lowest perplexity following the tutorial (https://jef.works/STdeconvolve/). Since STDeconvolve doesn’t explicitly annotate factors, we performed hierarchical clustering between the factors (x-axis) and the cell types (y-axis).

#### Archetypal Analysis (Starfysh)

We applied archetypal analysis to the ST data and identified 23 distinct archetypes. We reported the overlapping percentage between anchor spots and archetypal spots for each cell state **(Supplementary Fig. 1e)**.

##### Quantification of performance in deconvolution of cell states in real ST data

Performance in disentangling cell states was evaluated using the same Frobenius norm defined above. To ensure a fair comparison across reference-based and reference-free methods that have different dimensions of correlation matrices due to reference scRNA-seq annotations, we reported a Frobenius norm distance computed as follows: for each method, (1) 1,000 10×10 submatrices {*A*′_1_,…, *A*′_1000_} were sampled from the original correlation matrix *A* without replacement with randomly permuted cell states. (2) An array of Frobenius norm distance 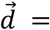 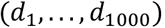, *d_i_* = ∥*A*′_*i*_ − *I*_10_∥_*F*_ was computed; (3) We reported the average value of *d_i_* in **Supplementary Fig. 1**. To test the improvement of Starfysh, we performed a Mann Whitney U-test between the distance array of Starfysh against the combination of all other methods (BayesPrism, Cell2Location, STDeconvolve).

For reference-free methods where the number of inferred factors and the number of cell types may differ, we permuted the correlation matrix such that each cell type (row) is aligned with the factor (column) with the highest correlation score, where the diagonal entries are descending sorted.

##### Runtime comparison across deconvolution methods on real ST data

Runtimes of the core deconvolution function in each method were measured on the same machine with 12-core AMD Ryzen 9 3900X CPU and a GeForce RTX 2080 GPU:

– Starfysh: `run_starfysh` (GPU-enabled)
– BayesPrism: `run.prism`
– Cell2Location: `RegressionModel.train(), Cell2location.train()` (GPU-enabled)
– STDeconvolve: `fitLDA`

### Breast tumor ST data collection and analysis

#### Sample collection and preparation

Tissues were collected from women undergoing surgery for primary breast cancer. All samples were obtained after informed consent and approval from the Institutional Review Board (IRB) at Memorial Sloan Kettering Cancer Center. Samples were obtained from the standard of care procedures. The samples were embedded fresh in Scigen Tissue Plus O.C.T. Compound (Fisher Scientific) and stored at −80C prior to sectioning. 10μm cryosections were mounted on Visium spatial gene expression slides (10x Genomics, #1000184). Two individual tumors were mounted in duplicate on the four 6.5mm × 6.5mm capture areas. The samples were processed as described in the manufacturer’s protocols.

#### Spatial transcriptomics by 10X Genomics Visium

Visium Spatial Gene Expression slides prepared by the Molecular Cytology Core at MSKCC were permeabilized at 37°C for 6 minutes and polyadenylated mRNA was captured by oligos bound to the slides. Reverse transcription, second strand synthesis, cDNA amplification, and library preparation proceeded using the Visium Spatial Gene Expression Slide & Reagent Kit (10X Genomics PN 1000184) according to the manufacturer’s protocol. After evaluation by real-time PCR, cDNA amplification included 13-14 cycles; sequencing libraries were prepared with 15 cycles of PCR. Indexed libraries were pooled equimolar and sequenced on a NovaSeq 6000 in a PE28/120 run using the NovaSeq 6000 SP Reagent Kit (200 cycles) (Illumina). An average of 228 million paired reads was generated per sample.

Tissues were stained with hematoxylin and eosin (H&E) and slides were scanned on a Pannoramic MIDI scanner (3DHistech, Budapest, Hungary) using a 20x/0.8NA objective.

The quality metrics for the collected ST data are shown in **Supplementary Table 5**.

### Analysis of ST data from breast tumor tissues

#### Data preprocessing

Starfysh is compatible with Scanpy^87^ and the preprocessing steps take the raw count matrix as input without normalization after filtering out ribosomal and mitochondrial genes. To account for expression sparsity and noise, we selected the top 2,000 highly variable genes including specified marker genes. By default, Starfysh does not filter low-quality spots by total counts or mitochondrial gene expression ratio, however, these options are provided for users.

#### Identification of tumor-associated anchors (TAAs)

The tumor-associated archetypes were defined as the anchor spots highly associated with tumor cell types. First, an initial set of cell state-enriched spots (e.g., 60 spots for each cell state) and *M* archetypes were identified based on the provided marker gene list and PCHA algorithm, respectively. Since archetypes are vertices non-overlapping with observed data, the 20 nearest neighbor spots for each archetype were identified, obtaining a set of “archetypal spots” as a 20 × *M* matrix. Then, archetypal spots with at least 35% overlap with a set of cell state-enriched spots were considered as aligned to that cell state and their markers are merged. Specifically, this is accomplished by identifying 10 differential expressed genes for the aligned archetypal spots and appending them to the corresponding marker geneset to form a refined marker geneset. Anchor spots were then updated based on the new marker gene list. The final anchors that are associated with any tumor cell geneset (including TNBC, MBC, Luminal A, Luminal B, ER) were considered as tumor-associated anchors (TAAs) (**Fig. 2d,h; Fig. 4c**).

#### Definition of hubs

Hubs were defined as groups of spots with a similar composition of cell states. To integrate ST samples from different patients, anchors are defined on merged data from all samples and Starfysh then infers the cell state proportion and latent variables for each spot in each sample, using the same anchor set. Spots were then clustered according to the inferred cell state proportion using Phenograph clustering (**Supplementary Fig. 18)**.

#### Entropy of spots

We used an entropy-based metric previously used for batch correction in single-cell data^35^ for evaluating the integration of samples. The Shannon entropy of spots denotes the mixing of spots across the samples. Specifically, we constructed a k-NN graph for each spot *i* to determine its nearest neighbors using Euclidean distance in the Starfysh latent space (z). These nearest neighbor spots formed a distribution of patients (*m* ∈ {1,․14}) for overall 14 patients studied in this paper, represented as *e_i_ ^m^*. The Shannon entropy is calculated as 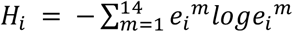. Higher entropy represents higher localized sample mixing across patients (**Fig. 3d**).

#### Kendall’s Tau correlation

Kendall’s Tau correlation is a metric for measuring the ordinal association between two measured quantities. We used this metric to quantify the heterogeneity of tumor-associated anchors (TAAs). Genes for TAAs were ranked based on the differential expression scores for each sample. Samples having similar TAAs were assumed to have a similar rank of differential genes, thus having higher scores of Kendall’s Tau correlation (**Fig. 2p**).

#### Definition of intratumoral, peritumoral and stromal regions

The intratumoral regions are defined as hubs with the mean of inferred proportions of all tumor states being larger than 0.3 (**Supplementary Fig. 19**). Histology information was considered to confirm the enrichment of tumor cells in these regions. Other hubs were ranked by the average distance to intratumoral hubs. With the incorporation of histology and total proportion of immune cells and stromal cells, hub 21 was considered as the boundary between peritumoral regions and stromal regions. Notably, the determined peritumoral regions are shared across all samples while stromal regions are sample-specific.

#### Spatial correlation

To measure the co-localization between cell states, we slightly modified the Spatial Cross-Correlation Index (SCI)^62^. SCI is defined as:

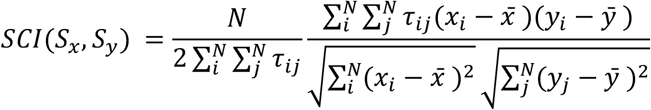

where *x*, *y* denote the predicted proportion for two cell states *S_x_*, *S_y_*, *i*, *j* ∈ [1,․, *N*] are indexes of spots within a certain hub, and 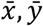 are the mean proportion of two cell states in the hubs. We defined the weight matrix *τ* as information between adjacent neighbors, as: *τ*_*ij*_ = 1 if the coordinate distance of spot *i* and spot *j* was less than 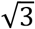 else *w*_*ij*_ = 0.

#### Inference of intercellular ligand-receptor interactions

To investigate the intercellular interactions in a hub, the top 5% spots with the highest inferred proportion of each cell state in the hub were selected. CellphoneDB^56^ was then applied to the selected spots with normalized gene expression. Visualization was performed by the Sankey diagram with `plotly` and Circos plot^91^.

#### Diffusion map analysis with intratumoral hubs

Intratumoral hubs were selected for diffusion map analysis (**Fig. 2h**), and diffusion map components showing gradients between intratumoral hubs were chosen (**Supplementary Fig. 13**). Diffusion map coordinates were used as inputs for the trajectory inference algorithm SCORPIUS^46^. Modules of genes that significantly (q-values<0.05) contributed to the trajectory of transitions between tumor hubs were identified (**Fig. 2i**). Over-representation analysis was conducted to understand the biological processes via Python package `gseapy` with gene sets including KEGG_2021_Human, GO_Biological_Process_2021, and Hallmark.

#### Genes with diffused expression patterns

Treg-enriched (proportion>0.05) spots in intratumoral hubs were selected, and the distance between all spots to the selected spots were calculated with the `sklearn.neighbors` Python package with the function `KDTree`. For each gene, the expression of spots with the same distance were averaged and smoothed with a window size of 7 for each sample. The mean and standard deviation of expression across all samples was computed and smoothed with `Gaussian_filter1d(sigma = 1.5)` with Python package `scipy` (mean and SD shown as solid line and shaded area in **Fig. 4k**).

## Supporting information

Supplementary table 3 Inferred cell state proportions by Starfysh

Supplementary table 5 Spatial transcriptomics qc metrics

Supplementary table 2 Marker gene list

Supplementary table 4 Metabolic signatures

## Data availability

The raw data discussed in this manuscript will be deposited in the National Center for Biotechnology Information’s Gene Expression Omnibus.

## Code availability

The code to reproduce the results in this manuscript is available on the GitHub repository (https://github.com/azizilab/starfysh) and has been deposited to Zenodo (https://zenodo.org/badge/latestdoi/520294385). The reference implementation of DestVI, RCTD, BayesTME, along with accompanying tutorials, is available at the GitHub repository too.

## Acknowledgment

We thank Benjamin Izar and Yiping Wang for fruitful discussions. We also thank Justin Hong for assistance with the Starfysh package and tutorials. We acknowledge the use of the Precision Pathology Biobanking Center, Integrated Genomics Operation Core, and the Molecular Cytology Core, funded by the NCI Cancer Center Support Grant (CCSG, P30 CA08748), Cycle for Survival, and the Marie-Josée and Henry R. Kravis Center for Molecular Oncology. Y.J. acknowledges support from the Columbia University Presidential Fellowship. J.L.M-F is supported by the National Institute of Health (NIH) National Human Genome Research Institute (NHGRI) grant R35HG011941 and National Science Foundation (NSF) CBET 2146007. D.B. is supported by NSF IIS 2127869, ONR N00014-17-1-2131, ONR N00014-15-1-2209. K.W.L is supported by NIH UH3 TR002151. A.Y.R. is supported by NIH National Cancer Institute (NCI) U54 CA274492 (MSKCC Center for Tumor-Immune Systems Biology) and Cancer Center Support Grant P30 CA008748, and the Ludwig Center at the Memorial Sloan Kettering Cancer Center. A.Y.R. is an investigator with the Howard Hughes Medical Institute. G.P. is supported by the Manhasset Women’s Coalition Against Breast Cancer. E.A. is supported by NIH NCI grant R00CA230195 and NSF CBET 2144542.

## Author contributions

E.A., G.P., A.Y.P. conceived of the study and provided overall supervision of the study. S.H., Y.J., A.N., and E.A. designed and developed Starfysh. G.P. provided clinical samples. S.R., B.D., I.V. prepared the samples and performed ST data acquisition experiments. S.H., Y.J., L.S., X.C., L.E.F., J.L.F., C.Y.P., R.M., Y-H.L., D.C., K.W.F., K.M., M.R. analyzed and interpreted the data. J.L.M-F., D.B., K.W.L. provided additional supervision. S.H., Y.J., A.N., L.S., A.Y.R, G.P., E.A. wrote the manuscript. All authors reviewed, contributed to, and approved the manuscript.

---

**Supplementary table 1.** Patient clinical information

| Patient ID | Replicates | ER | PR | Her2 | Age | Subtype |
| --- | --- | --- | --- | --- | --- | --- |
| P1_ER | 2 | 95 | 80 | - | 70 | Ductal (invasive) |
| P2_TNBC | 2 | 2 | 0 | - | 84 | Ductal (invasive) |
| P3_MBC | 2 | 0 | 0 | - | 71 | Metaplastic (maxing producing) |
| P4_MBC | 2 | 0 | 0 | - | 52 | Metaplastic (matric producing) |

**Supplementary table 2. Marker genesets for tumor epithelial, immune, and stromal cells in breast tumor tissues**

**Supplementary table 3. Inferred cell state proportions by Starfysh**

**Supplementary table 4. Genesets for metabolic pathways**

**Supplementary table 5. Spatial transcriptomics quality control metrics**

Supplementary tables 2-5 are included in the supplementary material.

## Supplementary figures

**Supplementary Figure 1.**
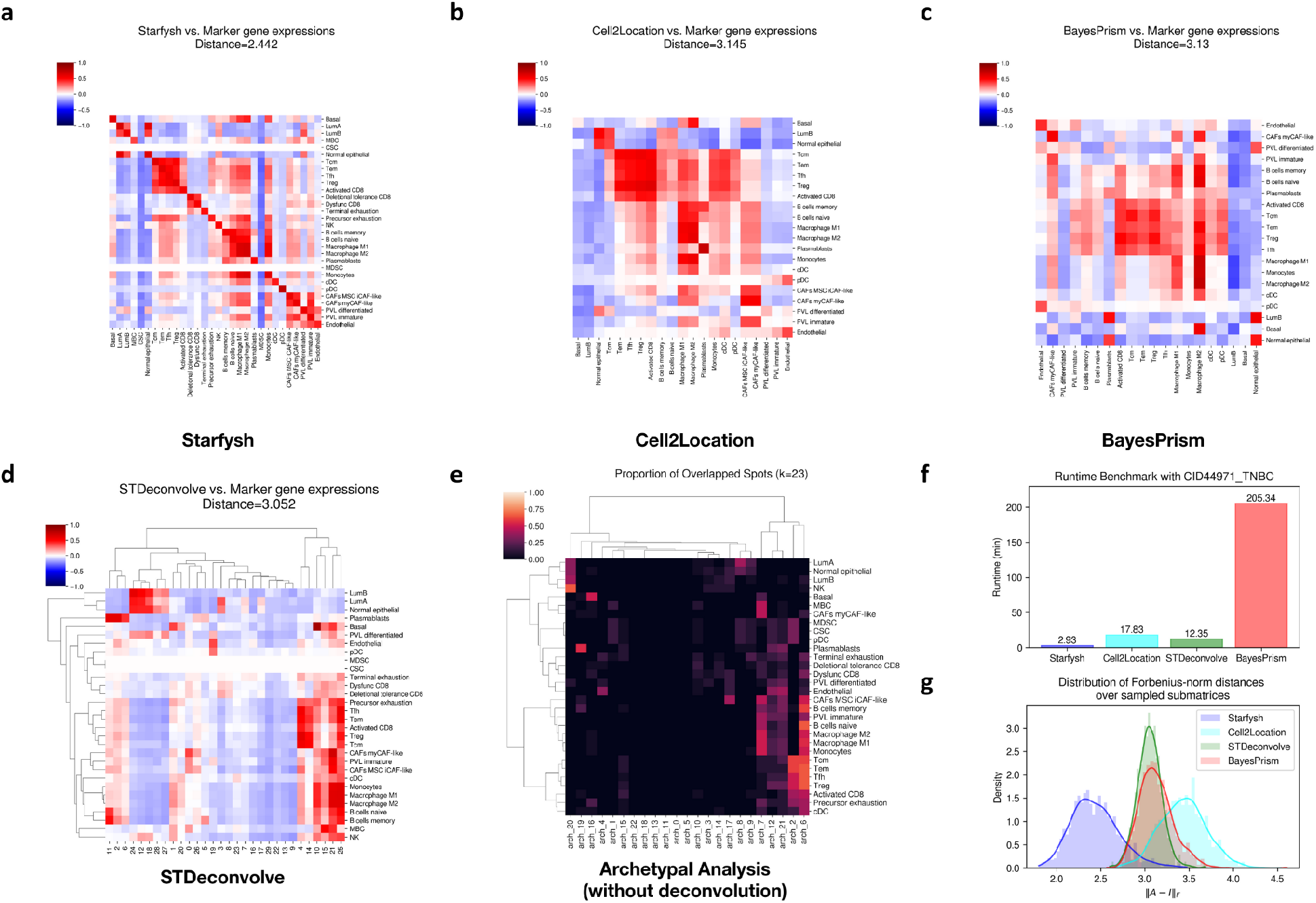
Performance in disentangling refined cell states in ST data from a TNBC Patient CID44971 ^19^ and comparison to reference-based and reference-free methods. Pearson correlation computed between the z-scored average expressions of cell-state specific genesets curated according to matched single-cell data^19^ and inferred proportions from deconvolution using **(a)** Starfysh. **(b)-(c)** reference-based methods: (b) Cell2Location and (c) BayesPrism. **(d)**. Reference-free method STDeconvolve. The performance of each method is summarized by computing the distance between the correlation matrix and an identity matrix defined as the Frobenius norm of the difference (**Methods**). Starfysh shows a significant improvement over other methods (Mann Whitney U test on permuted cell states p<1e-30). **(e)** Archetypal analysis without fitting Starfysh does not show interpretability and correspondence to cell states as seen with Starfysh (a) (**Methods**). (f) Runtime measured for each method applied to sample CID44971. (g). Distribution of Frobenius norm distance ∥*A*′ − *I*_10_∥_*F*_ over 1,000 sampled matrices from each method (**Methods**).

**Supplementary Figure 2.**
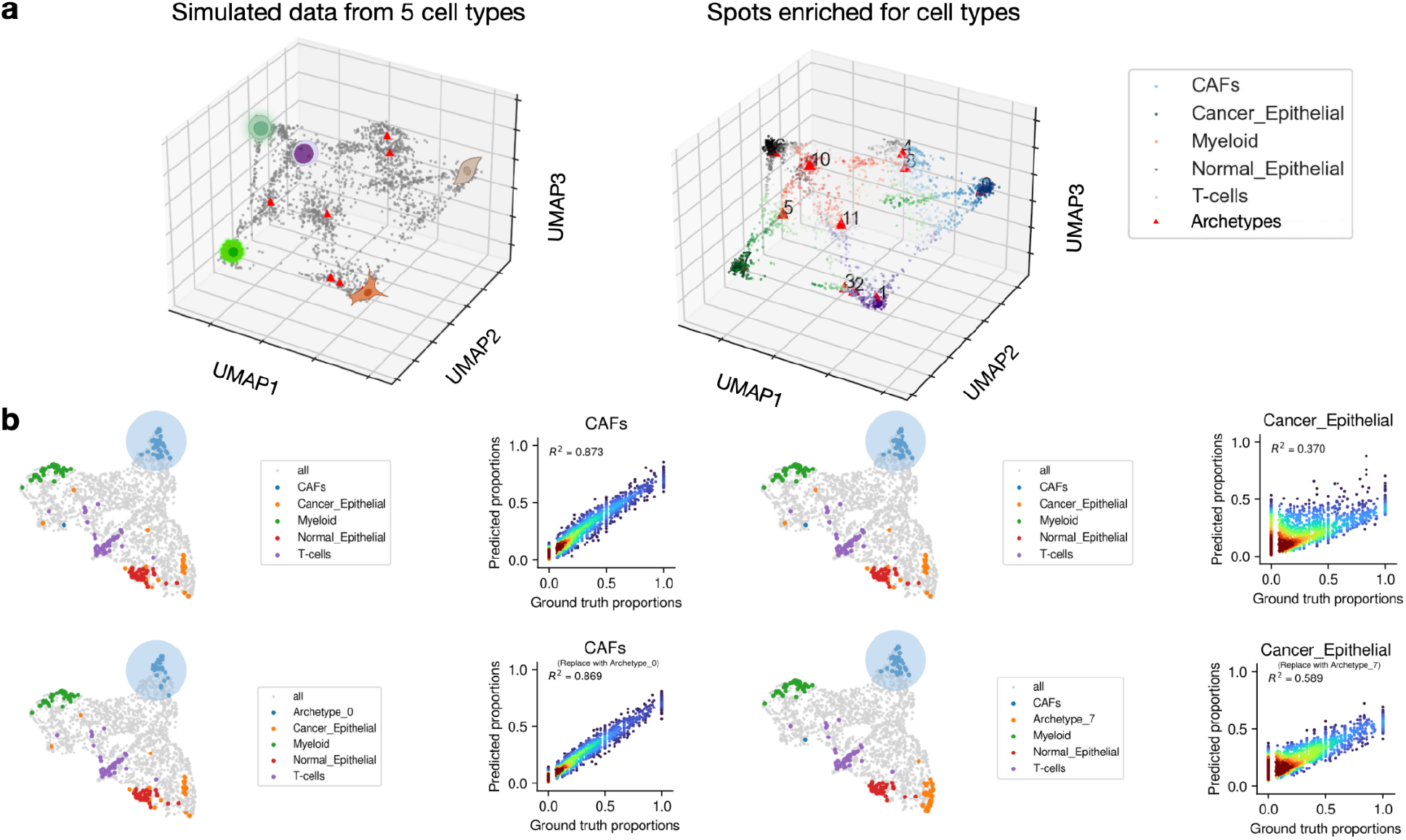
Archetypal analysis with the simulation of 5 cell-types. (a) 3D UMAP of ST data simulated (left) from mixtures of single cell data from real ST data (TNBC sample CID44971)^19^; each grey dot represents a spot; archetypes are denoted by red triangles. Inferred archetypes overlapping with spots enriched for one cell type (purest spots), and assist to refine inaccurate anchors due to incomplete marker genes (e.g., Cancer Epithelial) (**Methods**). Spots are colored according to the proportion of the most abundant cell type (right). (b) Verifying that archetypal analysis assists in refining the anchor spots and improving the deconvolution. Left: Replacing CAFs anchor spots (inferred from marker genesets) with the best aligned archetypes obtains on-par accuracy. Right: Replacing Cancer Epithelial with the most distant archetypal cluster to any anchor spots resulted in significant improvement in Cancer Epithelial deconvolution.

**Supplementary Figure 3.**
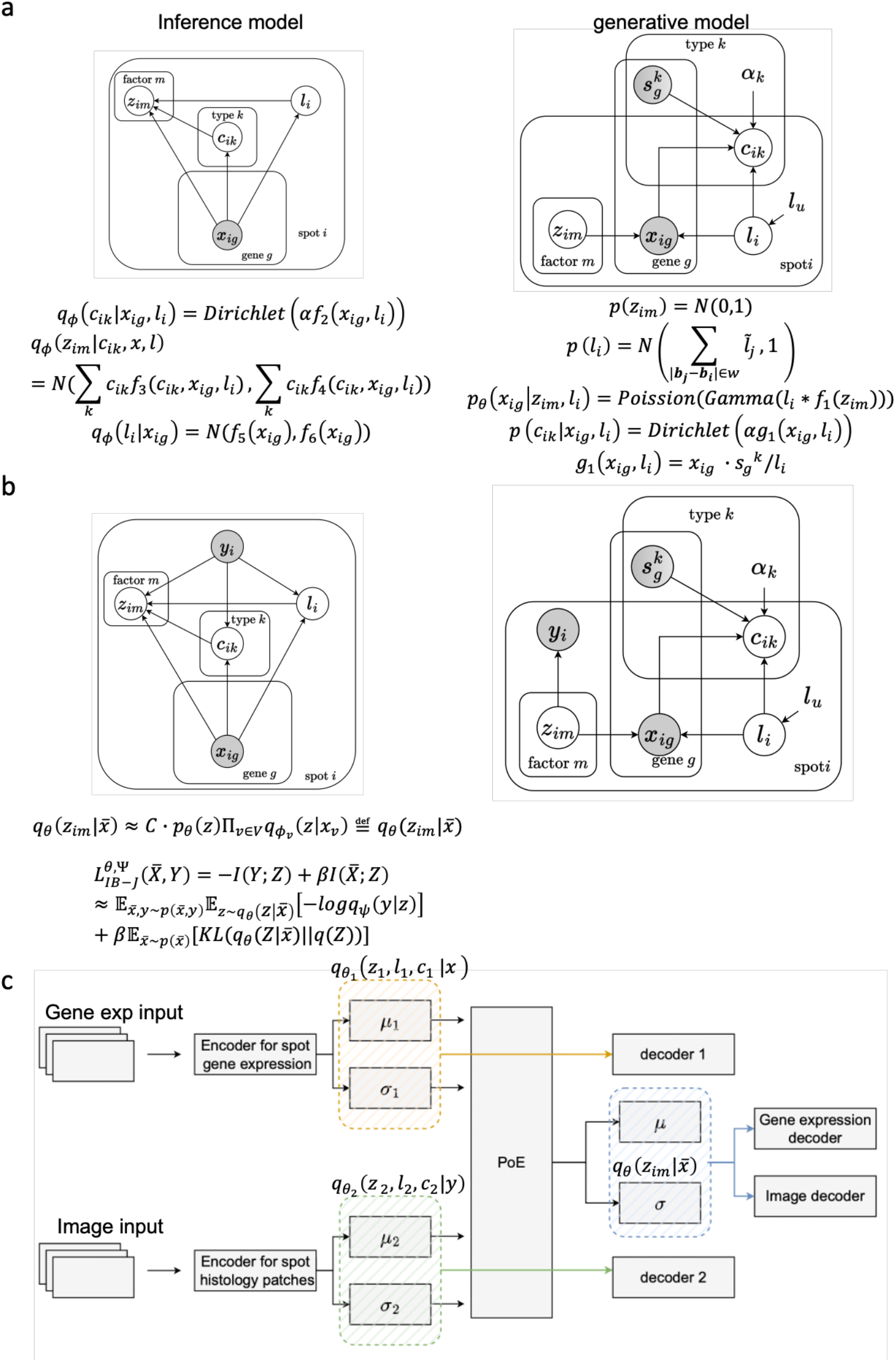
Details of the Starfysh model. (a) Inference and generative model of Starfysh without histology. (b) Inference and generative model of Starfysh with histology. (c) Product of experts integrating images and transcriptomic data.

**Supplementary Figure 4.**
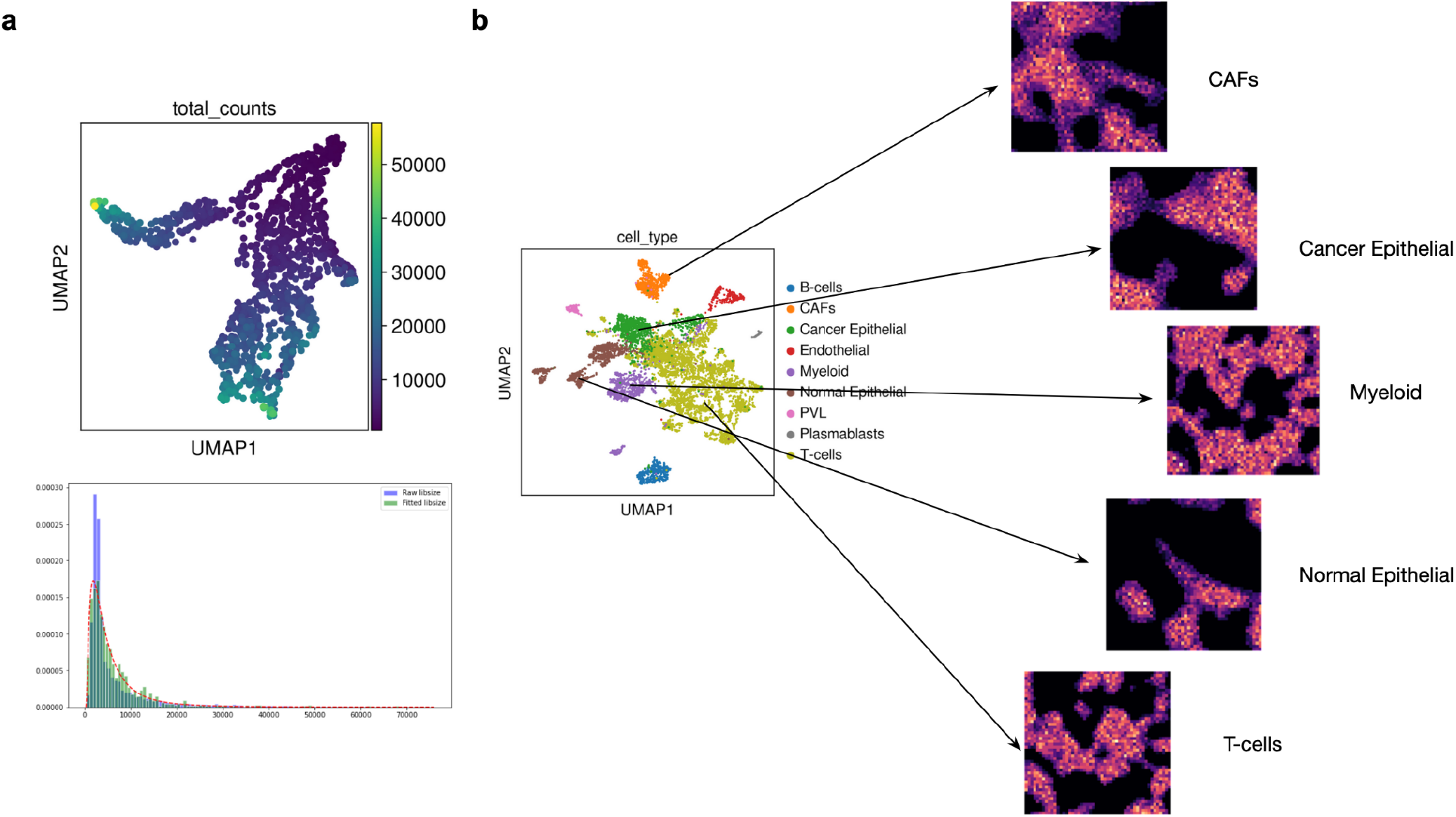
Simulating ST data from primary breast tumor tissues. (a) We simulated the library size of synthetic ST data with a Gamma distribution fit to real ST data (TNBC sample CID44971)^19^. (b) We sampled major cell types from real single-cell matched data (TNBC sample CID44971) to construct gene expressions and proportions of the synthetic spots with spatial dependencies using a 2D Gaussian Process (Methods).

**Supplementary Figure 5.**
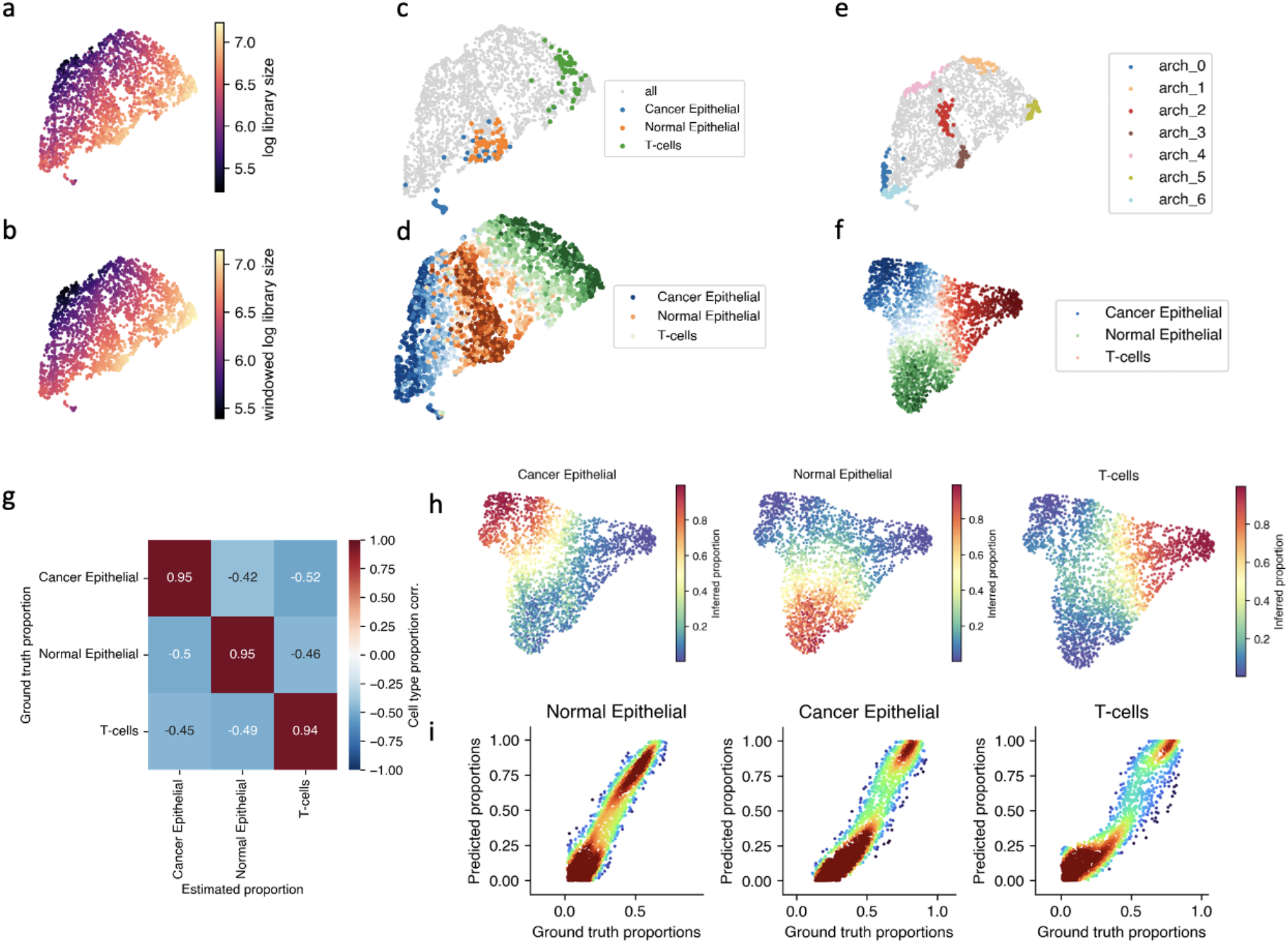
Performance on simulated data from 3 cell type without spatial dependencies. (a-d) UMAP projection of simulated data colored by (a) log library size (i.e., total counts per spot). (b) Log library size of simulated ST data (smoothed with neighboring spots (Methods). (c) Spots enriched for cell types. (d) Ground truth proportion of most enriched cell type. (e) Anchor spots corresponding to archetypes. (f) UMAP of inferred z from Starfysh colored by proportion of most enriched cell type. (g) Correlation between gene signature expression and ground-truth proportion. (h) Same as (f) colored by inferred proportions of cell types. (i) Scatter plot of inferred proportion vs. ground-truth proportions per spot. Spots are colored by density.

**Supplementary Figure 6.**
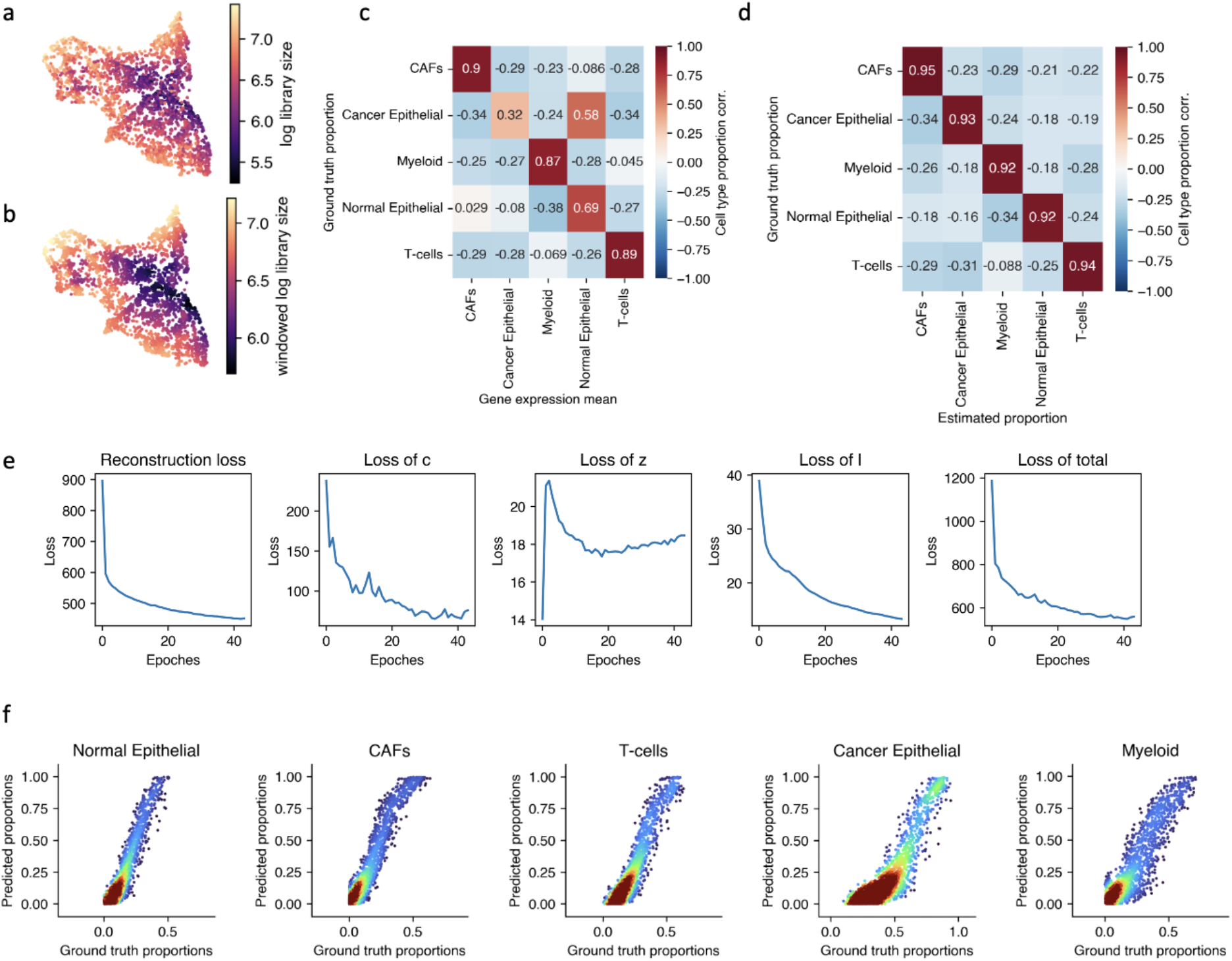
Performance of simulated data with 5 cell types without spatial information. (a-b) UMAP projection of simulated data colored by log library size (a) and smoothed log library size (b). (c) Correlation between gene signature and ground-truth proportion. (d) Correlation between inferred proportion by Starfysh and ground-truth proportion showing improved deconvolution compared to using geneset expressions alone. (e) Model training loss. (f) Scatter plot of inferred proportion vs. ground-truth proportions. Spots are colored by density.

**Supplementary Figure 7.**
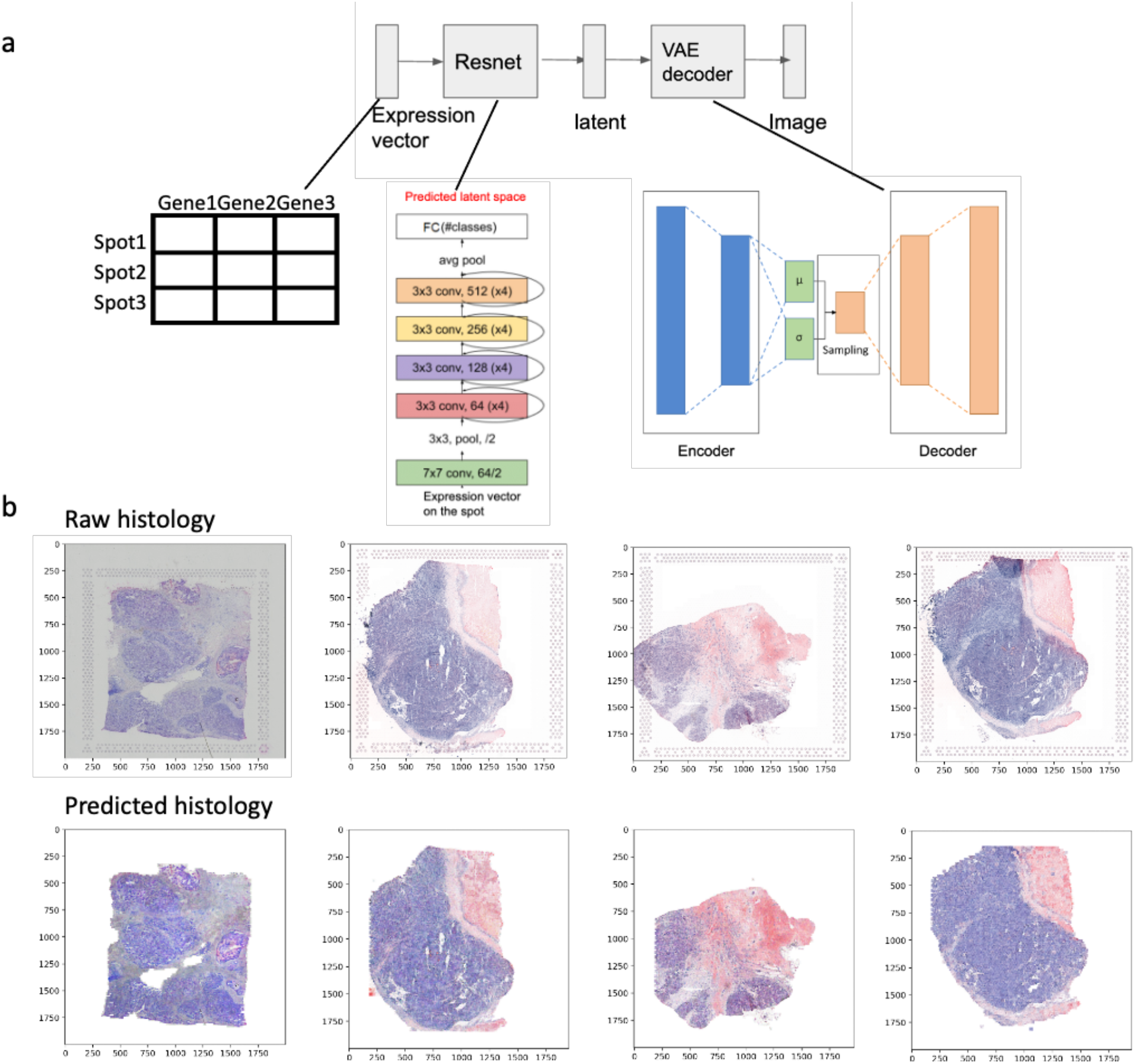
Simulating histology imaging data from gene expression. (a) The model components. (b) Example results on predicted images from gene expression data compared to ground truth histology images.

**Supplementary Figure 8.**
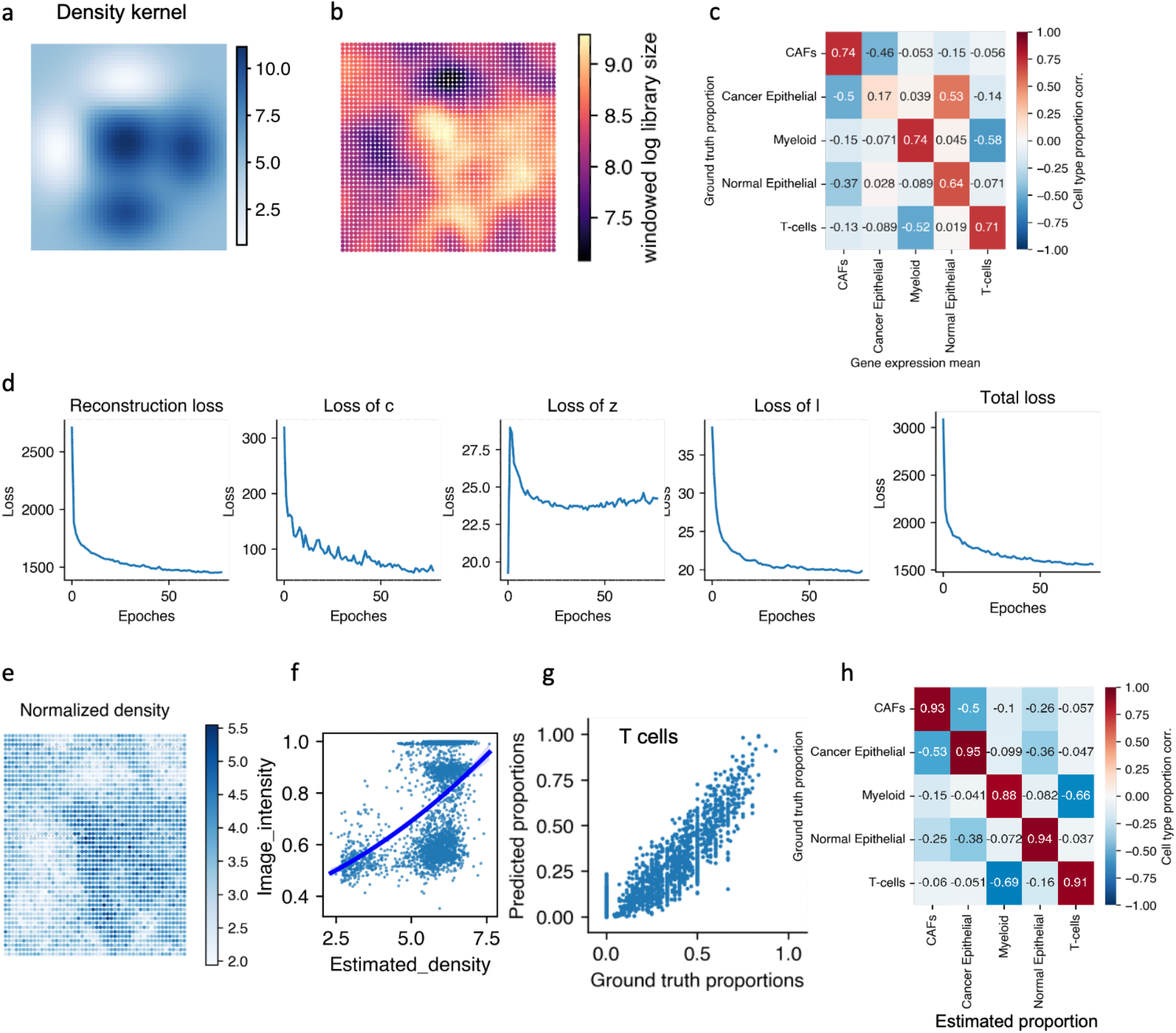
Performance of simulated data with 5 cell types with spatial information. (a) Density kernel added in data simulation to construct spatial dependencies. (b) Smoothed log library size. (c) Correlation between gene signature and ground-truth proportion. (d) Model training loss. (e) Estimation of cell density. (f) Correlation between inferred density and pixel intensity of histology image. (g) Scatter plot of inferred proportion vs. ground-truth proportions. (h) Correlation between ground-truth proportion and inferred proportion with spatial information.

**Supplementary Figure 9.**
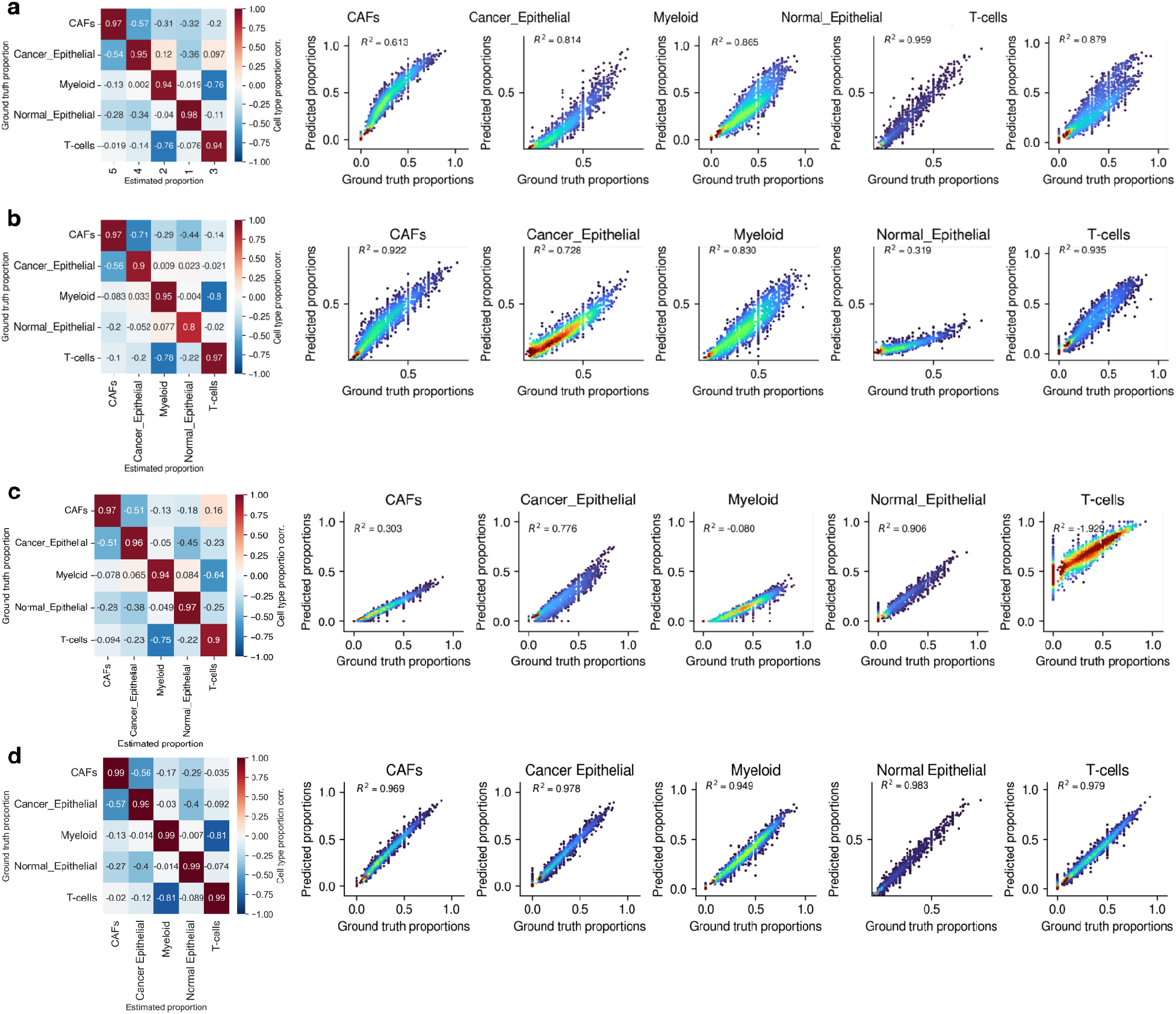
Benchmarking on simulated data (with spatial information) for 5 cell types. Left: Heatmap of Correlation between inferred proportion vs. ground-truth by STDeconvolve (a), Cell2Location (b), Tangram (c), and BayesPrism (d). Right: Scatter plots of inferred vs. ground-truth cell-type specific proportions. Spots are colored by density.

**Supplementary Figure 10.**
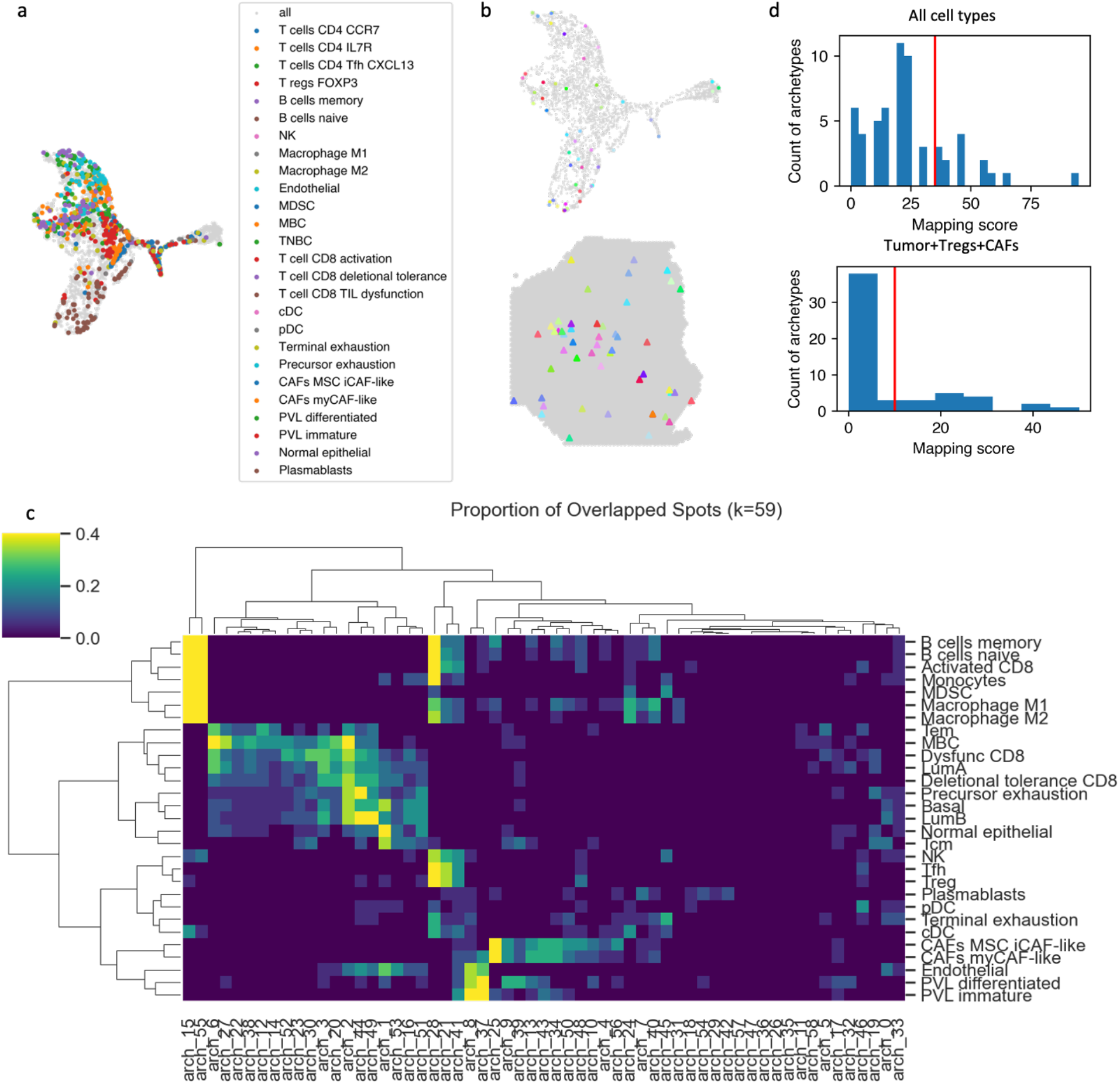
Mapping score of all archetypes with more than 35% overlaps with cell state-enriched spots. **(a)** UMAP of ST data from sample P2A_TNBC highlighting spots with cell state-enriched spots in color. (b) UMAP with archetypes shown in color. (c) Heatmap of the proportion of overlap between archetypes and cell state-enriched spots (d) Histogram of mapping scores between archetypes and cell states (red: data-driven choice of mapping threshold)

**Supplementary Figure 11.**
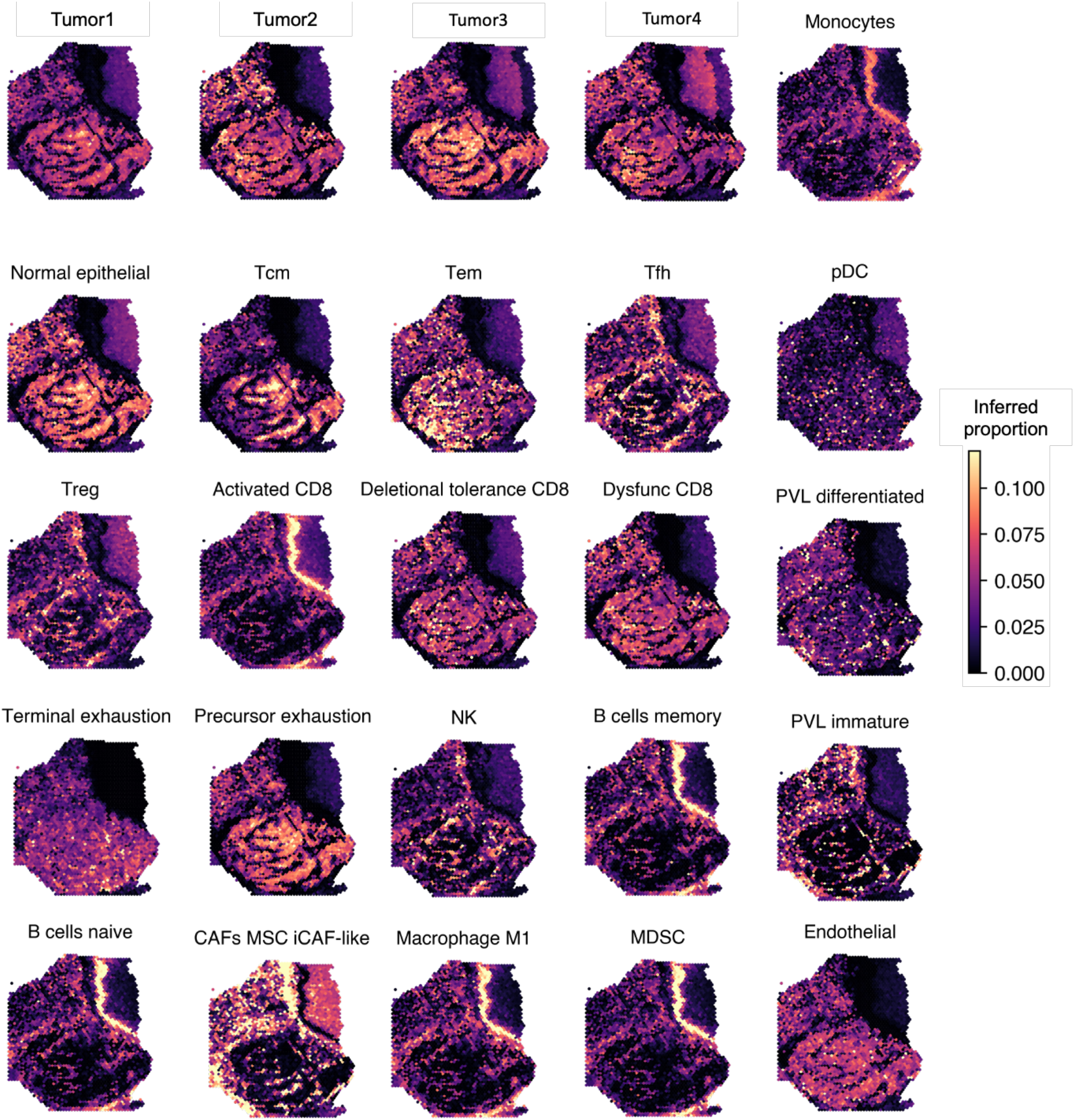
Inferred cell state proportions on P2A_TNBC by Starfysh.

**Supplementary Figure 12.**
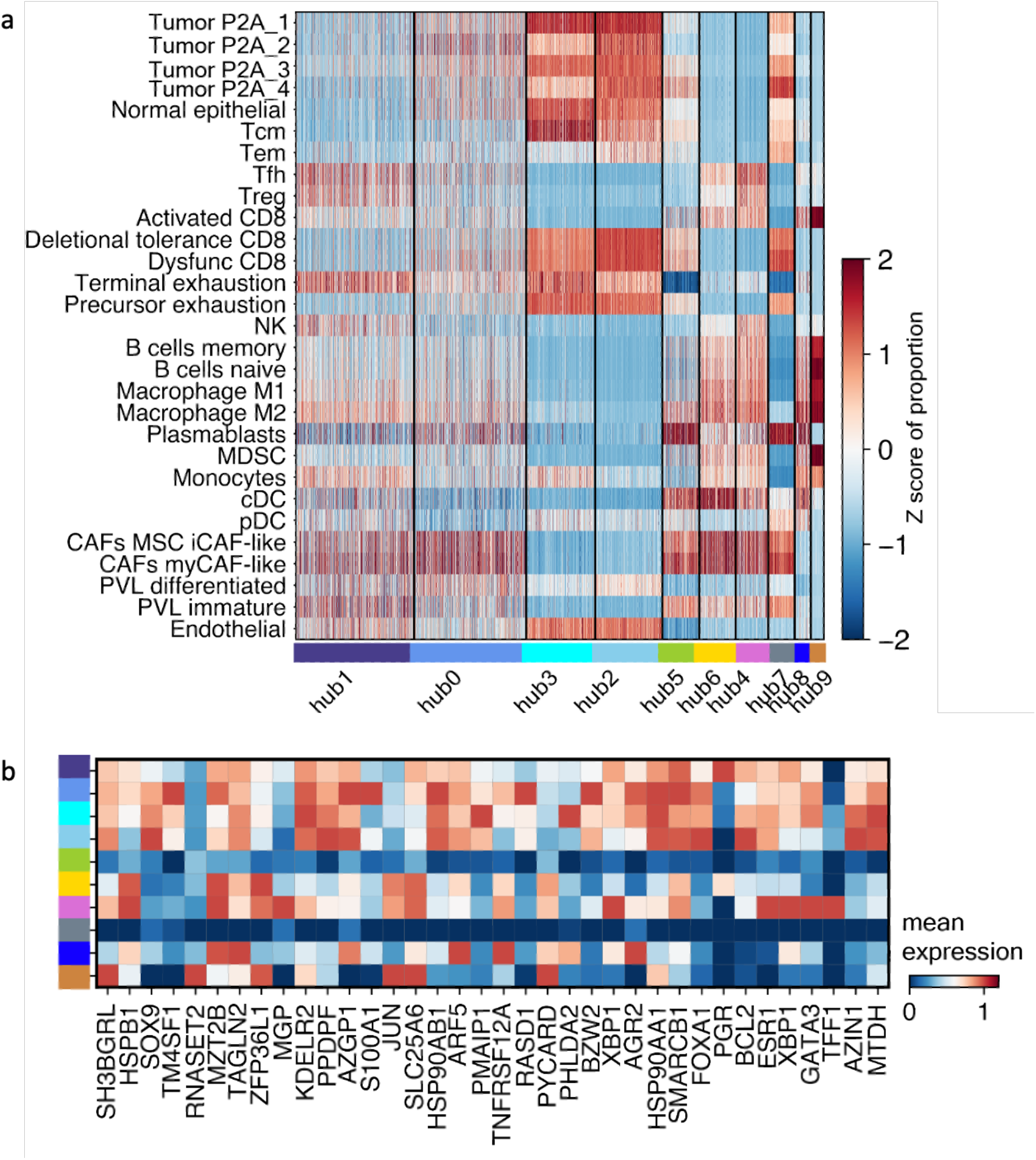
Definition of spatial hubs and tumor states in P2A_TNBC. (a) Heatmap of inferred proportions of cell states (rows) in spots (columns) grouped by hubs corresponding to Fig. 2a-l. (b) Heatmap of marker gene expression for LumA-like tumor cells in P2A.

**Supplementary Figure 13.**
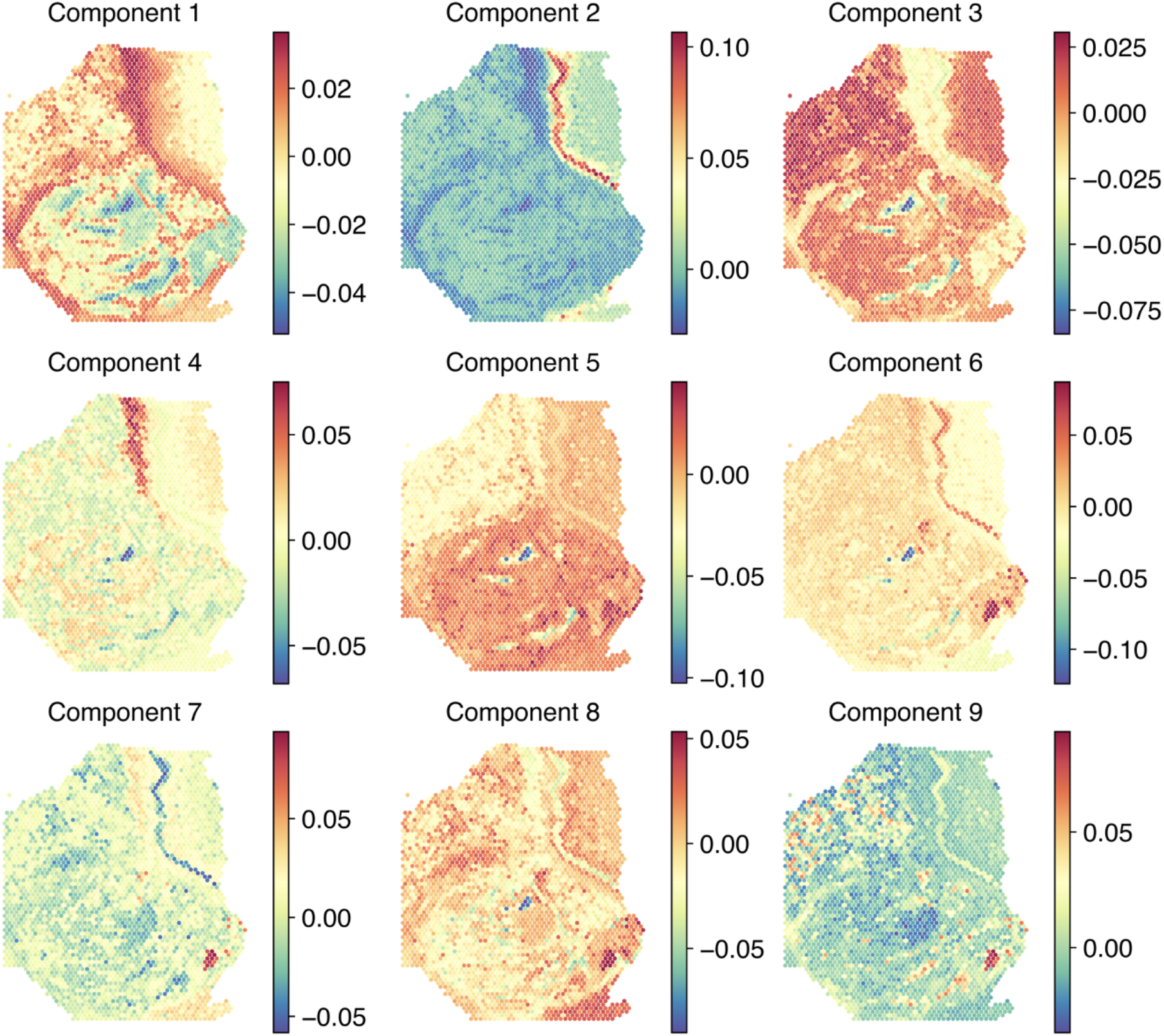
Spatial distribution of diffusion components in P2A_TNBC.

**Supplementary Figure 14.**
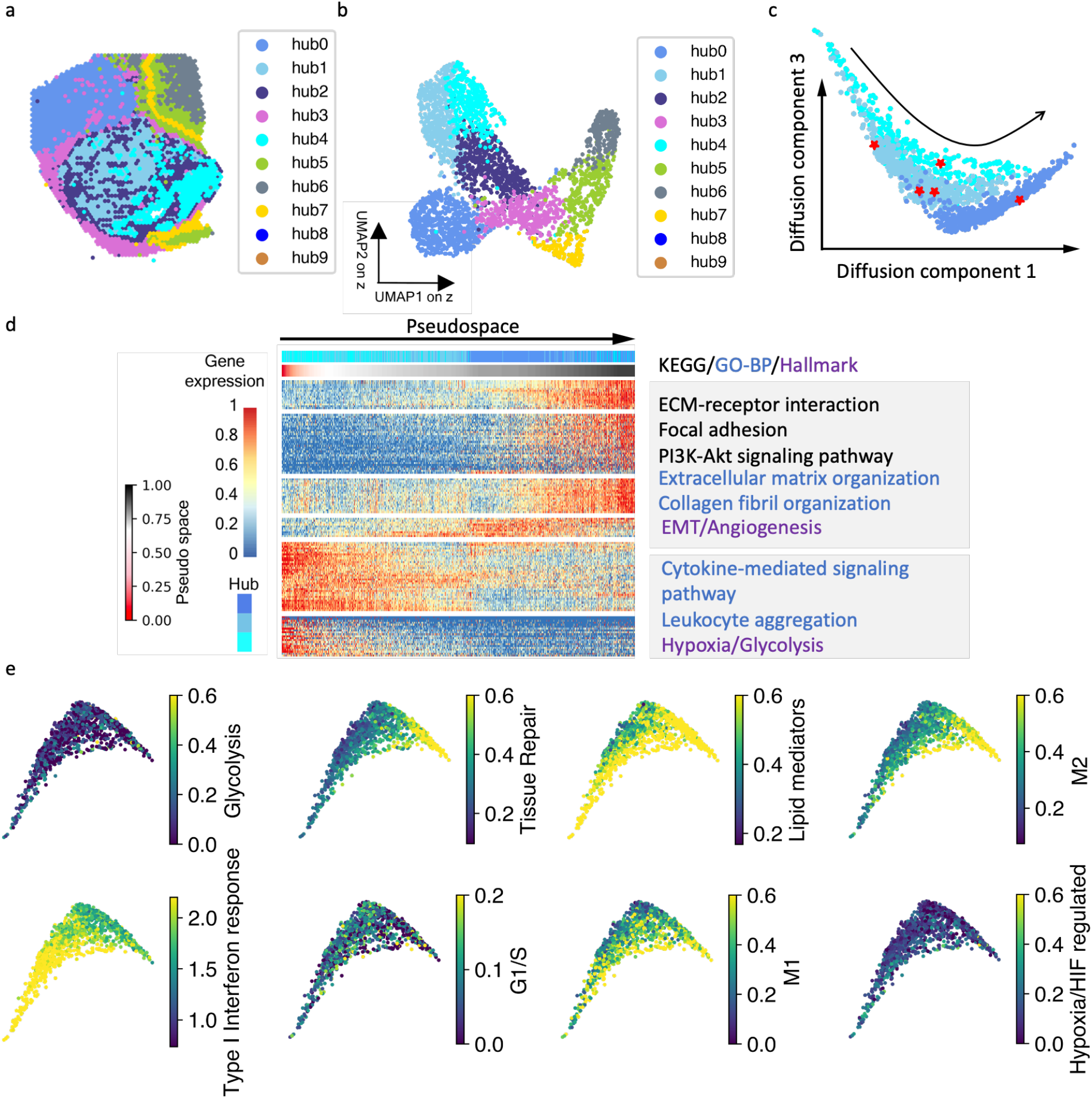
Starfysh characterized spatial heterogeneity in P2B_TNBC. (a) Spatial arrangement of hubs in the context of the issues for P2B (adjacent tissue slice to P2A shown in Fig. 2). (b) UMAP embedding of spots colored with spatial hubs. (c) Diffusion map analysis reveals a continuous trajectory between spots enriched for tumor cell states (hub 1,2,3,5,6). (d) Heatmaps of expression of modules of genes with positive or negative correlation with the projection of cells along the trajectory. (e) Metabolic signatures from intratumoral hubs of patient P2A_TNBC. Expression of metabolic signatures (**Supplementary Table 4**) found in the P2B_TNBC dataset was averaged and plotted on the embedding of diffusion components of tumor transition.

**Supplementary Figure 15.**
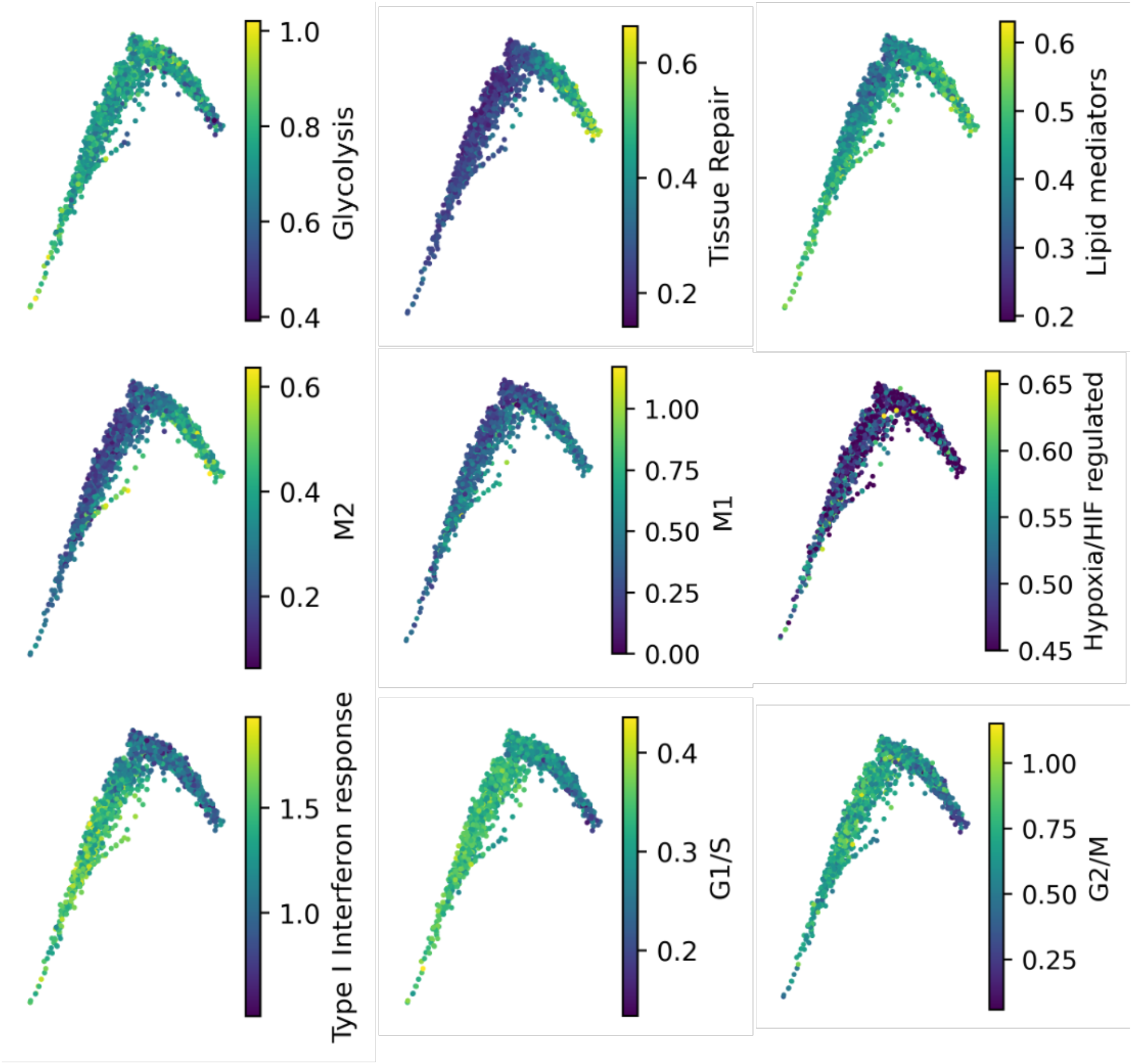
Metabolic signatures from intratumoral hubs of patient P2A_TNBC. Expression of metabolic signatures (**Supplementary Table 4**) found in the P2A_TNBC dataset was averaged and plotted on the embedding of diffusion components of tumor transition.

**Supplementary Figure 16.**
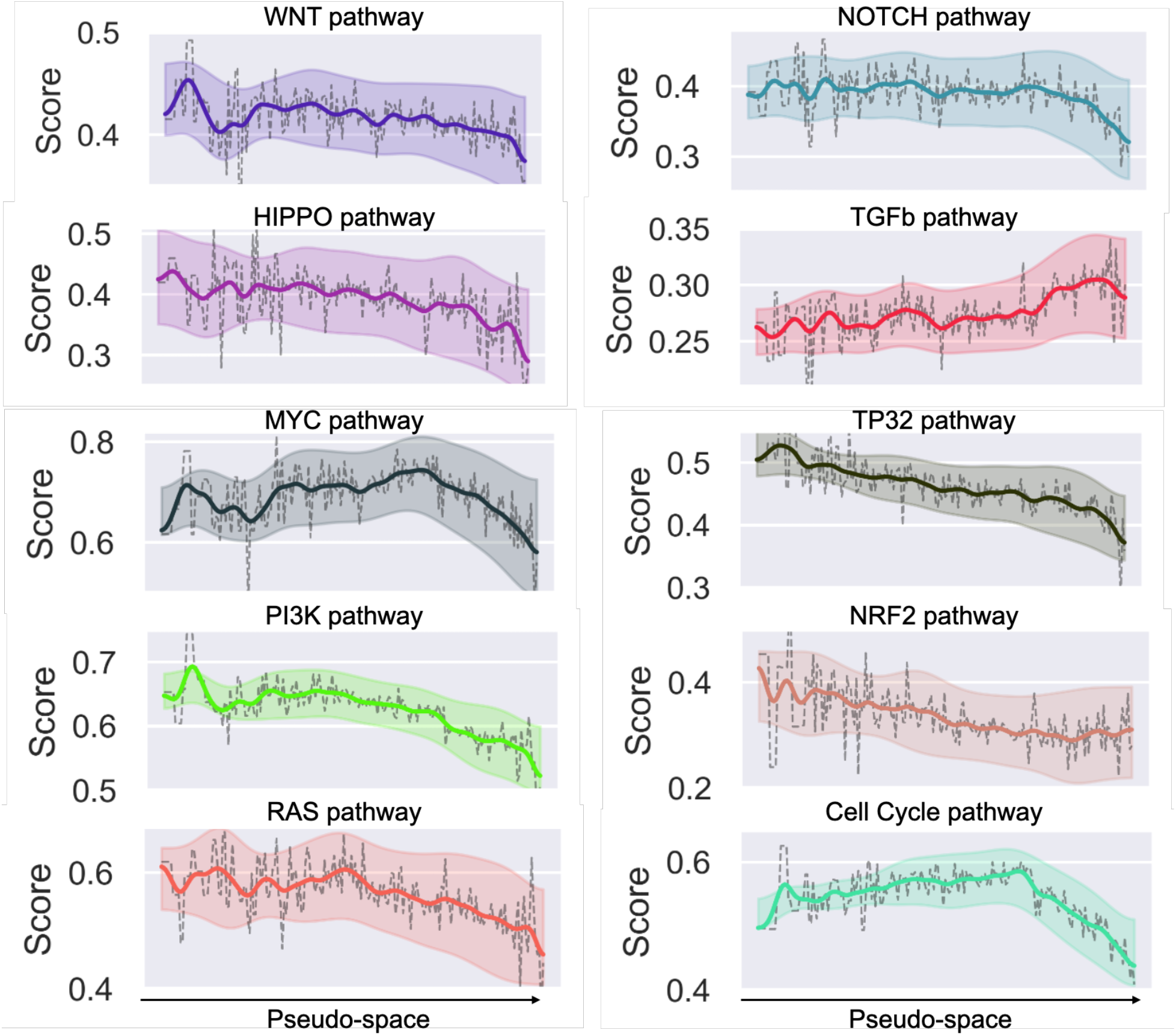
Oncogenic pathways associated with different tumor state transition in patient P2A_TNBC. The score represents the mean expression of genes associated with oncogenic pathways. Dashed lines denote the mean score of spots along with pseudo-space axis. Solid lines denote values smoothed by Gaussian Filter with sigma = 4. The shaded area represents the standard deviation smoothed by Gaussian Filter with sigma = 10.

**Supplementary Figure 17.**
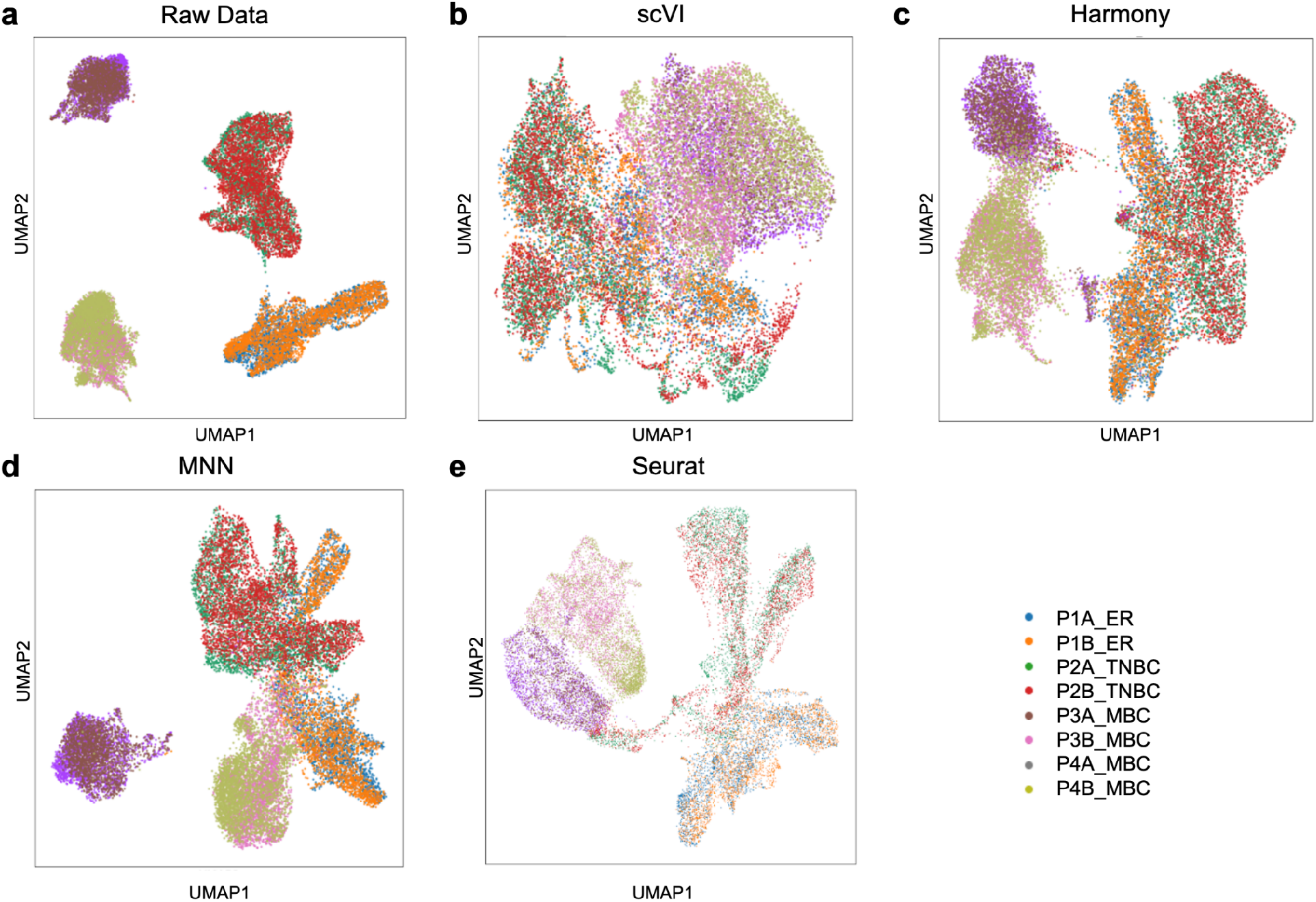
UMAP visualization of batch integration on breast tumor samples with other methods. (a) UMAP of raw counts (b)-(e) Integration with scVI (b), Harmony (c), MNN (d), and Seurat 4.0 (e).

**Supplementary Figure 18.**
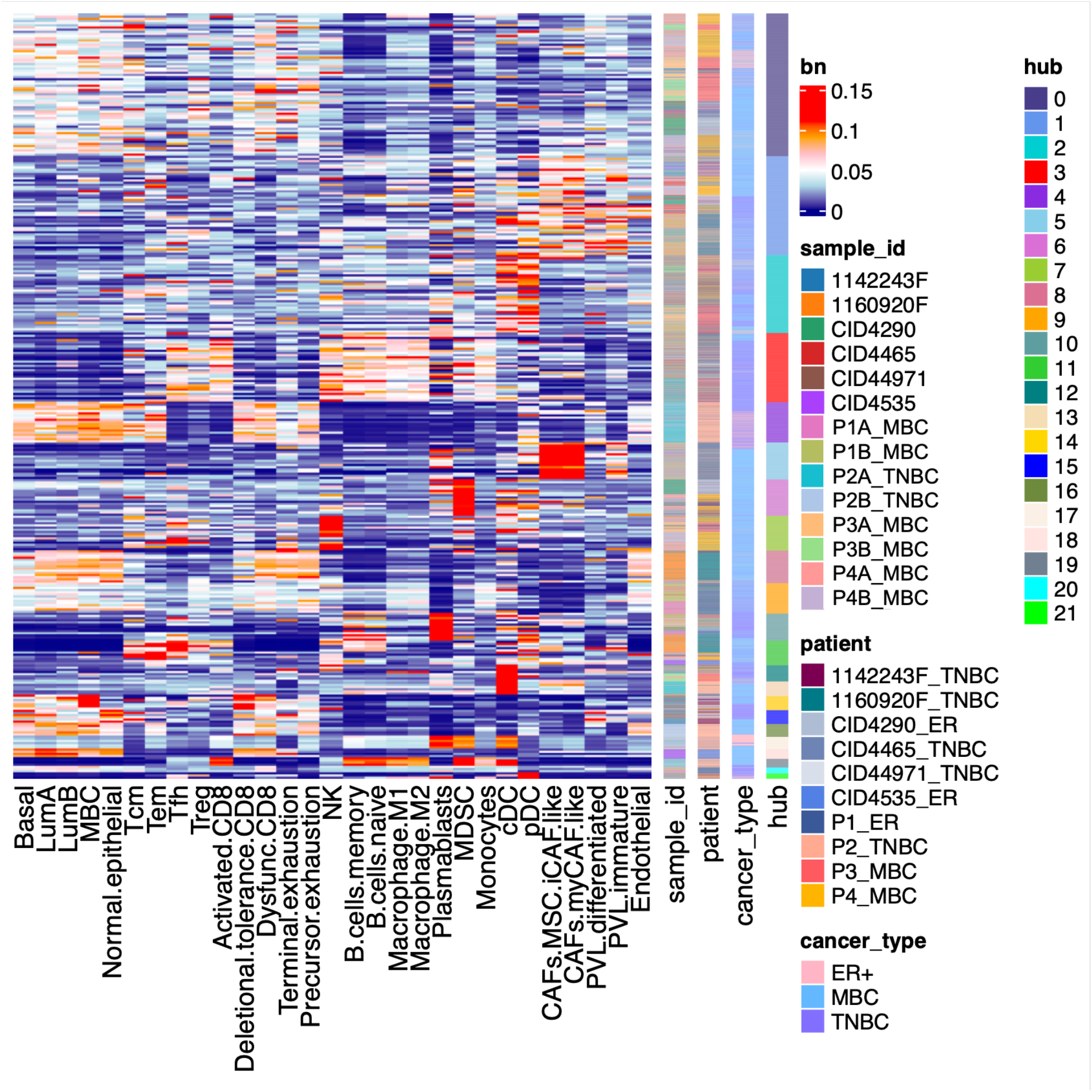
Heatmap of inferred factors in all samples with each cell state. The heatmap shows cell states as columns and spots as rows. Each spot is labeled by sample names, patient names, cancer types, and hubs. Spots are ordered by inferred hubs.

**Supplementary Figure 19.**
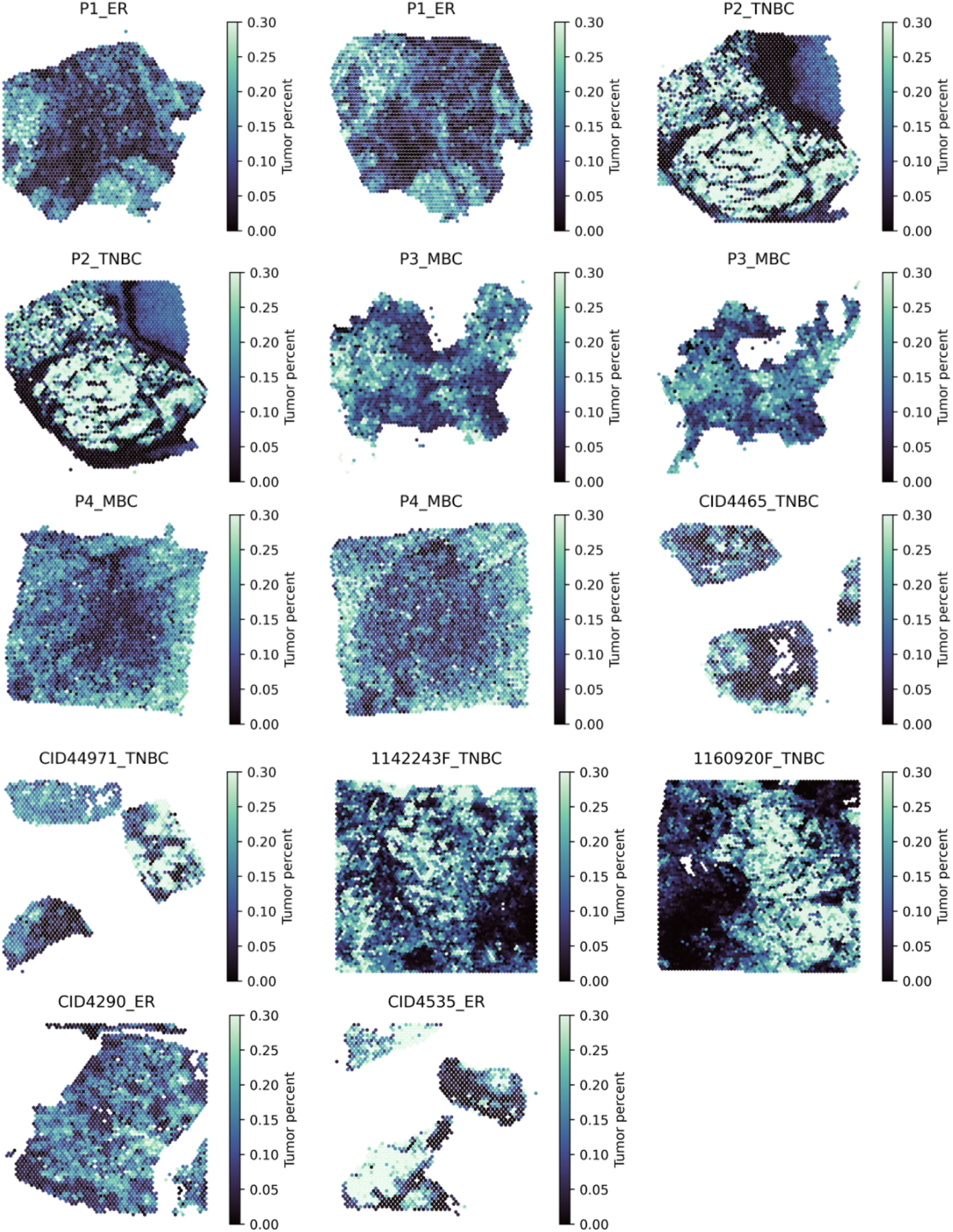
Spatial distribution of percentages of any tumor states in each sample.

**Supplementary Figure 20.**
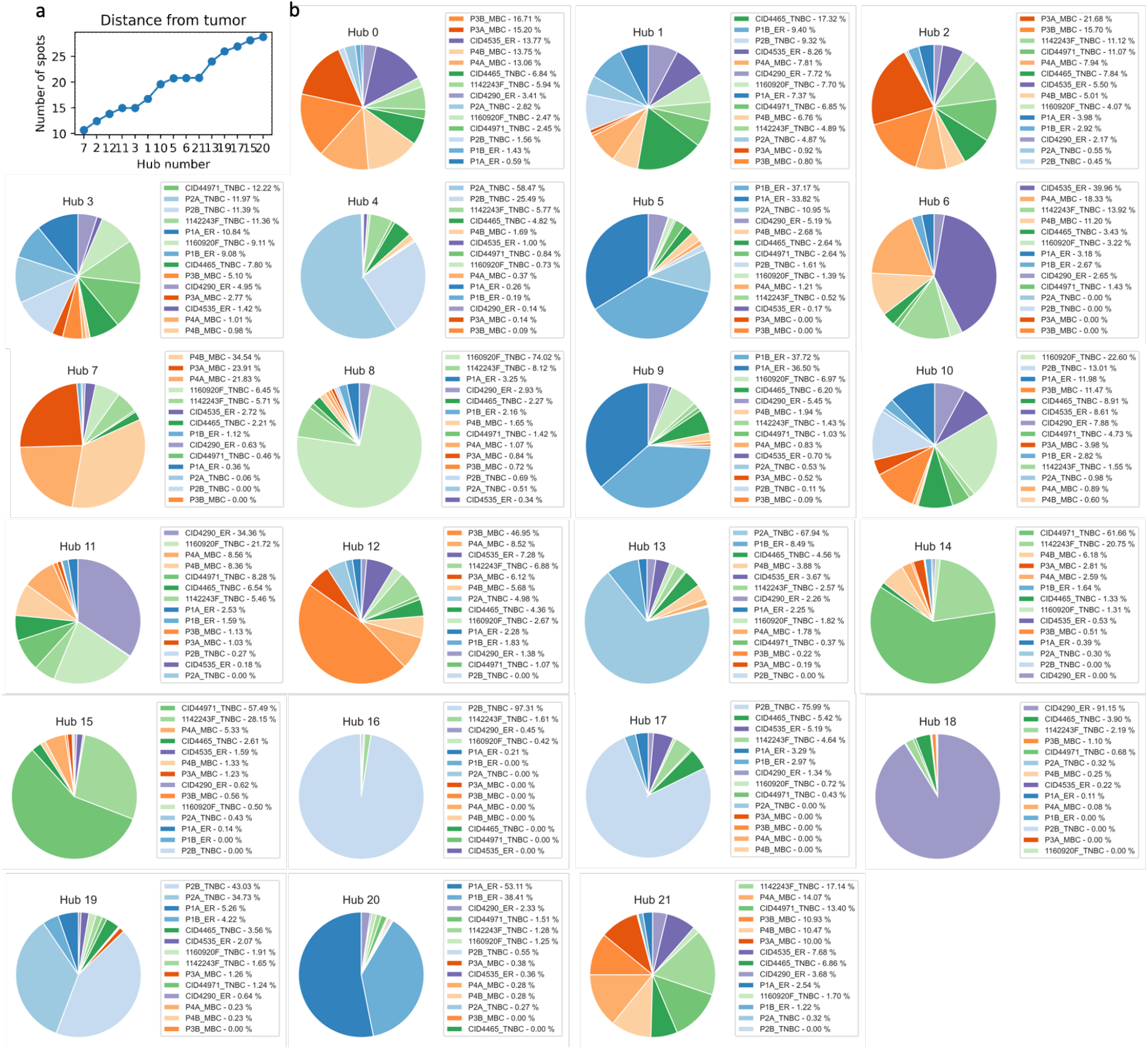
(a) Distance of hubs from tumor center. (b) Pie chart of sample proportions in each hub.

**Supplementary Figure 21.**
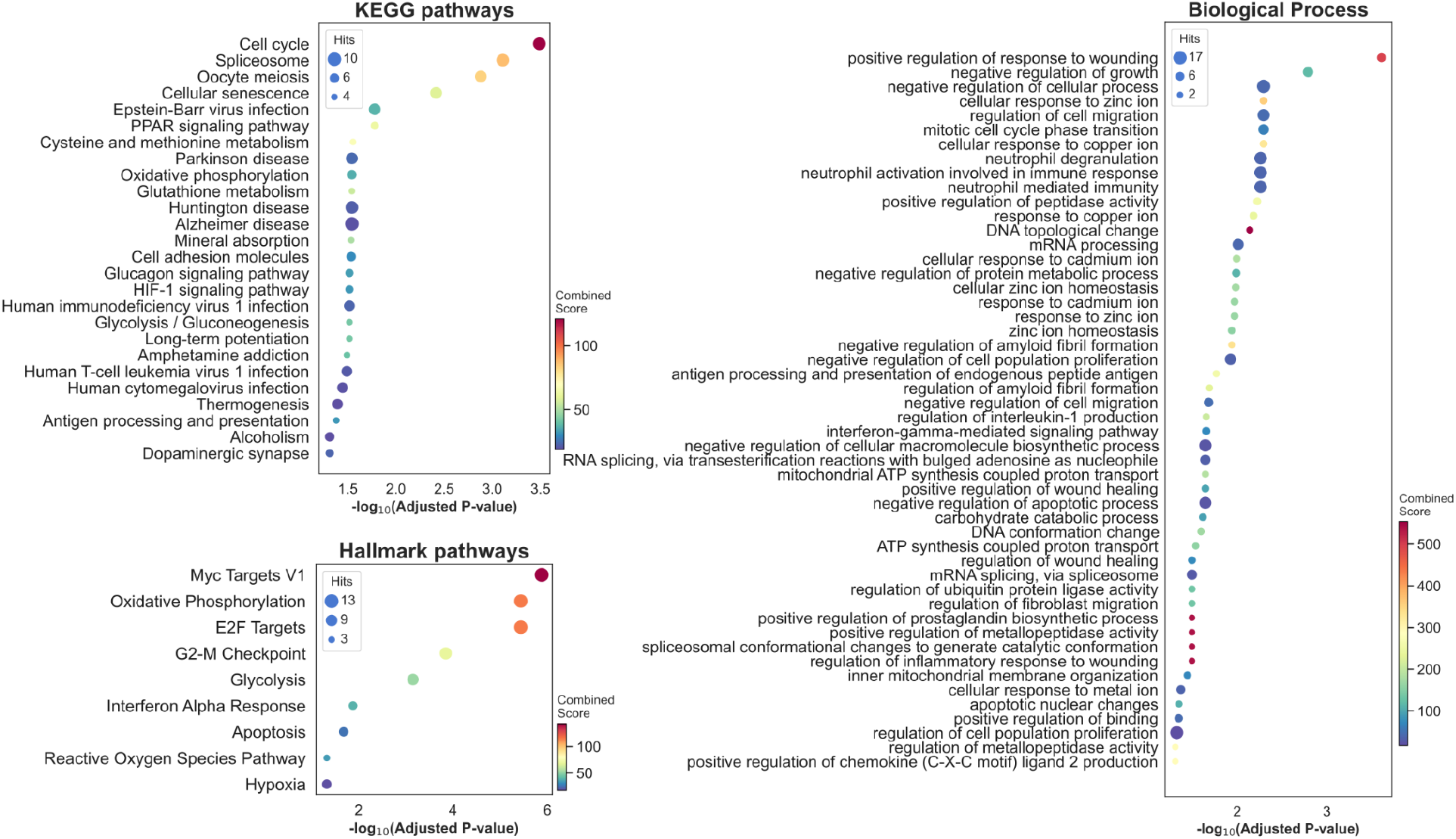
KEGG pathway and biological processes identified for differential genes enriched in TNBC-enriched tumor states.

**Supplementary Figure 22.**
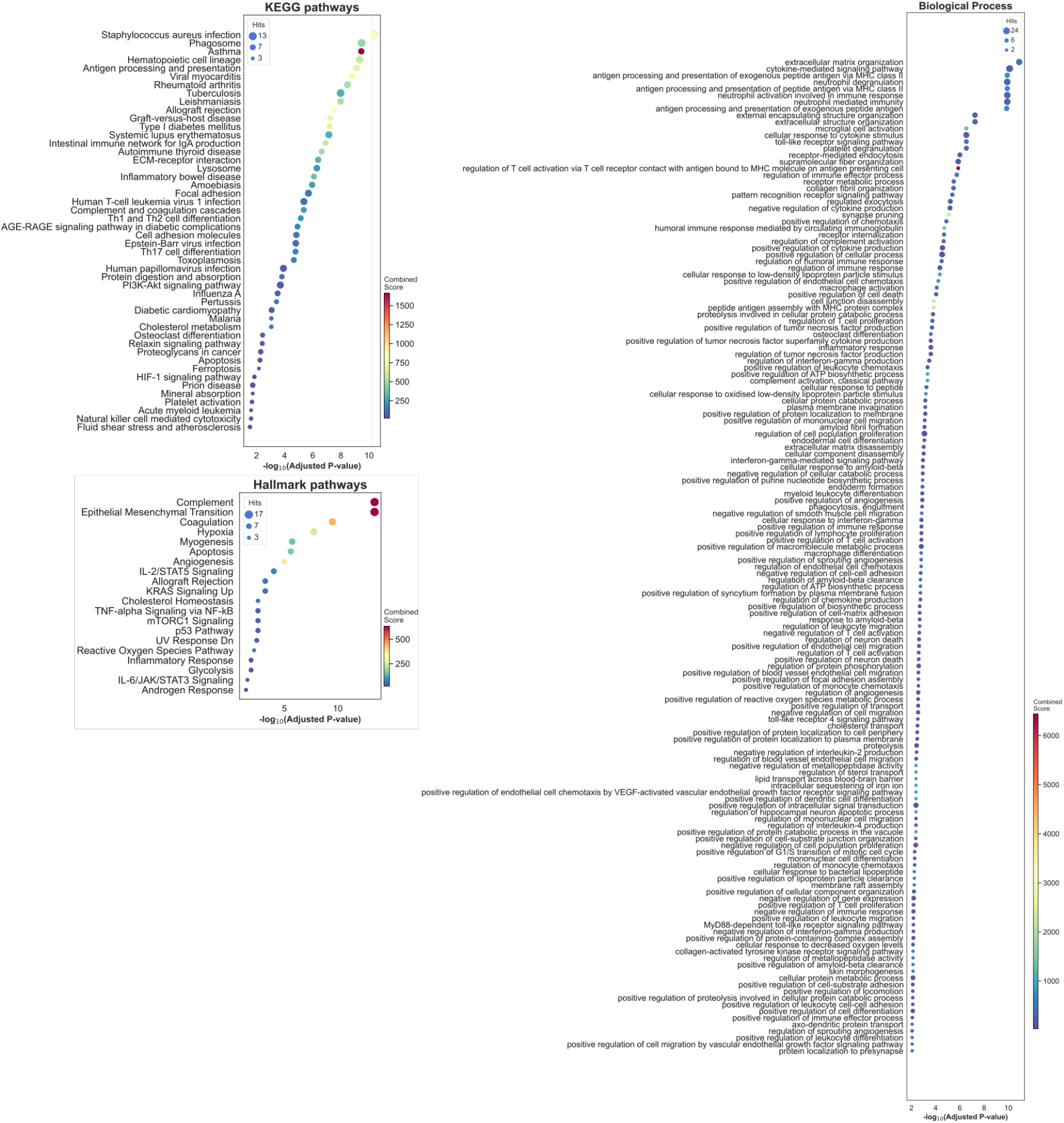
KEGG pathway and biological processes identified for differential genes enriched in MBC-enriched states.

**Supplementary Figure 23.**
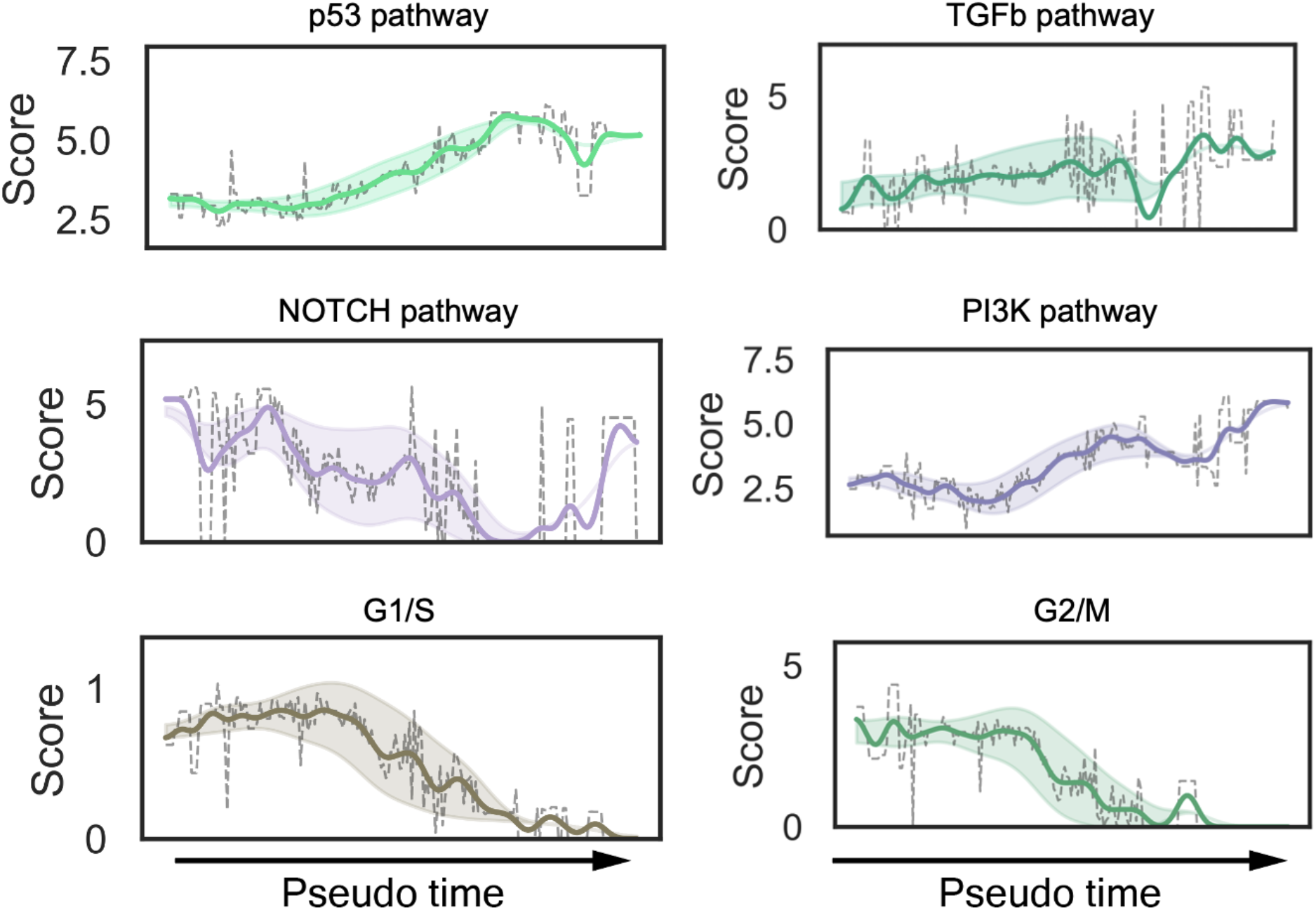
Oncogenic pathways along pseudo-time transition in TAAs across all samples. The score represents the mean expression of genes associated with oncogenic pathways.

**Supplementary Figure 24.**
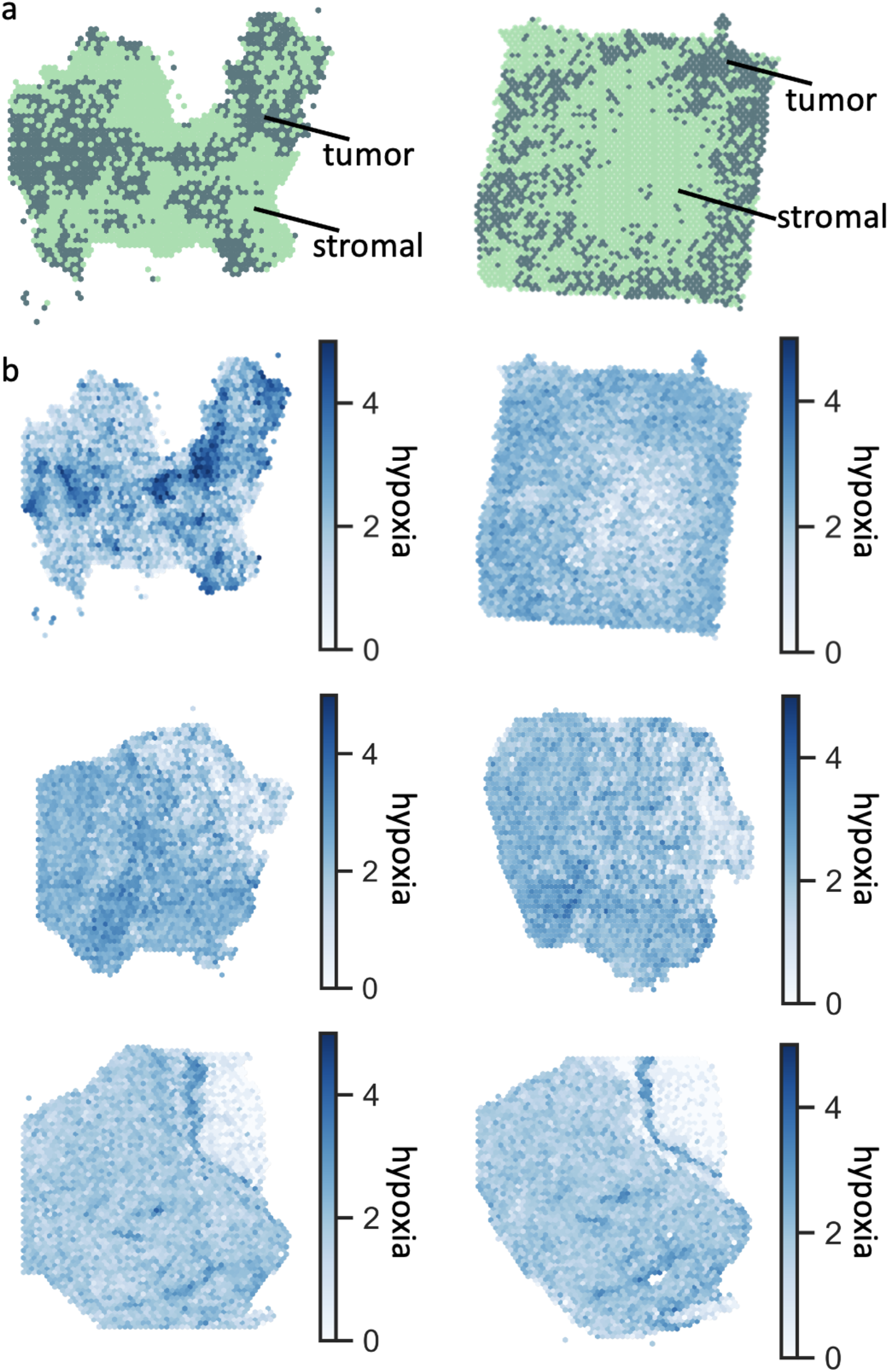
(a) Tumor and stromal regions identified based on histology. (b) Spatial distribution of expression of the hypoxia genesets in MBC, ER, and TNBC samples.

**Supplementary Figure 25.**
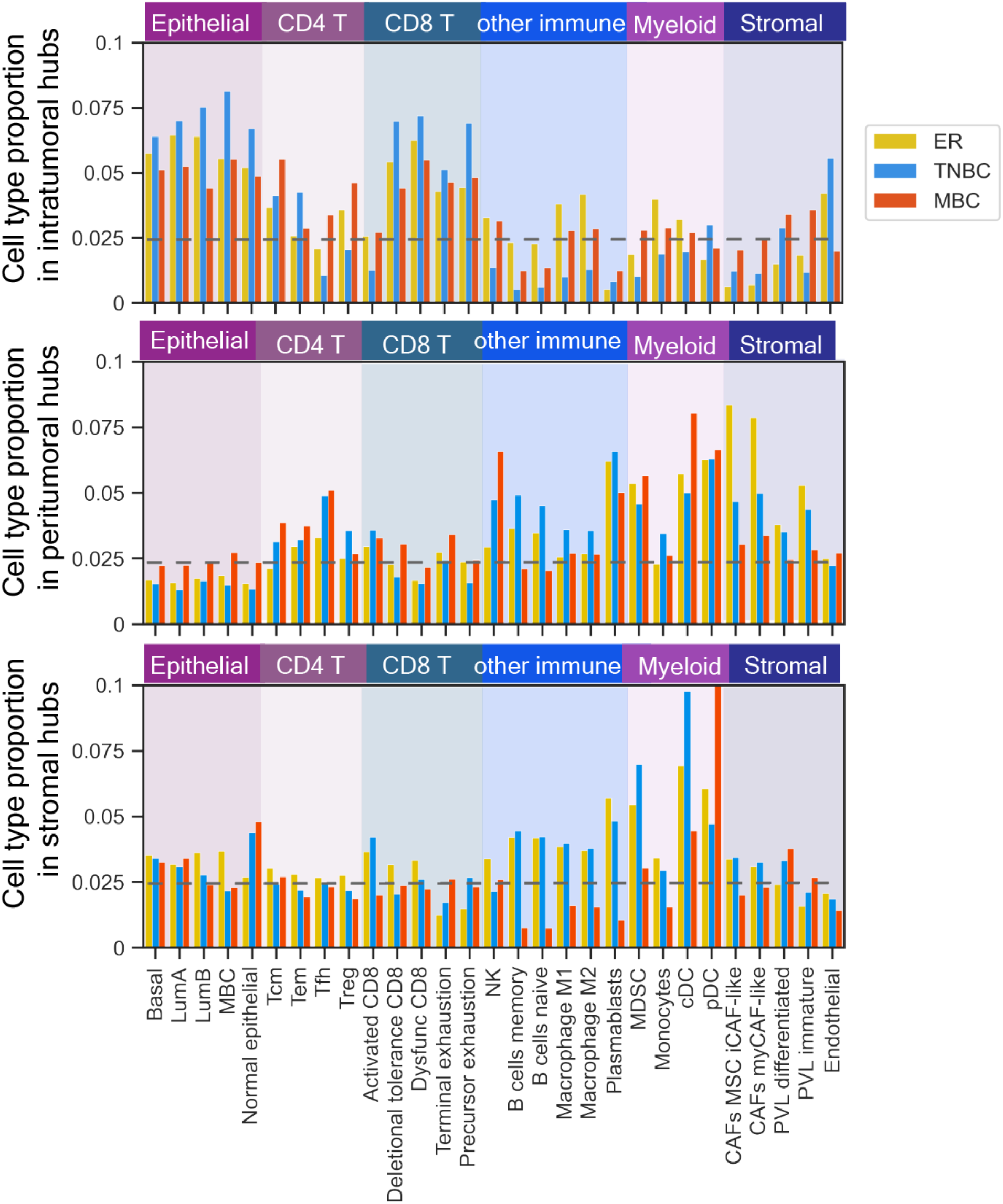
Distribution of cell states in intratumoral, peritumoral, and stromal hubs across all samples. A dashed line indicates a threshold used in determining whether the cell state will be filtered in co-localization analysis.

**Supplementary Figure 26.**
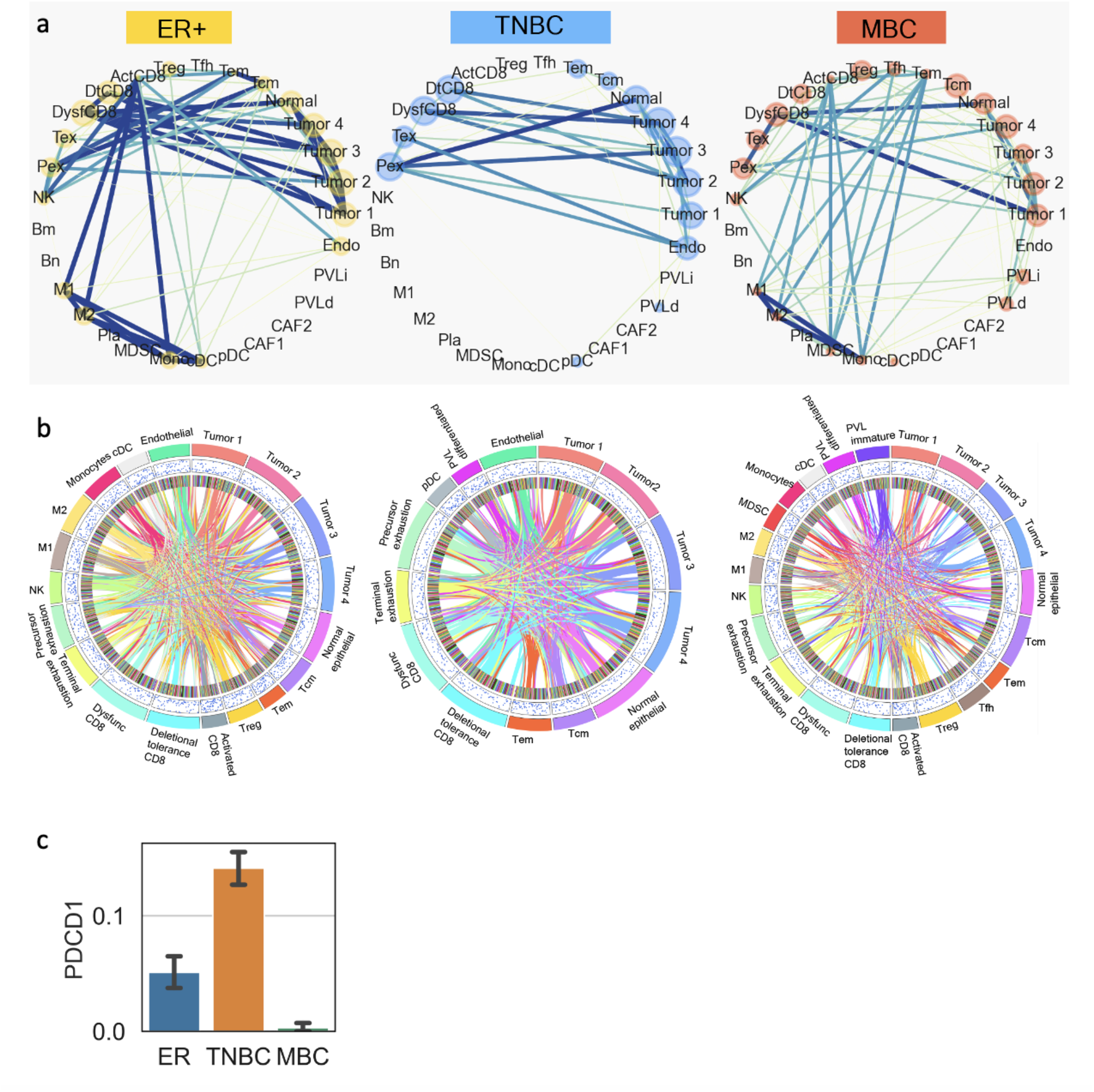
(a) Co-localization score (SCI) for each cell type in intratumoral hubs. (b) Intercellular interactions predicted by CellphoneDB. (c) Expression of PDCD1 in ER, TNBC, and MBC samples.

**Supplementary Figure 27.**
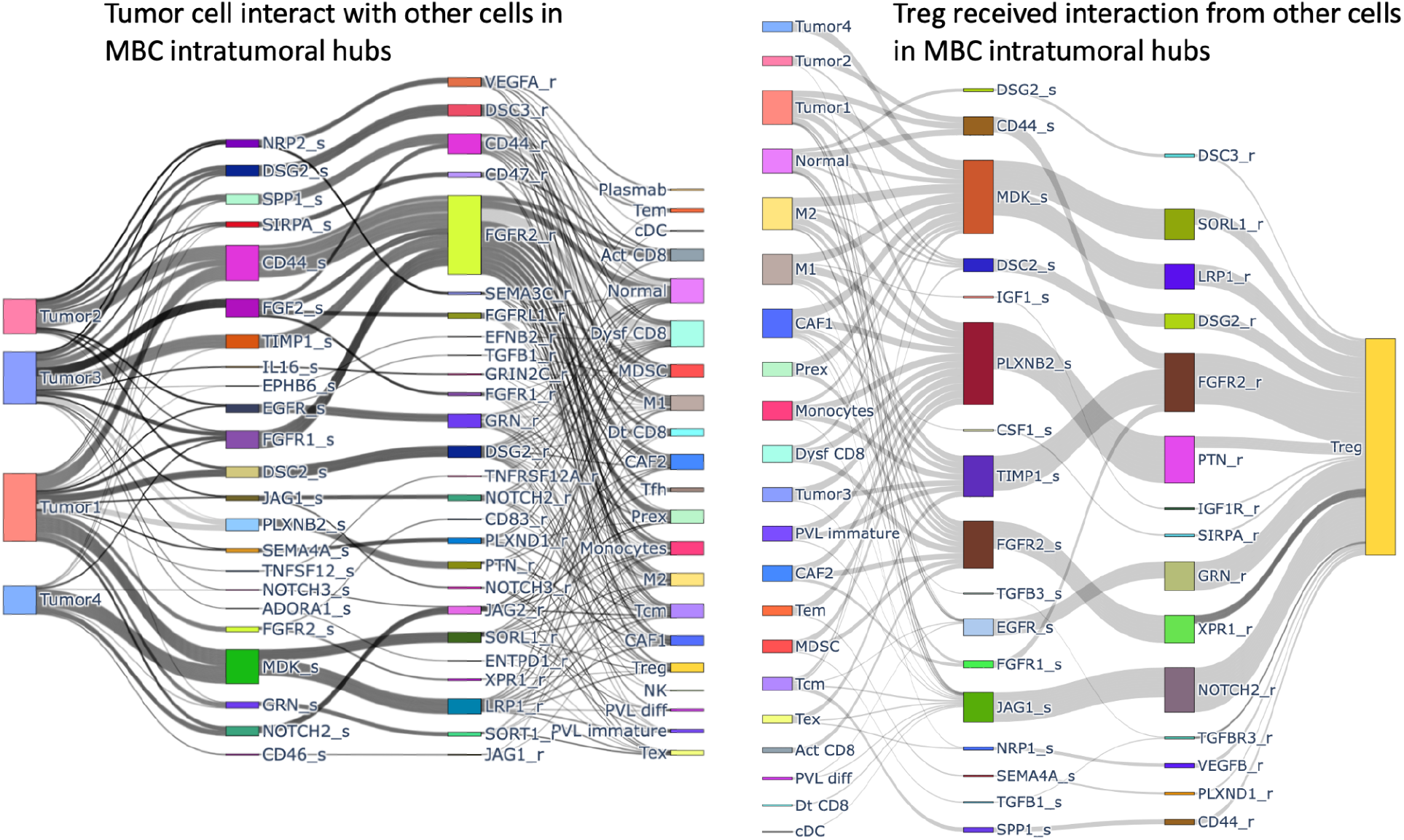
Sankey diagram of ligand-receptor interactions in MBC intratumoral hubs. Left: ligand-receptor interactions predicted between tumor epithelial cells as senders and major immune cells (filtered out states with low proportions) as receivers. Right: ligand-receptor interactions predicted between all other cell states (filtered states with low proportions) as senders and Tregs as receivers.

**Supplementary Figure 28.**
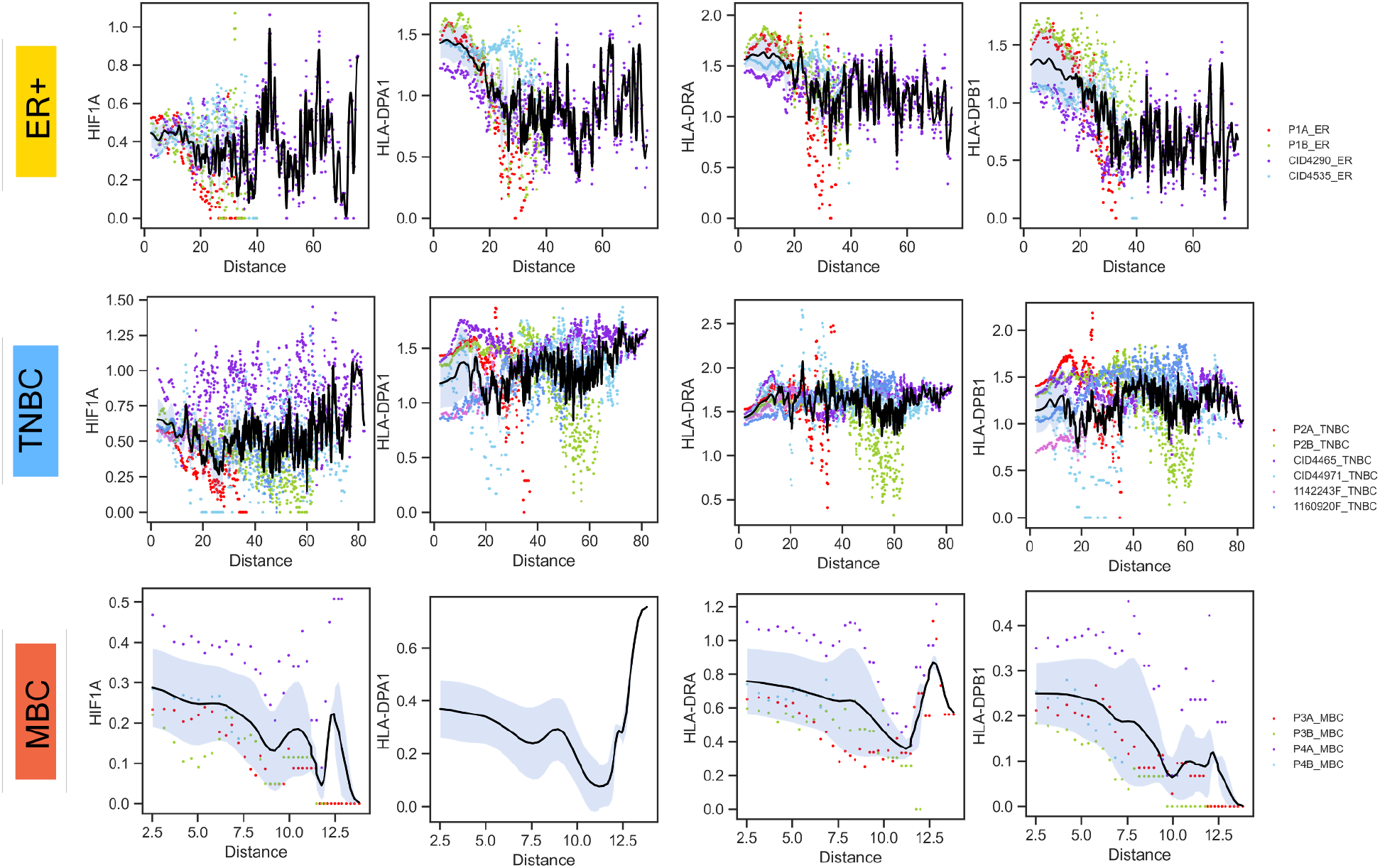
Expression of HIF1, and HLA genes in tumor subtypes vs. distance to Treg-enriched hub 3.

**Supplementary Figure 29.**
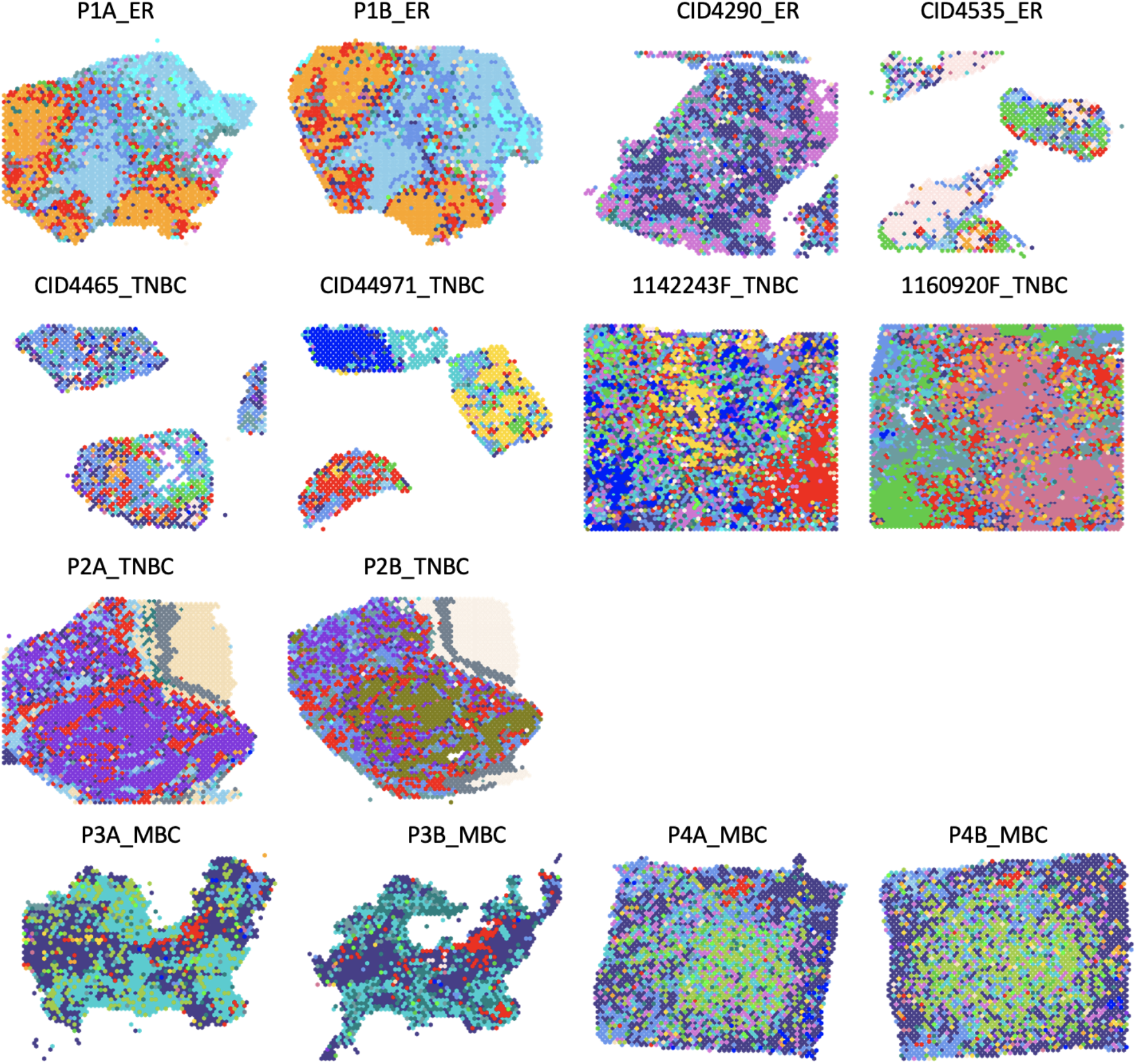
Organization of hubs in all samples studied.

**Supplementary Figure 30.**
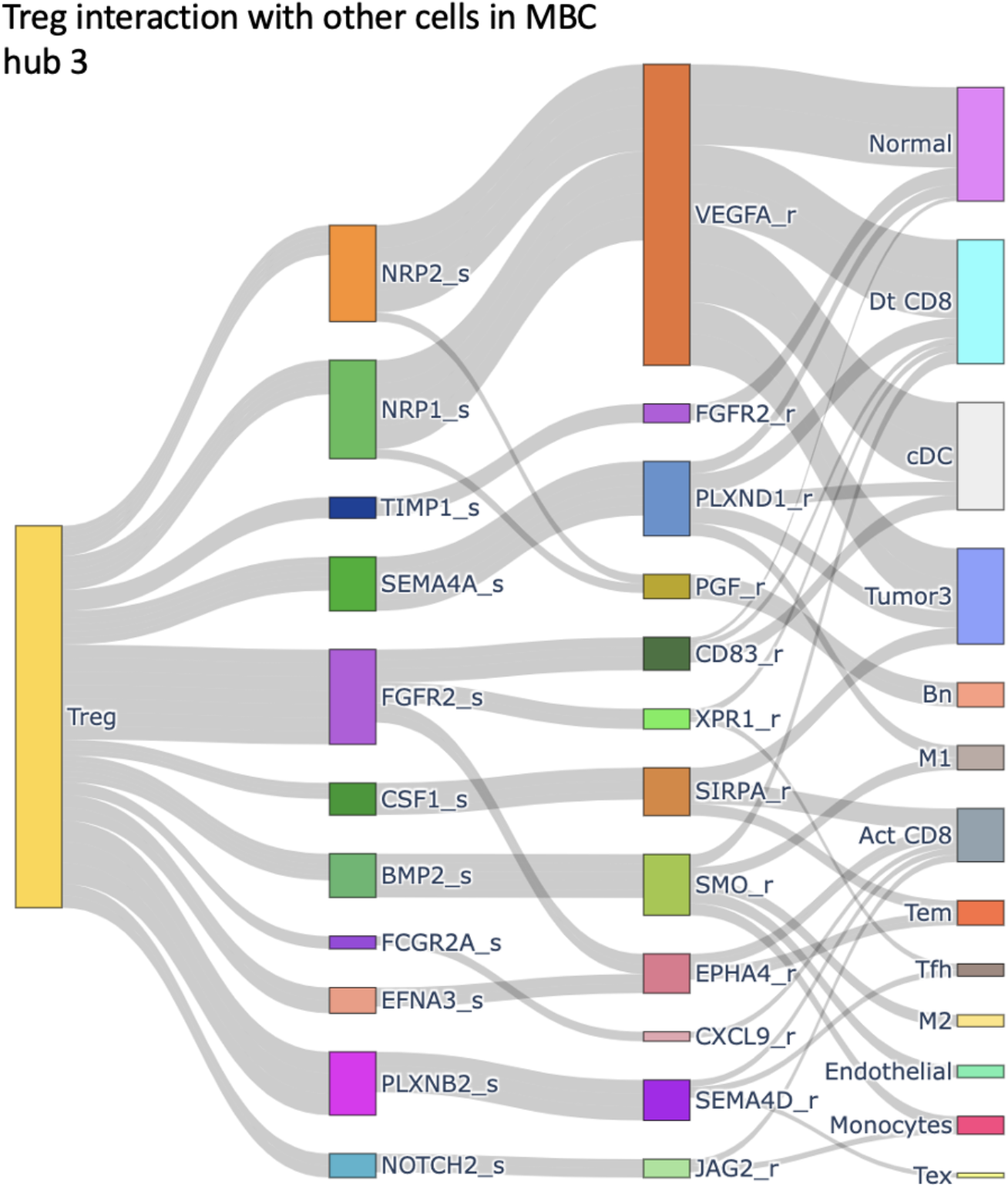
Sankey diagram of predicted ligand-receptor interactions in MBC hub 3 region. Tregs are senders and all other cells (filtering out states with low proportions) as receivers.

## References

1. Armingol, E., Officer, A., Harismendy, O. & Lewis, N. E. Deciphering cell–cell interactions and communication from gene expression. Nature Reviews Genetics vol. 22 71–88 Preprint at https://doi.org/10.1038/s41576-020-00292-x (2021).

2. Ståhl, P. L. et al. Visualization and analysis of gene expression in tissue sections by spatial transcriptomics. Science 353, 78–82 (2016).

3. Chen, W.-T. et al. Spatial Transcriptomics and In Situ Sequencing to Study Alzheimer’s Disease. Cell vol. 182 976–991.e19 Preprint at https://doi.org/10.1016/j.cell.2020.06.038 (2020).

4. Baccin, C. et al. Combined single-cell and spatial transcriptomics reveal the molecular, cellular and spatial bone marrow niche organization. Nature Cell Biology vol. 22 38–48 Preprint at https://doi.org/10.1038/s41556-019-0439-6 (2020).

5. Srivatsan, S. R. et al. Embryo-scale, single-cell spatial transcriptomics. Science 373, 111–117 (2021).

6. Liu, Y. et al. High-Spatial-Resolution Multi-Omics Sequencing via Deterministic Barcoding in Tissue. Cell 183, 1665–1681.e18 (2020).

7. Rodriques, S. G. et al. Slide-seq: A scalable technology for measuring genome-wide expression at high spatial resolution. Science 363, 1463–1467 (2019).

8. Kleshchevnikov, V. et al. Cell2location maps fine-grained cell types in spatial transcriptomics. Nat. Biotechnol. 40, 661–671 (2022).

9. Lopez, R. et al. DestVI identifies continuums of cell types in spatial transcriptomics data. Nat. Biotechnol. (2022) doi:10.1038/s41587-022-01272-8.

10. Biancalani, T. et al. Deep learning and alignment of spatially resolved single-cell transcriptomes with Tangram. Nat. Methods 18, 1352–1362 (2021).

11. Andersson, A. et al. Single-cell and spatial transcriptomics enables probabilistic inference of cell type topography. Commun Biol 3, 565 (2020).

12. Cable, D. M. et al. Robust decomposition of cell type mixtures in spatial transcriptomics. Nat. Biotechnol. 40, 517–526 (2022).

13. Chu, T., Wang, Z., Pe’er, D. & Danko, C. G. Cell type and gene expression deconvolution with BayesPrism enables Bayesian integrative analysis across bulk and single-cell RNA sequencing in oncology. Nat Cancer 3, 505–517 (2022).

14. Miller, B. F., Huang, F., Atta, L., Sahoo, A. & Fan, J. Reference-free cell type deconvolution of multi-cellular pixel-resolution spatially resolved transcriptomics data. Nat. Commun. 13, 2339 (2022).

15. Su, J. et al. A Unified Modular Framework to Incorporate Structural Dependency in Spatial Omics Data. bioRxiv 2022.10.25.513785 (2022) doi:10.1101/2022.10.25.513785.

16. Ma, Y. & Zhou, X. Spatially informed cell-type deconvolution for spatial transcriptomics. Nat. Biotechnol. 1–11 (2022).

17. Bergenstråhle, L. et al. Super-resolved spatial transcriptomics by deep data fusion. Nat. Biotechnol. 40, 476–479 (2022).

18. Zhao, E. et al. Spatial transcriptomics at subspot resolution with BayesSpace. Nat. Biotechnol. 39, 1375–1384 (2021).

19. Wu, S. Z. et al. A single-cell and spatially resolved atlas of human breast cancers. Nat. Genet. 53, (2021).

20. Gayoso, A. et al. A Python library for probabilistic analysis of single-cell omics data. Nat. Biotechnol. 40, 163–166 (2022).

21. Lopez, R., Regier, J., Cole, M. B., Jordan, M. I. & Yosef, N. Deep generative modeling for single-cell transcriptomics. Nat. Methods 15, 1053–1058 (2018).

22. Gayoso, A. et al. Joint probabilistic modeling of single-cell multi-omic data with totalVI. Nat. Methods 18, 272–282 (2021).

23. Lotfollahi, M. et al. Mapping single-cell data to reference atlases by transfer learning. Nat. Biotechnol. 40, 121–130 (2021).

24. Lotfollahi, M., Naghipourfar, M., Theis, F. J. & Wolf, F. A. Conditional out-of-distribution generation for unpaired data using transfer VAE. Bioinformatics 36, i610–i617 (2020).

25. Xu, C. et al. Probabilistic harmonization and annotation of single-cell transcriptomics data with deep generative models. Mol. Syst. Biol. 17, e9620 (2021).

26. Boyeau, P. et al. Deep generative modeling for quantifying sample-level heterogeneity in single-cell omics. bioRxiv 2022.10.04.510898 (2022) doi:10.1101/2022.10.04.510898.

27. Maaløe, L., Sønderby, C. K., Sønderby, S. K. & Winther, O. Auxiliary Deep Generative Models. in International Conference on Machine Learning 1445–1453 (PMLR, 2016).

28. Lee, C. & van der Schaar, M. A Variational Information Bottleneck Approach to Multi-Omics Data Integration. (2021) doi:10.48550/arXiv.2102.03014.

29. He, K., Zhang, X., Ren, S. & Sun, J. Deep residual learning for image recognition. in 2016 IEEE Conference on Computer Vision and Pattern Recognition (CVPR) (IEEE, 2016). doi:10.1109/cvpr.2016.90.

30. Zhang, H. et al. BayesTME: A unified statistical framework for spatial transcriptomics. bioRxiv 2022.07.08.499377 (2022) doi:10.1101/2022.07.08.499377.

31. Szabo, P. A. et al. Single-cell transcriptomics of human T cells reveals tissue and activation signatures in health and disease. Nat. Commun. 10, 1–16 (2019).

32. Vitale, I., Shema, E., Loi, S. & Galluzzi, L. Intratumoral heterogeneity in cancer progression and response to immunotherapy. Nat. Med. 27, 212–224 (2021).

33. Tirosh, I. et al. Dissecting the multicellular ecosystem of metastatic melanoma by single-cell RNA-seq. Science 352, 189–196 (2016).

34. Defining T Cell States Associated with Response to Checkpoint Immunotherapy in Melanoma. Cell 175, 998–1013.e20 (2018).

35. Azizi, E. et al. Single-Cell Map of Diverse Immune Phenotypes in the Breast Tumor Microenvironment. Cell 174, 1293–1308.e36 (2018).

36. Piscuoglio, S. et al. Genomic and transcriptomic heterogeneity in metaplastic carcinomas of the breast. npj Breast Cancer 3, 1–11 (2017).

37. Levine, J. H. et al. Data-Driven Phenotypic Dissection of AML Reveals Progenitor-like Cells that Correlate with Prognosis. Cell 162, 184–197 (2015).

38. Coifman, R. R. et al. Geometric diffusions as a tool for harmonic analysis and structure definition of data: diffusion maps. Proc. Natl. Acad. Sci. U. S. A. 102, 7426–7431 (2005).

39. Haghverdi, L., Buettner, F. & Theis, F. J. Diffusion maps for high-dimensional single-cell analysis of differentiation data. Bioinformatics 31, 2989–2998 (2015).

40. Chen, Z., Wu, J., Wang, L., Zhao, H. & He, J. Tumor-associated macrophages of the M1/M2 phenotype are involved in the regulation of malignant biological behavior of breast cancer cells through the EMT pathway. Medical Oncology vol. 39 Preprint at https://doi.org/10.1007/s12032-022-01670-7 (2022).

41. Reddy, T. P. et al. A comprehensive overview of metaplastic breast cancer: clinical features and molecular aberrations. Breast Cancer Res. 22, 121 (2020).

42. McQuerry, J. A. et al. Pathway activity profiling of growth factor receptor network and stemness pathways differentiates metaplastic breast cancer histological subtypes. BMC Cancer 19, 881 (2019).

43. Djomehri, S. I. et al. Quantitative proteomic landscape of metaplastic breast carcinoma pathological subtypes and their relationship to triple-negative tumors. Nat. Commun. 11, 1–15 (2020).

44. Bachireddy, P. et al. Mapping the evolution of T cell states during response and resistance to adoptive cellular therapy. Cell Rep. 37, 109992 (2021).

45. Morris, E. A. & Liberman, L.. Breast MRI: Diagnosis and Intervention. (Springer Science & Business Media, 2005).

46. Cannoodt, R. et al. SCORPIUS improves trajectory inference and identifies novel modules in dendritic cell development. bioRxiv 079509 (2016) doi:10.1101/079509.

47. Tadros, A. B. et al. Survival Outcomes for Metaplastic Breast Cancer Differ by Histologic Subtype. Ann. Surg. Oncol. 28, 4245–4253 (2021).

48. Moreno, A. C. et al. Outcomes after Treatment of Metaplastic Versus Other Breast Cancer Subtypes. J. Cancer 11, 1341 (2020).

49. Wong, W. et al. Poor response to neoadjuvant chemotherapy in metaplastic breast carcinoma. npj Breast Cancer 7, 1–7 (2021).

50. Schwartz, T. L., Mogal, H., Papageorgiou, C., Veerapong, J. & Hsueh, E. C. Metaplastic breast cancer: histologic characteristics, prognostic factors and systemic treatment strategies. Exp. Hematol. Oncol. 2, 31 (2013).

51. Kalaw, E. et al. Metaplastic breast cancers frequently express immune checkpoint markers FOXP3 and PD-L1. Br. J. Cancer 123, 1665–1672 (2020).

52. Hapke, R. Y. & Haake, S. M. Hypoxia-induced epithelial to mesenchymal transition in cancer. Cancer Lett. 487, 10–20 (2020).

53. Romeo, E., Caserta, C. A., Rumio, C. & Marcucci, F. The Vicious Cross-Talk between Tumor Cells with an EMT Phenotype and Cells of the Immune System. Cells 8, (2019).

54. Ye, L.-Y. et al. Hypoxia-Induced Epithelial-to-Mesenchymal Transition in Hepatocellular Carcinoma Induces an Immunosuppressive Tumor Microenvironment to Promote Metastasis. Cancer Res. 76, 818–830 (2016).

55. Muz, B., de la Puente, P., Azab, F. & Azab, A. K. The role of hypoxia in cancer progression, angiogenesis, metastasis, and resistance to therapy. Hypoxia (Auckl) 3, 83–92 (2015).

56. Efremova, M., Vento-Tormo, M., Teichmann, S. A. & Vento-Tormo, R. CellPhoneDB: inferring cell–cell communication from combined expression of multi-subunit ligand–receptor complexes. Nat. Protoc. 15, 1484–1506 (2020).

57. Shu, C. et al. Virus-Like Particles Presenting the FGF-2 Protein or Identified Antigenic Peptides Promoted Antitumor Immune Responses in Mice. Int. J. Nanomedicine 15, 1983 (2020).

58. Liu, L. et al. Sp1 induced gene TIMP1 is related to immune cell infiltration in glioblastoma. Sci. Rep. 12, 1–18 (2022).

59. D’Costa, Z. et al. Gemcitabine-induced TIMP1 attenuates therapy response and promotes tumor growth and liver metastasis in pancreatic cancer. Cancer Res. 77, 5952–5962 (2017).

60. Palakurthi, S. et al. The Combined Effect of FGFR Inhibition and PD-1 Blockade Promotes Tumor-Intrinsic Induction of Antitumor Immunity. Cancer Immunol Res 7, 1457–1471 (2019).

61. Bollyky, P. L. et al. CD44 costimulation promotes FoxP3+ regulatory T cell persistence and function via production of IL-2, IL-10, and TGF-beta. J. Immunol. 183, 2232–2241 (2009).

62. Miller, B. F., Bambah-Mukku, D., Dulac, C., Zhuang, X. & Fan, J. Characterizing spatial gene expression heterogeneity in spatially resolved single-cell transcriptomic data with nonuniform cellular densities. Genome Res. 31, 1843–1855 (2021).

63. da Silva, E. M. et al. TERT promoter hotspot mutations and gene amplification in metaplastic breast cancer. NPJ Breast Cancer 7, 43 (2021).

64. Pareja, F. et al. The genomic landscape of metastatic histologic special types of invasive breast cancer. npj Breast Cancer 6, 1–10 (2020).

65. Shin, E. & Koo, J. S. Glucose Metabolism and Glucose Transporters in Breast Cancer. Front Cell Dev Biol 9, 728759 (2021).

66. Lien, E. C. et al. Glutathione biosynthesis is a metabolic vulnerability in PI(3)K/Akt-driven breast cancer. Nat. Cell Biol. 18, 572–578 (2016).

67. Dumond, A. & Pagès, G. Neuropilins, as Relevant Oncology Target: Their Role in the Tumoral Microenvironment. Front Cell Dev Biol 8, 662 (2020).

68. Parish, I. A. et al. The molecular signature of CD8+ T cells undergoing deletional tolerance. Blood 113, 4575–4585 (2009).

69. Klement, J. D. et al. An osteopontin/CD44 immune checkpoint controls CD8+ T cell activation and tumor immune evasion. J. Clin. Invest. 128, 5549–5560 (2018).

70. Denhardt, D. T., Noda, M., O’Regan, A. W., Pavlin, D. & Berman, J. S. Osteopontin as a means to cope with environmental insults: regulation of inflammation, tissue remodeling, and cell survival. J. Clin. Invest. 107, 1055–1061 (2001).

71. Weng, X., Maxwell-Warburton, S., Hasib, A., Ma, L. & Kang, L. The membrane receptor CD44: novel insights into metabolism. Trends Endocrinol. Metab. 33, 318–332 (2022).

72. Holtzhausen, A. et al. TAM Family Receptor Kinase Inhibition Reverses MDSC-Mediated Suppression and Augments Anti–PD-1 Therapy in Melanoma. Cancer Immunol Res 7, 1672–1686 (2019).

73. Tanaka, M. & Siemann, D. W. Gas6/Axl Signaling Pathway in the Tumor Immune Microenvironment. Cancers 12, 1850 (2020).

74. Brown, W. S., Akhand, S. S. & Wendt, M. K. FGFR signaling maintains a drug persistent cell population following epithelial-mesenchymal transition. Oncotarget 7, 83424–83436 (2016).

75. Perez-Garcia, J., Muñoz-Couselo, E., Soberino, J., Racca, F. & Cortes, J. Targeting FGFR pathway in breast cancer. Breast 37, 126–133 (2018).

76. Abdel-Wahab, N. et al. Checkpoint inhibitor therapy for cancer in solid organ transplantation recipients: an institutional experience and a systematic review of the literature. J Immunother Cancer 7, 106 (2019).

77. Stickels, R. R. et al. Highly sensitive spatial transcriptomics at near-cellular resolution with Slide-seqV2. Nat. Biotechnol. 39, 313–319 (2021).

78. Wang, Y. et al. Multi-modal single-cell and whole-genome sequencing of small, frozen clinical specimens. Nature Genetics, In Press.

79. Cutler, A. & Breiman, L. Archetypal Analysis. Technometrics vol. 36 338–347 Preprint at https://doi.org/10.1080/00401706.1994.10485840 (1994).

80. van Dijk, D. et al. Recovering Gene Interactions from Single-Cell Data Using Data Diffusion. Cell 174, 716–729.e27 (2018).

81. Mohammadi, S., Ravindra, V., Gleich, D. F. & Grama, A. A geometric approach to characterize the functional identity of single cells. Nat. Commun. 9, 1516 (2018).

82. Wang, Y. & Zhao, H. Non-linear archetypal analysis of single-cell RNA-seq data by deep autoencoders. PLoS Comput. Biol. 18, e1010025 (2022).

83. Mørup, M. & Hansen, L. K. Archetypal analysis for machine learning and data mining. Neurocomputing vol. 80 54–63 Preprint at https://doi.org/10.1016/j.neucom.2011.06.033 (2012).

84. Hastie, T., Tibshirani, R. & Friedman, J. The Elements of Statistical Learning. Springer Series in Statistics Preprint at https://doi.org/10.1007/978-0-387-84858-7 (2009).

85. Albergante, L., Bac, J. & Zinovyev, A. Estimating the effective dimension of large biological datasets using Fisher separability analysis. 2019 International Joint Conference on Neural Networks (IJCNN) Preprint at https://doi.org/10.1109/ijcnn.2019.8852450 (2019).

86. Kuchroo, M. et al. Multiscale PHATE identifies multimodal signatures of COVID-19. Nat. Biotechnol. 40, 681–691 (2022).

87. Wolf, F. A., Angerer, P. & Theis, F. J. SCANPY: large-scale single-cell gene expression data analysis. Genome Biol. 19, 15 (2018).

88. Blei, D. M., Kucukelbir, A. & McAuliffe, J. D. Variational inference: A review for statisticians. J. Am. Stat. Assoc. 112, 859–877 (2017).

89. Paszke, A. et al. PyTorch: An Imperative Style, High-Performance Deep Learning Library. Adv. Neural Inf. Process. Syst. 32, (2019).

90. Kingma, D. P. & Ba, J. Adam: A Method for Stochastic Optimization. (2014) doi:10.48550/arXiv.1412.6980.

91. Hideto, M., Base, M., Bullock, D., bw & snajder-r. ponnhide/pyCircos: pyCircos: Circos plot in matplotlib. (2022) doi:10.5281/zenodo.6477641.

